# GLS-driven glutamine catabolism contributes to prostate cancer radiosensitivity by regulating the redox state, stemness and ATG5-mediated autophagy

**DOI:** 10.1101/2021.01.07.425771

**Authors:** Anna Mukha, Uğur Kahya, Annett Linge, Oleg Chen, Steffen Löck, Vasyl Lukiyanchuk, Susan Richter, Tiago C. Alves, Mirko Peitzsch, Vladyslav Telychko, Sergej Skvortsov, Giulia Negro, Bertram Aschenbrenner, Ira-Ida Skvortsova, Peter Mirtschink, Fabian Lohaus, Tobias Hölscher, Hans Neubauer, Mahdi Rivandi, André Franken, Bianca Behrens, Nikolas H. Stoecklein, Marieta Toma, Ulrich Sommer, Sebastian Zschaeck, Maximilian Rehm, Graeme Eisenhofer, Christian Schwager, Amir Abdollahi, Christer Groeben, Leoni A. Kunz-Schughart, Gustavo B. Baretton, Michael Baumann, Mechthild Krause, Claudia Peitzsch, Anna Dubrovska

**Affiliations:** OncoRay - National Center for Radiation Research in Oncology, Faculty of Medicine and University Hospital Carl Gustav Carus, Technische Universität Dresden and Helmholtz-Zentrum Dresden-Rossendorf, Germany; Institute of Radiooncology – OncoRay, Helmholtz-Zentrum Dresden-Rossendorf (HZDR) Dresden, Germany; German Cancer Consortium (DKTK), Partner Site Dresden, Germany; German Cancer Research Center (DKFZ), Heidelberg, Germany; Department of Radiotherapy and Radiation Oncology, Faculty of Medicine and University Hospital Carl Gustav Carus, Technische Universität Dresden, Germany; National Center for Tumor Diseases (NCT) Partner Site Dresden, Germany; Department of Cell Signaling, Institute of Cell Biology, NAS of Ukraine, Lviv, Ukraine; Institute of Clinical Chemistry and Laboratory Medicine, University Hospital Carl Gustav Carus, Technische Universität Dresden, Germany; Department for Clinical Pathobiochemistry, Faculty of Medicine and University Hospital Carl Gustav Carus, Technische Universität Dresden, Germany; Department of Therapeutic Radiology and Oncology, Medical University of Innsbruck, Innsbruck, Austria; EXTRO-Lab, Tyrolean Cancer Research Institute, Innsbruck, Austria; Department of Obstetrics and Gynecology, Medical Faculty and University Hospital of the Heinrich-Heine University Düsseldorf, Germany; General, Visceral and Paediatric Surgery, University Hospital and Medical Faculty of the Heinrich-Heine University Düsseldorf, Düsseldorf, Germany; Institute of Pathology, University of Bonn, Bonn, Germany; Institute of Pathology, Universitätsklinikum Carl Gustav Carus Dresden, Dresden, Germany; Heidelberg Ion-Beam Therapy Center (HIT), Department of Radiation Oncology, Heidelberg University Hospital (UKHD), National Center for Tumor Diseases (NCT), Heidelberg, Germany; German Cancer Consortium (DKTK) Core Center, Clinical Cooperation Units (CCU) Translational Radiation Oncology and Radiation Oncology, Heidelberg, Germany; Heidelberg Institute of Radiation Oncology (HIRO), National Center for Radiation Research in Oncology (NCRO), German Cancer Research Center (DKFZ) and Heidelberg University Hospital (UKHD), Heidelberg, Germany; Division of Molecular and Translational Radiation Oncology, Heidelberg Medical Faculty (HDMF), Heidelberg University, Heidelberg, Germany; Department of Urology, Medical Faculty Carl Gustav Carus, TU Dresden, Dresden, Germany

**Keywords:** Prostate cancer, Radioresistance, Glutamine metabolism, Cancer stem cells, Autophagy, MYC, GLS1

## Abstract

Radiotherapy is one of the curative treatment options for localized prostate cancer (PCa). The curative potential of radiotherapy is mediated by irradiation-induced oxidative stress and DNA damage in tumor cells. However, PCa radiocurability can be impeded by tumor resistance mechanisms and normal tissue toxicity. Metabolic reprogramming is one of the major hallmarks of tumor progression and therapy resistance. Here, we found that radioresistant PCa cells and prostate cancer stem cells (CSCs) have a high glutamine demand. Glutaminase (GLS)-driven catabolism of glutamine serves not only for energy production but also for the maintenance of the redox state. Consequently, glutamine depletion or inhibition of critical regulators of glutamine utilization, such as glutaminase (GLS) and the transcription factor MYC results in PCa radiosensitization. On the contrary, we found that a combination of glutamine metabolism inhibitors with irradiation does not cause toxic effects on nonmalignant prostate cells. Glutamine catabolism contributes to the maintenance of CSCs through regulation of the alpha-ketoglutarate (α-KG)-dependent chromatin-modifying dioxygenase. The lack of glutamine results in the inhibition of CSCs with a high aldehyde dehydrogenase (ALDH) activity, decreases the frequency of the CSC populations *in vivo* and reduces tumor formation in xenograft mouse models. Moreover, this study shows that activation of the ATG5-mediated autophagy in response to a lack of glutamine is a tumor survival strategy to withstand radiation-mediated cell damage. In combination with autophagy inhibition, the blockade of glutamine metabolism might be a promising strategy for PCa radiosensitization. High blood levels of glutamine in PCa patients significantly correlate with a shorter prostate-specific antigen (PSA) doubling time. Furthermore, high expression of critical regulators of glutamine metabolism, GLS1 and MYC, is significantly associated with a decreased progression-free survival in PCa patients treated with radiotherapy. Our findings demonstrate that GLS-driven glutaminolysis is a prognostic biomarker and therapeutic target for PCa radiosensitization.

## Introduction

Radiotherapy is one of the mainstay curative treatments for clinically localized prostate cancer (PCa). Most PCa patients with low-risk disease can be cured with surgery (prostatectomy) or radiotherapy with five-year progression-free survival rates over 90%, whereas for high-risk PCa patients outcome remains poor ^1^. The curative potential of radiotherapy depends on the capability of tumor cells to deal with irradiation-induced oxidative stress and DNA damage. Treatment failure is often attributed to the tumor resistance mechanisms and normal tissue toxicity ^2, 3^. The current risk stratification system for PCa based on both clinical and pathologic factors is associated with substantional heterogeneity in outcomes, hence, additional biological markers that predict tumor radiosensitivity are required ^1^.

Reprogramming of cellular metabolism plays an essential role in tumor initiation, progression, and therapy resistance ^4^. Specific biochemical and molecular metabolic features of PCa tissues might serve as biomarkers for identifying the patients most likely to respond to radiotherapy. To meet high energetic demands, fast-growing tumor cells often reprogram their metabolic pathways and exhibit a high level of nutrient consumption. Cancer cells fuel their growth and proliferation through the metabolism of two primary substrates, glucose and glutamine; the latter is an essential donor of nitrogen and carbon for growth-promoting pathways. Although most normal tissues can synthesize glutamine, it becomes conditionally essential amino acid for fast-growing tumors ^5-8^. High demand for glutamine is particularly true for tumors that acquire oncogene-driven glutamine addiction, including PCa with high MYC expression ^5, 9, 10^. Glutamine contributes to production of energy and building blocks in cancer cells by feeding into the tricarboxylic acid (TCA) cycle with subsequent ATP generation through mitochondrial oxidative phosphorylation (OXPHOS) or recycling of reducing equivalents by lactate secretion (glutaminolysis) ^11-13^. Glutamine also plays an essential role in the epigenetic reprogramming as glutaminolysis converts it to alpha-ketoglutarate (α-KG), a critical cofactor for the Jumonji-domain-containing histone demethylases and Fe(II)/α-KG dioxygenase that regulate DNA demethylation and repair ^14, 15^. Glutamine is also involved in maintaining homeostasis of reactive oxygen species (ROS) as an amino acid component of the glutathione (GSH) system and is vital for nucleotide and amino acid biosynthesis ^8, 11, 16^.

Similar to many other cancers, PCa has a hierarchical organization where cancer stem cell (CSC) populations maintain and propagate the tumor mass ^10^. Although CSCs and their non-tumorigenic non-CSC progenies within the same tumor clone share a common genotype, they display different epigenetic profiles associated with specific metabolic programs and signaling networks ^17^. Many of these signaling pathways confer cell adaptation to microenvironmental stresses, including a shortage of nutrients and anti-cancer therapies ^10, 17-19^. A body of evidence suggests increased radioresistance of CSCs compared to the tumor bulk. In support of this, recent clinical studies used CSC related biomarkers for prediction of radiotherapy outcome ^20, 21^. Nevertheless, the role of glutamine metabolism in the maintenance of PCa CSCs and radioresistance remains unclear and is the focus of this study.

Here, we report that PCa cells with acquired radioresistance rewire their metabolism and become more glutamine dependent. In these cells, glutamine plays an important role not only in fueling biosynthesis and energy production pathways, but also supports GSH synthesis and the maintenance of the redox state.Consequently, glutamine depletion or inhibition of the critical regulators of glutamine utilization, such as glutaminase (GLS), the enzyme converting glutamine to glutamate, and the transcription factor MYC results in PCa radiosensitization. On the contrary, the combination of this inhibition with irradiation is less toxic for nonmalignant prostate cells. In response to Gln deprivation, PCa cells activate ATG5-mediated autophagy as a survival mechanism to overcome nutrient stress, suggesting that a combination of glutamine metabolism blockade and autophagy inhibition might be a promising strategy for PCa radiosensitization.

Similar to radioresistant PCa cells, prostate CSCs have a high glutamine demand. Glutamine catabolism contributes to the maintenance of CSCs by epigenetic resetting through regulation of the chromatin-modifying dioxygenase. Inhibition of glutamine metabolism results in depleting the CSC population confirmed by *in vivo* limiting dilution assay. Consequently, glutamine deprivation in combination with irradiation resulted in a loss of tumor-forming capacity in glutamine-addicted PCa cells.

In PCa patients treated with radiotherapy, GLS and MYC expression levels are significantly associated with decreased relapse-free survival. In agreement with this observation, we found that elevated blood level of glutamine is significantly associated with a short prostate specific antigen (PSA) doubling time. Together, our results indicate that GLS-driven glutaminolysis is a prognostic biomarker and a promising therapeutic target for PCa radiosensitization.

## Results

### Gln metabolism is dysregulated in radioresistant PCa cells

To identify the metabolic traits associated with radiation resistance, we analyzed radioresistant isogenic sublines (RR) of the conventional PCa cell lines DU145, LNCaP and PC3, as described earlier ^22, 23^. These RR cells exhibit more efficient DNA repair, possess fast *in vivo* tumor growth, and have an enriched cancer stem cell (CSC) phenotype, including high aldehyde dehydrogenase (ALDH) activity ^22, 23^. Global gene expression profiling demonstrated that radioresistant properties of LNCaP RR and DU145 RR cells are associated with a broad metabolic reprogramming, including deregulation of genes involved in the control of tricarboxylic acid (TCA) cycle and amino acid metabolism (**Figure 1 a and Supplementary Figure S 1 a**). This reprogramming driven by acquired radioresistance was more pronounced in DU145 as compared to LNCaP cells (**Figure 1 b and Supplementary Figure S 1 b**), although gene expression deregulation had a weak correlation between two cell lines. Targeted metabolomics of amino acids and biogenic amines in parental and RR cells revealed that intracellular levels of glutamate (Glu) are significantly upregulated in both PC3 RR and DU145 RR cells, but not in LNCaP RR cells as compared to the corresponding parental cells (**Figure 1 c**). With a focus on amino acids and biogenic amines, we performed principal component analysis (PCA) to visualize metabolic differences and the relationship of the analyzed parental and RR cells. We observed that metabolic differences between cell lines are stronger than within each parental / RR pair. These differences are driven in particular by altered levels of the glucogenic amino acids, including Glu and glutamine (Gln) (**Supplementary Figure S 1 c**).

**Figure 1.**
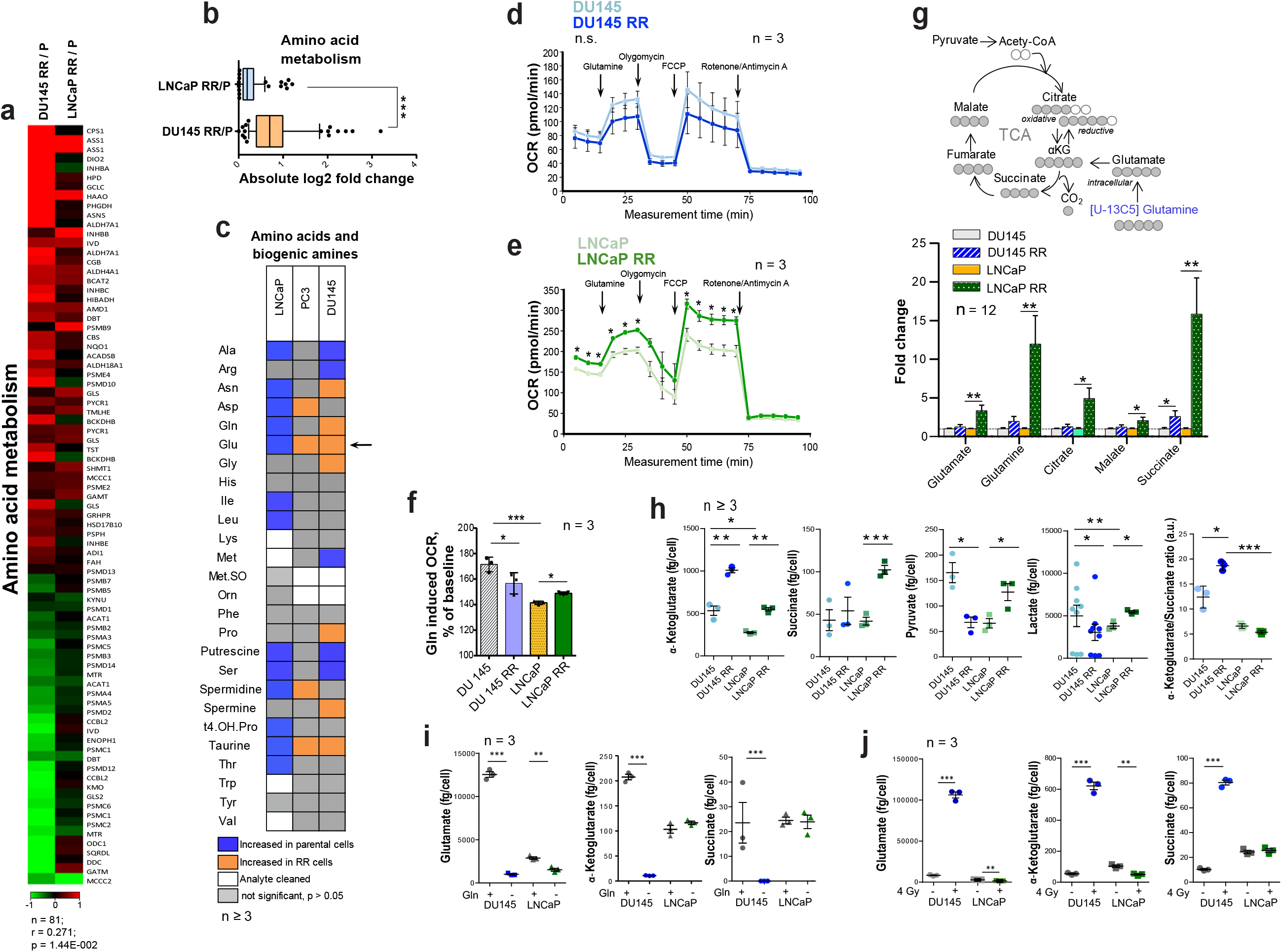
Deregulation of Gln metabolism in radioresistant PCa cells. **a** Gene expression analysis showed that radioresistant properties of LNCaP RR and DU145 RR cells are associated with similar deregulation pattern of genes involved in the control of amino acid metabolism. **b** Absolute log2 fold change of genes included in amino acid metabolism signature. **c** Targeted metabolomics of amino acids and biogenic amines in parental and RR cells revealed that Glu is significantly upregulated in both PC3 RR and DU145 RR cells, but not in LNCaP RR cells as compared to the corresponding parental cells. **d, e** Agilent Seahorse XF analysis of Gln-starved parental and RR DU145 and LNCaP cells, accordingly. Real-time oxygen consumption rate (OCR) was measured after the addition of Gln to the final concentration of 2 mM. Data are mean ± s.e.m. **f** Increase in OCR over the baseline after Gln supplementation. Data are mean ± s.d. **g** LC-MS/MS based analysis of the contribution of ^13^C_5_-Gln to TCA cycle intermediates based on isotopic steady state labeling. ^13^C_5_-Gln was supplemented to the culture medium at concentration of 2 mM, and cells were harvested 4 h later. **h** The intracellular levels of TCA cycle metabolites measured in parental and RR cells by LC-MS/MS. Data are mean ± s.e.m. **i** Quantification of intracellular metabolites in control cells and cells deprived for Gln for 24 h. Data are mean ± s.e.m. **j** Intracellular levels of Gln, α-KG, and succinate in control and irradiated cells. Cells were sham-irradiated or irradiated with 4 Gy of X-ray and analyzed 72 h after irradiation. Data are mean ± s.e.m. *p < 0.05; **p < 0.01;***p < 0.001.

The catabolism of glucogenic amino acids produces TCA cycle intermediates and drives oxygen-consuming energy-producing pathways. Gln oxidation was measured by monitoring the real-time oxygen consumption rate (OCR) after Gln addition to substrate-free base medium (**Figure 1 d, e**). Gln supplementation caused a substantial increase in OCR over the baseline in all analyzed cell cultures confirming that Gln is an important regulator of mitochondrial activity (**Figure 1 f**).

In contrast to DU145, LNCaP RR and parental cells showed a significant difference in the levels of basal respiration, ATP production, and maximal respiration after Gln starvation (**Figure 1 e**). Analysis of Gln contribution to TCA cycle metabolites using ^13^C_5_-Gln and its conversion to M5 glutamate, M4 succinate, M4 malate and M4 citrate in PCa cells revealed that intracellular pool of the isotope-labelled metabolites was significantly higher in LNCaP RR cells (**Figure 1 g**). Next, we examined the intracellular pool of TCA cycle metabolites in parental and RR cells (**Figure 1 h**). Although α-KG levels were significantly increased in both RR cell sublines, the intracellular concentration of other metabolites, including succinate, pyruvate, and lactate, was only significantly higher in LNCaP RR compared to parental cells. In contrast, these metabolites were unchanged or decreased in DU145 RR cells, indicating that the entry of α-KG into the TCA cycle is repressed. Indeed, the α-KG/succinate ratio was robustly elevated in both DU145 and DU145 RR cells (**Figure 1 h**). ^13^C_5_-Gln metabolic flux analysis suggested that the Glutaminase (GLS)/TCA cycle ratio is significantly higher in DU145 RR than in the parental cells (**Supplementary Figure 1 d, e, f**).

In contrast to DU145 cells, Gln supplementation did not contribute significantly to α-KG and succinate production in parental LNCaP cells (**Figure 1 i**), most likely due to higher glucose contribution to α-KG production in LNCaP than DU145 cells as also shown by other studies ^24^.

These results indicate that radioresistance of DU145 PCa cells is associated with a decreased contribution of Gln to the TCA cycle and the ability to utilize Gln-derived α-KG for other purposes than energy homeostasis.

### Targeting of Gln metabolism for PCa radiosensitization

Beyond its role as a carbon donor in the TCA cycle, Gln also plays a vital role in redox control. In particular, Gln-derived Glu and α-KG are utilized for the biosynthesis of glutathione (GSH). GSH serves to neutralize reactive oxygen species (ROS) and protects cells from oxidative stress^5^. The curative potential of X-ray radiation depends on the balance between DNA damage and repair processes. Radiation-induced ROS production causes oxidative DNA damage. High intracellular GSH levels are exploited by cancer cells to resist radiotherapy, and GSH depletion is an attractive strategy for tumor radiosensitization ^25-27^. GSH production is increased in response to radiation-induced oxidative stress as one of the adaptive mechanisms^28^. In line with this, intracellular levels of Glu, α-KG, and succinate were induced by X-ray radiation in DU145 cells (**Figure 1 j**).

As Gln metabolism plays an essential role in GSH biosynthesis, it might also contribute to PCa radioresistance. Indeed, gene expression analysis showed that Gln deprivation results in the deregulation of genes, which control oxidative stress response and DNA damage response (DDR) in all analyzed cell lines (**Figure 2 a**). This deregulation was more pronounced in DU145 as compared to LNCaP cells, and in parental versus RR in both cell lines (**Figure 2 b**). Consistent with these data, Gln starvation lowered protein expression of crucial DDR regulators that we also described for other tumor models ^29^ (**Figure 2 c** and **Supplementary Figure 2 a**). Of note, expression of ataxia-telangiectasia mutated (ATM) kinase and related signaling networks were substantially more affected by Gln deprivation in DU145 than in LNCaP cells (**Figure 2 c** and **Supplementary Figure 2 a, b**). Consistently, after Gln starvation DU145 cells showed upregulation of the DNA damage marker γH2A.X.

**Figure 2.**
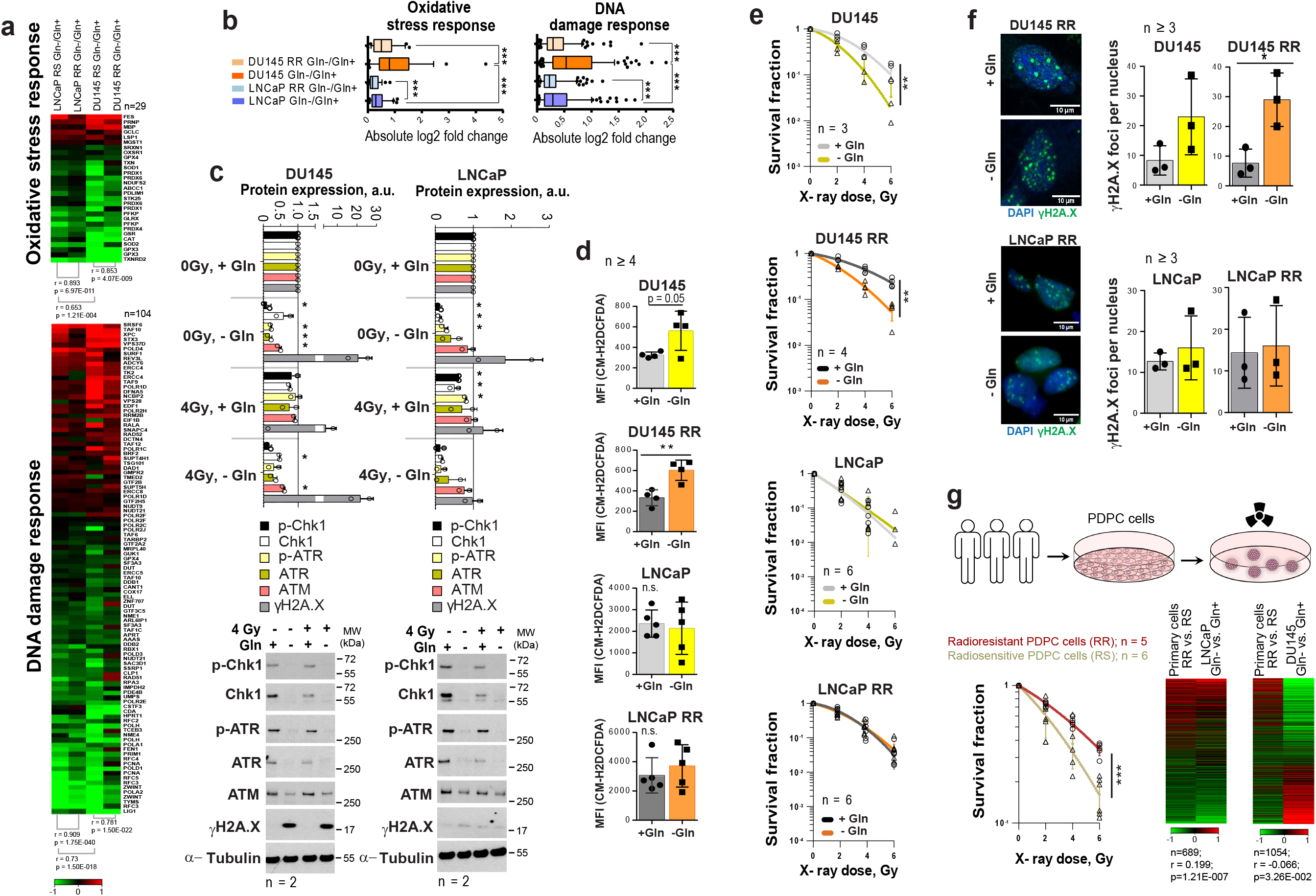
Gln deprivation increased oxidative stress and PCa cell sensitivity to X-ray radiation in Gln-dependent PCa cells. **a** Gene expression analysis showed that Gln deprivation results in consistent deregulation patterns of genes that control oxidative stress response and DNA damage response (DDR),although this deregulation is more pronounced for radiosensitive (RS) parental cells in each pair as compared to radioresistant (RR) cells, as well as for DU145 in comparison to LNCaP cells. **b** Absolute log2 fold change of genes involved in oxidative stress response and DNA damage response signatures. **c** Western blot analysis of key DDR regulators in response to Gln depletion for 72 h and quantification of the Western blot data. Cells grown in presence of glutamine were used as control. Data are mean ± s.d. **d** Gln starvation increased the intracellular level of ROS in DU145 and DU145 RR cells. Data are mean ± s.d. **e** Radiobiological colony-forming assay of PCa cells grown with or without Gln supplementation. Data are mean ± s.d. **f** Analysis of residual γH2AX foci number after 24 h of glutamine deprivation followed by 4 Gy of X-ray irradiation. Cells were analyzed 24 h after irradiation. Control cells were cultured in presence of glutamine. Data are mean ± s.d. Scale bars = 10 µm. **g** Comparative analysis of gene expression data from patient-derived primary cultures (PDPC) and Gln-deprived PCa cell lines. PDPC cultures were analyzed for their relative radioresistance using a 3-D radiobiological clonogenic assay. Data are mean ± s.d. Lists of genes significantly deregulated (p<0.05) in DU145 and LNCaP in response to glutamine starvation were compared with the list of genes differentially expressed (p < 0.05) between radioresistant (RR) and radiosensitive (RS) PDPC samples. For the genes present in both lists, Gln-/Gln+ and RR/RS fold changes were calculated. Finally, the Pearson correlation for those fold change values was determined. *p < 0.05; **p < 0.01; ***p < 0.001.

Gln starvation decreased the ratio of reduced glutathione (GSH) to oxidized glutathione (GSSG) in DU145 and LNCaP cells (**Supplementary Figure 2 c**), and activated an endoplasmic reticulum (ER) stress response, which was more pronounced in DU145 P and RR than in LNCaP cells and might serve as another source of ROS ^30^ (**Supplementary Figure 2 d**). Indeed, ROS levels were increased in DU145 P and RR, but not in LNCaP cells (**Figure 2 d**).

In addition to oxidative damage, Gln depletion causes accumulation of mutagenic DNA alkylation. This is due to reduction of intracellular α-KG, a co-substrate for Alkylated DNA Repair Protein AlkB Homolog family (ALKBH) dioxygenases, which directly repair DNA alkylation damage ^31^. Analyses of *ALKBH1* and *ALKBH3* expression levels after Gln starvation showed their specific upregulation in DU145, which can be partially rescued by α-KG repletion (**Supplementary Figure 2 e**). In line with this, the radiobiological clonogenic survival assays and γH2A.X foci analyses of DNA double strand breaks (DSB) showed that Gln-starved DU145 P and RR cells are more radiosensitive than control cells grown in Gln containing medium (**Figure 2 e, f** and **Supplementary Figure 2 f, g**). The radiosensitizing effect of Gln starvation can be partially rescued in DU145 cells by α-KG supplementation; however, this effect was not significant (**Supplementary Figure 2 h**). In contrast, LNCaP P and RR Gln-starved cells did not show significant radiosensitization (**Figure 2 e, f** and **Supplementary Figure 2 f, g, h**).

In order to further validate our observations from the established cell lines, we employed eleven patient-derived primary cell cultures (PDPC). First, we performed a functional analysis of PDPC radioresistance using the 3-D radiobiological clonogenic assay (**Figure 2 g**). Based on the results of this analysis, we grouped the PDPC as radiosensitive (RS, n = 6) or radioresistant (RR, n = 5). Comparative analysis of gene expression data from PDPCs and Gln-deprived PCa cell lines revealed that the gene signature of PDPC radioresistance inversely correlated with the Gln starvation gene signature from DU145 cells, but directly correlated with the Gln starvation gene signature from LNCaP cells (**Figure 2 g**). This correlation was not strong, yet significant. This finding indicated that in Gln-deprived conditions, Gln-dependent DU145 cells, but not LNCaP cells, might trigger molecular mechanisms associated with increased radiosensitivity in primary PCa cells.

Next, we analyzed PCa radiosensitivity after inhibition of the key Gln metabolism regulators, MYC or GLS. GLS was significantly upregulated in both LNCaP RR and DU145 RR cell lines than in the parental cells, whereas MYC was upregulated only in DU145 RR cells (**Supplementary Figure 3 a**). siRNA-mediated knockdown of MYC or GLS gene expression resulted in more pronounced radiosensitization in DU145 P and RR than in LNCaP P and RR cells (**Figure 3 a, b, and Supplementary Figure 3 b**). No additive effect was observed between individual and simultaneous knockdown of *MYC* and *GLS* in DU145 cells, suggesting that these two genes are involved in similar pathways regulating cell radiosensitivity. MYC regulates *GLS* directly through inducing its gene expression as it was shown for LNCaP cells or indirectly through regulation of Gln transporter *SLC1A5* as it was shown for both LNCaP and DU145 cells (**Supplementary Figure 3 c**). The radiosensitizing effect of *MYC* knockdown in LNCaP cells was reversed by Gln deprivation. It can be explained by additional MYC-dependent pro-survival mechanisms that can be partially activated by Gln depletion in this cell model. These mechanisms will be discussed in the next chapter.

**Figure 3.**
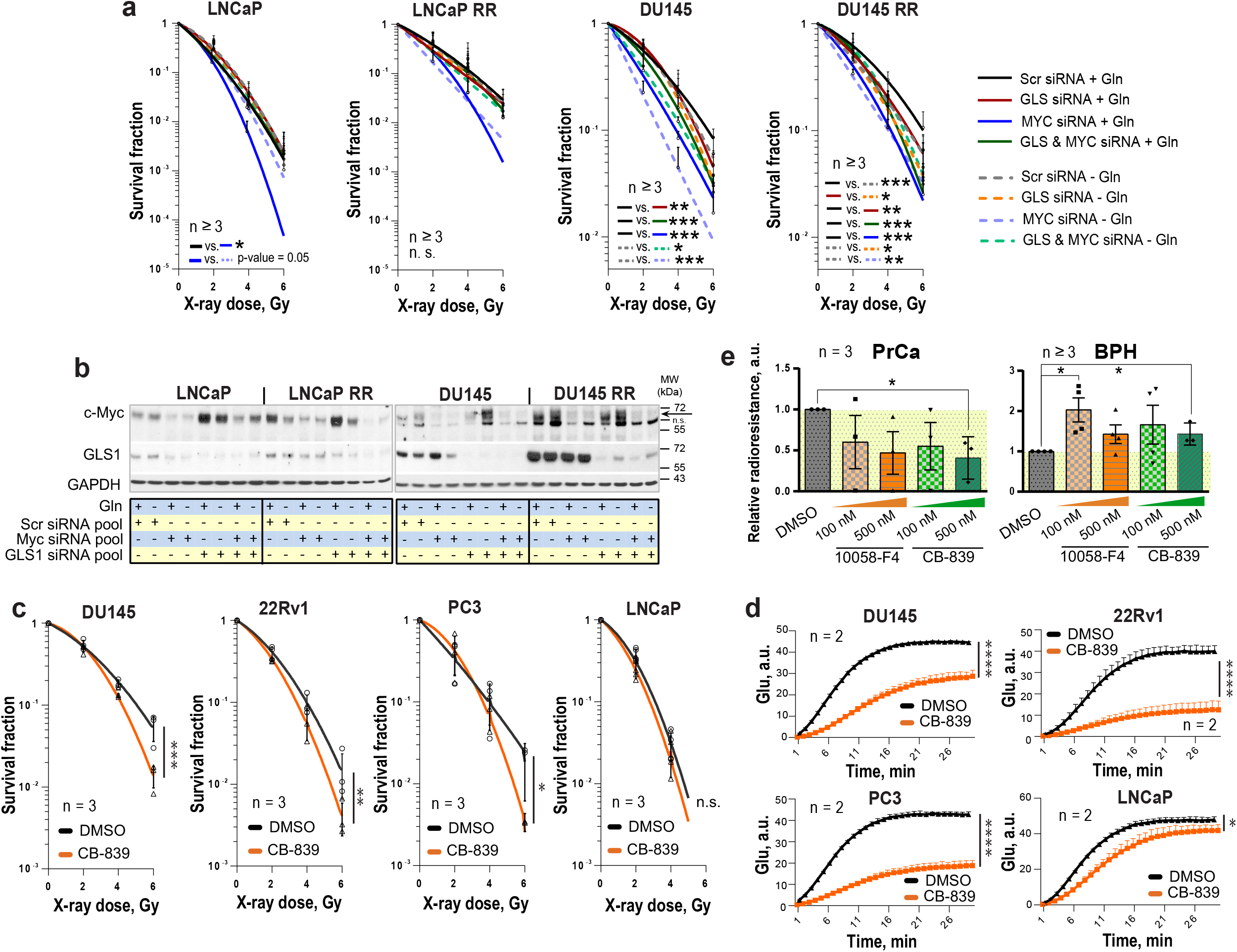
Genetic and chemical inhibition of Gln metabolism results in PCa cell radiosensitization. **a** Analysis of the relative PCa radiosensitivity in response to siRNA-mediated knockdown of *MYC* or *GLS* in cells grown with or without Gln supplementation for 24 h. Data are mean ± s.d. **b** Western blot analysis of PCa cells after siRNA-mediated knockdown of *MYC* or *GLS* gene expression, with or without Gln starvation. **c** Radiobiological colony-forming assay of PCa cells with or without inhibition of GLS. Cells were pretreated with GLS inhibitor CB-839 at LD_50_ concentrations for 48 h directly prior clonogenic analysis. Cells were irradiated with indicated X-ray doses directly upon plating. Data are mean ± s.d. **d** Cells were incubated overnight with LD_50_ concentrations of GLS inhibitor CB-839. The levels of Glu production were measured to confirm GLS enzymatic activity inhibition. Cells treated with DMSO were used as control. Data are mean ± s.d. **e** Radiobiological colony-forming assay of primary PCa and benign prostate hyperplasia (BPH) cultures with or without inhibition of MYC or GLS. Cells were pretreated with MYC inhibitor 10058-F4 or with GLS inhibitor CB-839 at indicated concentrations for 24 h directly prior 3-D clonogenic analysis. Cells treated with DMSO were used as control. Relative radiosensitivity was measured after 4 Gy of X-rays. Data are mean ± s.e.m.*p < 0.05; **p < 0.01, ***p < 0.001, ****p < 0.0001.

To analyze the effect of chemical inhibition of Gln metabolism on PCa radioresistance, four established PCa cell lines, DU145, 22Rv1, PC3 and LNCaP were treated for 48 h with LD_50_ doses of MYC inhibitor 10058-F4 or GLS inhibitor CB-839 and analyzed by radiobiological clonogenic assay (**Figure 3 c** and **Supplementary Figure 4 a, b**). The LD_50_ doses were determined for each cell line individually (**Supplementary Figure 4 c**), and the levels of Glu production were measured to confirm inhibition of GLS enzymatic activity (**Figure 3 d** and **Supplementary Figure 4 d**). GLS inhibition resulted in significant radiosensitization in all cell lines but LNCaP cells with a more pronounced effect in DU145 cells, whereas MYC inhibition had a radiosensitizing effect only in DU145 cells. Like the established cell lines, primary PCa cell cultures were significantly sensitized to irradiation by GLS inhibition. Both, GLS and MYC inhibition did not cause toxic effects on benign prostate hyperplasia (BPH) cells. Instead, it resulted in slightly but significantly increased radioresistance in BPH cultures (**Figure 3 e and Supplementary Figure 4 e**). We conclude that targeting the Gln metabolism results in the radiosensitization of the Gln-dependent PCa.

### Molecular mechanisms regulating PCa cell viability and radiosensitivity upon Gln starvation

The growth-inhibiting effect of Gln deprivation can be observed in all PCa cell lines tested (**Supplementary Figure 5 a**). To unravel the intracellular signaling mechanisms specifically regulated in Gln-dependent cells, we analyzed Gln starved cells for global transcriptomic changes. In accordance with previously published studies, deprivation of Gln led to the activation of apoptotic signaling ^32^, with stronger gene expression changes in the DU145 cell lines (**Supplementary Figure 5 b, c**). In support of these data, deprivation of Gln inhibited cell viability and increased apoptotic cell death of Gln-dependent DU145 P and RR cells, but not in LNCaP cell lines (**Figure 4 a, b, c**).

**Figure 4.**
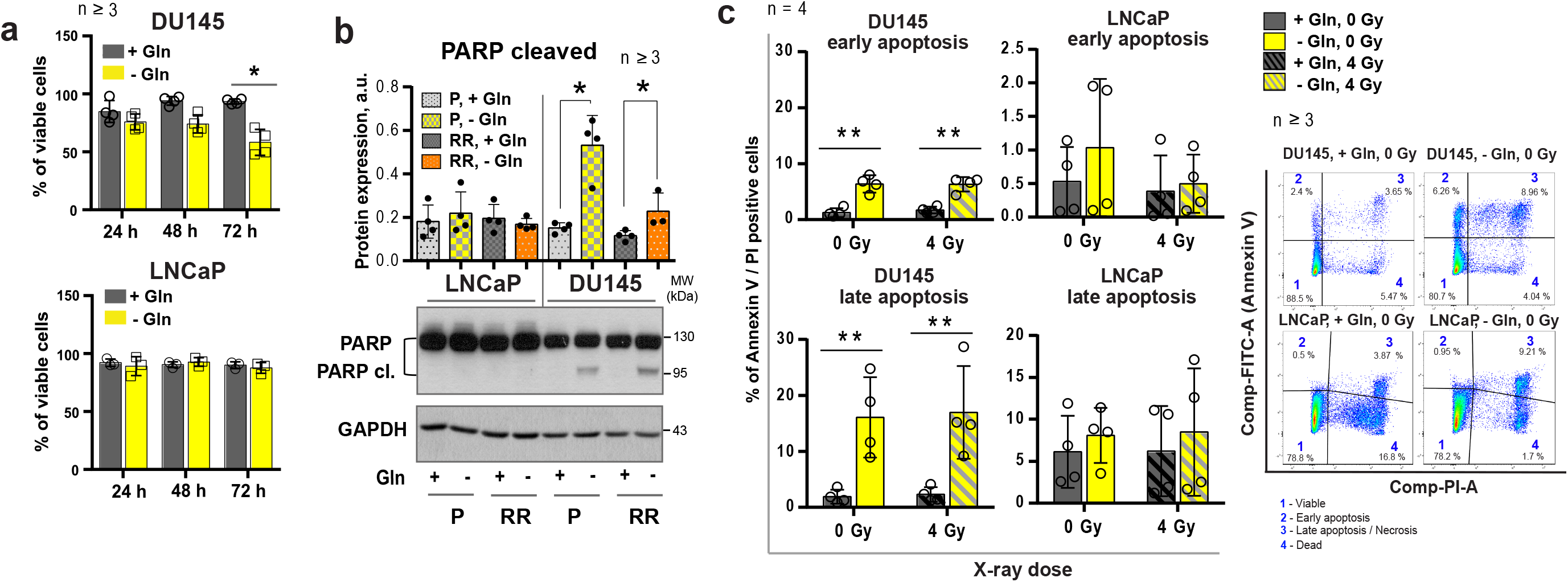
Regulation of PCa cell death upon glutamine deprivation. **a** Analysis of cell viability using Trypan blue exclusion assay. Data are mean ± s.d. **b** Western blot analysis of PARP cleavage after glutamine starvation of PCa cells for 24 h. Cells grown in the presence of glutamine were used as control. Data are mean ± s.e.m. **c** Cell death has been evaluated by flow cytometry using Annexin V-FITC and PI staining. Data are mean ± s.d. *p < 0.05; **p < 0.01.

It is known that activation of MYC-dependent autophagy serves as a pro-survival mechanism in response to nutrient starvation providing metabolic substrates including glutamate and α-KG ^33-35^. We assumed that activation of the autophagy could contribute to the LNCaP resistance to Gln catabolism inhibition. Gln depletion specifically activated autophagy in LNCaP but not in DU145 cells, as confirmed by Western blot detection of LC3B-II protein and fluorescence microscopy analysis of autophagosome formation (**Figure 5 a, b**). Analysis of gene expression revealed that DU145 cells have undetectable mRNA levels of *ATG5, a* key component for autophagy^36^ (**Figure 5 c**). Inhibition of autophagy by chloroquine (CQ), which prevents autolysosome maturation, or by siRNA-mediated *ATG5* knockdown resulted in the LNCaP radiosensitization in response to Gln starvation (**Figure 5 d, e, f** and **Supplementary Figure 5 d, e**). These results suggest that autophagy might acts as a pro-survival mechanism under Gln deprivation conditions, and inhibition of autophagy increases radiosensitivity in Gln-independent PCa cells.

**Figure 5.**
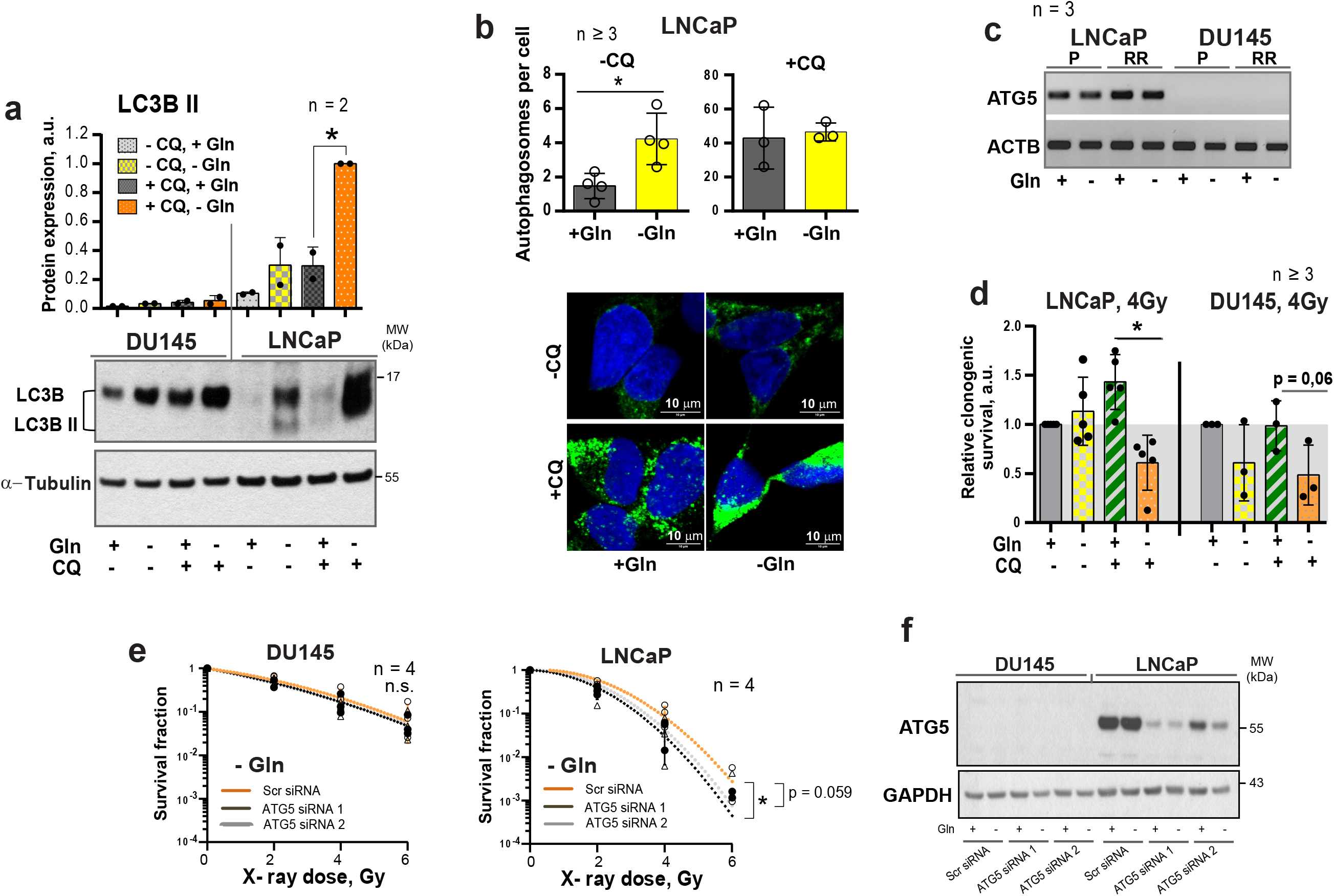
Regulation of autophagy in response to glutamine deprivation. **a** Western blot detection of LC3B-II protein in PCa cells grown with or without glutamine starvation for 24 h. Cells treated with autophagy inhibitor Chloroquine (CQ) at a concentration of 20 μM for 24 h were used as control. Data are mean ± s.d. **b** Monitoring LC3-positive puncta (autophagosomes) in LNCaP cells starved for Gln for 24 h, treated with CQ at a concentration of 20 μM for 24 h, or both. Data are mean ± s.d. **c** PCR analysis of ATG5 expression in LNCaP and DU145, parental (P) and radioresistant (RR) cells. **d** Analysis of the radiosensitizing effect of autophagy inhibition in Gln starved cells. Cells were cultured in medium with or without Gln for 24 h before irradiation. For autophagy inhibitions, cells were treated with CQ at a concentration of 10 μM for 1 h before and 1 h after irradiation with 4 Gy of X-rays. Sham irradiated cells were used as control. Cells were used for radiobiological colony-forming assay. Data are mean ± s.d. **e** Radiobiological clonogenic analysis of PCa cells after ATG5 knockdown in combination with Gln starvation. Cells were transfected with two ATG5 siRNA or scrambled siRNA for 24 h and then supplemented with fresh media without glutamine. 24 h later, cells were used for radiobiological colony-forming assay. Data are mean ± s.d. **f**. Representative Western blot analysis of DU145 and LNCaP cells transfected with ATG5 siRNA or scrambled siRNA. *p < 0.05.

### Gln metabolism contributes to tumor cell reprogramming and regulation of CSCs

α-KG is an essential co-factor in the regulation of the epigenetic enzymes, such as α-KG-dependent dioxygenases. Many of these chromatin-modifying enzymes are involved in the maintenance of stem cells, including CSCs ^14, 37, 38^. Enzymatic function of α-KG-dependent dioxygenases, such as Jumonji C (JmjC)-domain-containing histone demethylases, depends on the entry of glutamine-derived α-KG into the TCA cycle ^14^. In our study, we found that DU145 P and DU145 RR cells have a high α-KG / succinate ratio. The elevated ratio of α-KG / succinate suggests that α-KG is less oxidized in mitochondria and instead used for other purposes, such as epigenetic regulation (**Figure 1 i**). In support of this assumption, Gln starvation led to increased trimethylation of histone H3 at lysine 27 (H3K27me3) (**Figure 6 a**) and induced deregulation of genes involved in CSC maintenance (**Supplementary Figure 6 a**). This regulation is reciprocal, and inhibition of histone methylation by the epigenetic CSC inhibitor DZNeP ^23, 39^ also downregulated GLS expression (**Supplementary Figure 6 b)**.

**Figure 6.**
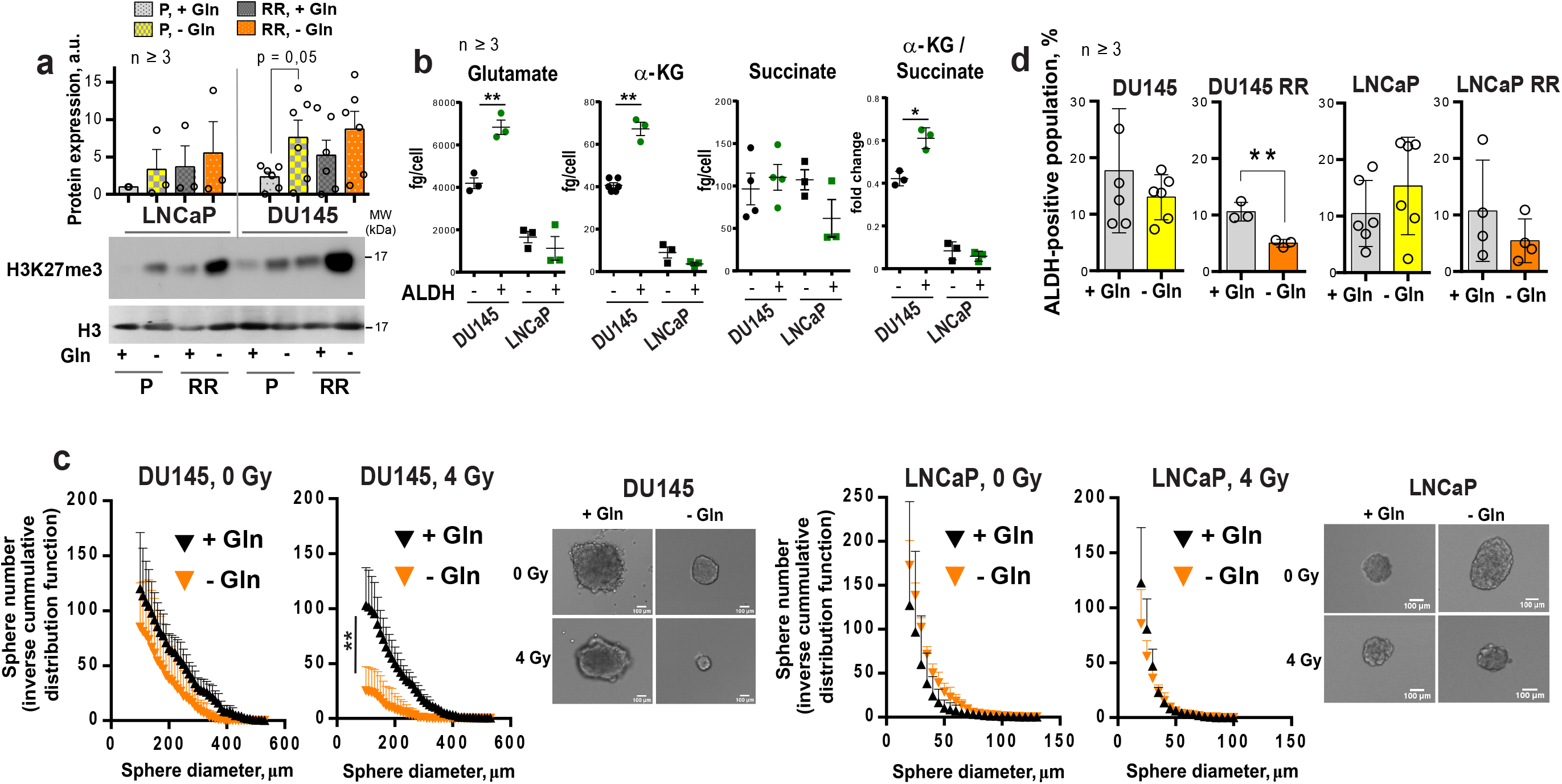
Glutamine metabolism regulates tumor cell reprogramming and *in vitro* CSC properties. **a** Western blot analysis of H3K27me3 levels and quantification of the Western blot data. H3K27me3 levels were normalized to the levels of total H3. Data are mean ± s.e.m. **b** The intracellular levels of Glu, α-KG and succinate measured in ALDH^+^ and ALDH^-^ cells by LC-MS/MS. Data are mean ± s.d. **c** Glutamine starvation significantly decreased the number and size of formed tumor spheres after 4 Gy of X-ray irradiation in DU145 cells. Inverse cumulative distribution function (CDF) was defined as a number of spheres larger than a given size. Representative images of spheres are shown for DU145 and LNCaP cells, which were depleted or replenished with Gln for 72 h in combination with 4 Gy of X-ray or sham irradiation. Data are mean ± s.d. **d** Flow cytometry analysis of ALDH positive cell population in response to Gln starvation for 72 h. Cells replenished with Gln were used as control. Data are mean ± s.d. *p < 0.05; **p < 0.01.

We and others have shown that PCa cells with high ALDH activity (ALDH^+^) have functional characteristics of CSCs such as high *in vitro* sphere-forming capacity and *in vivo* tumor-initiating potential, as well as high radioresistance ^22, 23, 40^. Similar to RR PCa cells, ALDH^+^ cells have activation of Gln metabolism, which is reflected by a significantly higher intracellular concentration of Glu and α-KG in ALDH^+^ compared to ALDH^-^ cells. Moreover, in comparison to their ALDH^-^ counterparts, DU145 ALDH^+^ cells have a significantly elevated α-KG / succinate ratio (**Figure 6 b**). Similar to the ALDH^+^ population, PCa cells growing under sphere-forming conditions also showed substantial changes in amino acid metabolism (**Supplementary Figure 6 c**). In agreement, Gln-dependent DU145 cells cultured in the absence of Gln demonstrated inhibition of sphere-forming properties, most significantly after cell irradiation (**Figure 6 c** and **Supplementary Figure 7 a**). Gln starvation resulted in the inhibition of the ALDH^+^ cell populations in DU145 RR cells (**Figure 6 d**). Primary patient-derived PCa cells (PDPC) cultured under Gln-conditions showed a trend toward downregulation of ALDH1A1 protein expression. In contrast, Gln deprivation of primary BPH cultures resulted in significantly increased ALDH1A1 expression, partially explaining the high resistance of BPH to chemical inhibition of Gln metabolism (**Supplementary Figure 7 b** and **Figure 3 e**). Moreover, treatment with a chemical inhibitor of MYC, 10058-F4, lowered the radiation-induced ALDH^+^ cell subsets in DU145 cells (**Supplementary Figure 7 c**).

To test whether inhibition of Gln metabolism can influence tumor-initiating properties, PCa cells were pre-incubated in the presence or absence of Gln for 72 h, irradiated with 6 Gy of X-rays or sham irradiated, and injected into NMRI nude mice. Analysis of tumor growth in these xenograft models showed that Gln deprivation resulted in a significant reduction of tumor take rate and even in loss of tumor-forming capacity of DU145 P and RR cells after combining Gln starvation with cell irradiation (**Figure 7 a, b** and **Supplementary Figure 7 d**). *In vivo* limiting dilution analysis confirmed that Gln starved DU145 P and RR cells showed a significantly lower frequency of tumor-initiating cells than control Gln replenished cells (**Figure 7 c)**. In contrast, LNCaP cells neither significantly lost their tumor-initiating properties nor demonstrated radiosensitization after Gln deprivation in these *in vivo* assays (**Figure 7 a, b, c** and **Supplementary Figure 7 d**). These results suggest that Gln metabolism contributes to epigenetic reprogramming and regulation of CSC populations in Gln-dependent PCa cells.

**Figure 7.**
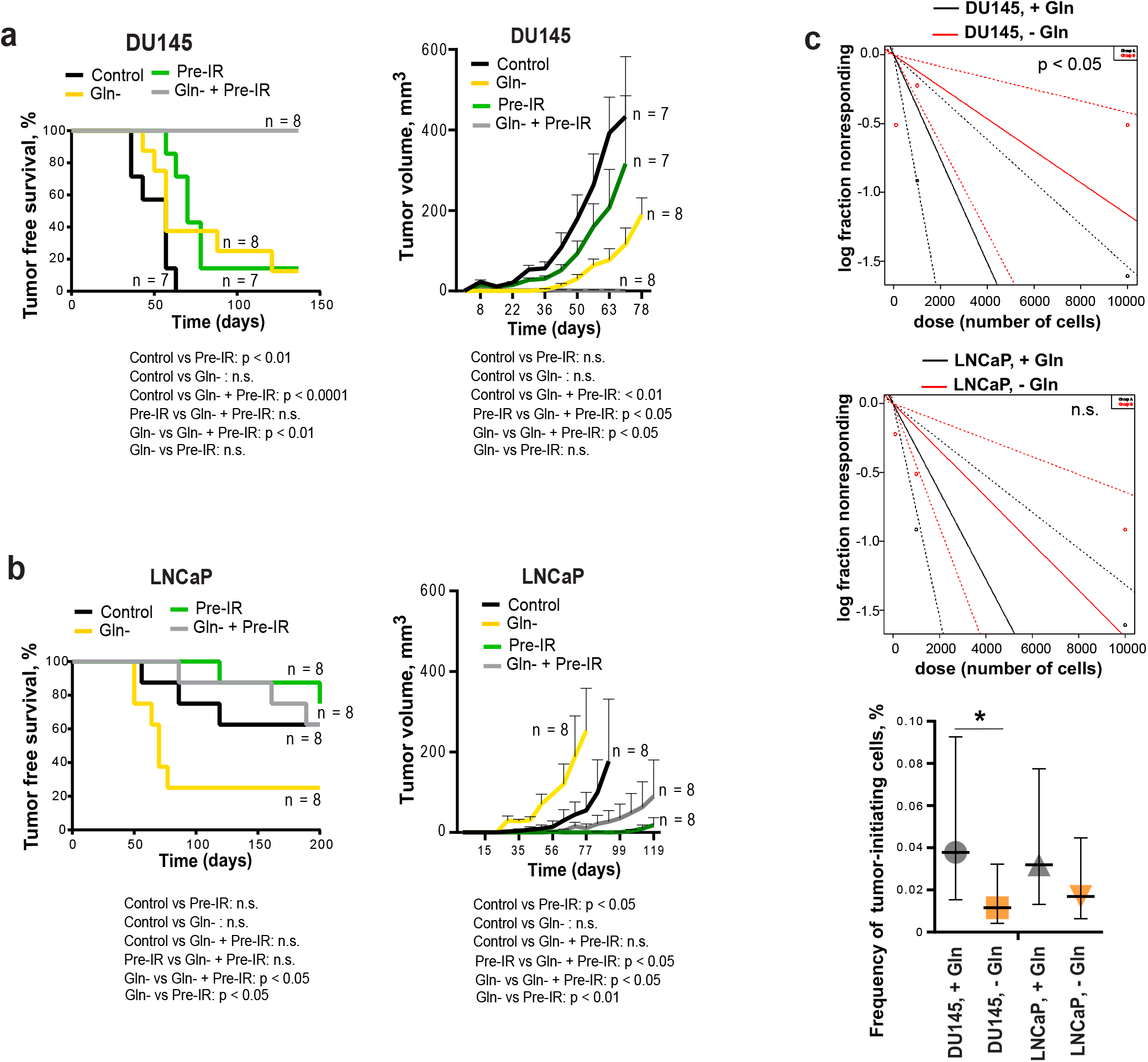
Glutamine metabolism regulates tumor growth and *in vivo* CSC frequency. Analysis tumor take and tumor growth for DU145 (**a**) and LNCaP (**b**) in NMRI (nu/nu) xenograft models. PCa cells were pre-incubated in the presence or absence of Gln for 72 h, irradiated with 6 Gy of X-rays or sham irradiated, and injected into the mice in Matrigel as 5×10^3^ viable cells/implant. **c** Analysis of tumor-initiating cell frequency. PCa cells were pre-incubated in the presence or absence of Gln for 72 h and injected into the NMRI (nu/nu) in Matrigel as 10^2^, 10^3^ and 10^4^ viable cells/implant. Limiting dilution assay calculations were performed using the ELDA software and presented as plots of the log fraction of mice bearing no tumors at day 137 (DU145) or day 202 (LNCaP) (log fraction nonresponding) as a function of the number of cells injected in mice (dose). The slope of the lines indicate log-active cell fraction, dotted lines indicate 95% confidence interval (c.i.). Frequency of tumor-initiating cells (%) is shown on the bottom panel. Data are mean ± c.i. *p < 0.05.

### Gln metabolism as a marker of tumor radioresistance

The Myc transcriptional factor coordinates the expression of genes necessary for tumor cells to become glutamine-dependent ^9^. The acquisition of cell radioresistance is associated with *MYC* upregulation in Gln-dependent PCa cells. In turn, Gln depletion in these cells inhibited the MYC-dependent transcriptional program (**Supplementary Figure 8 a, b**).

Other studies previously demonstrated that high expression of *MYC* contributes to the development of highly aggressive prostate tumors with elevated metastatic potential ^10^, and that circulating tumor cells (CTCs) in PCa patients have recurrent *MYC* gene amplification ^41^. In our study, we conducted microarray-based comparative genomic hybridization (aCGH) analysis of single CTCs isolated from the blood of two patients with metastatic PCa. We found that CTCs from the same patient have heterogeneous genomic profiles, whereas CTCs with a high gain at the cytoband 8q24.21 (*MYC* locus) were detected in both analyzed patients (**Figure 8 a**).

**Figure 8.**
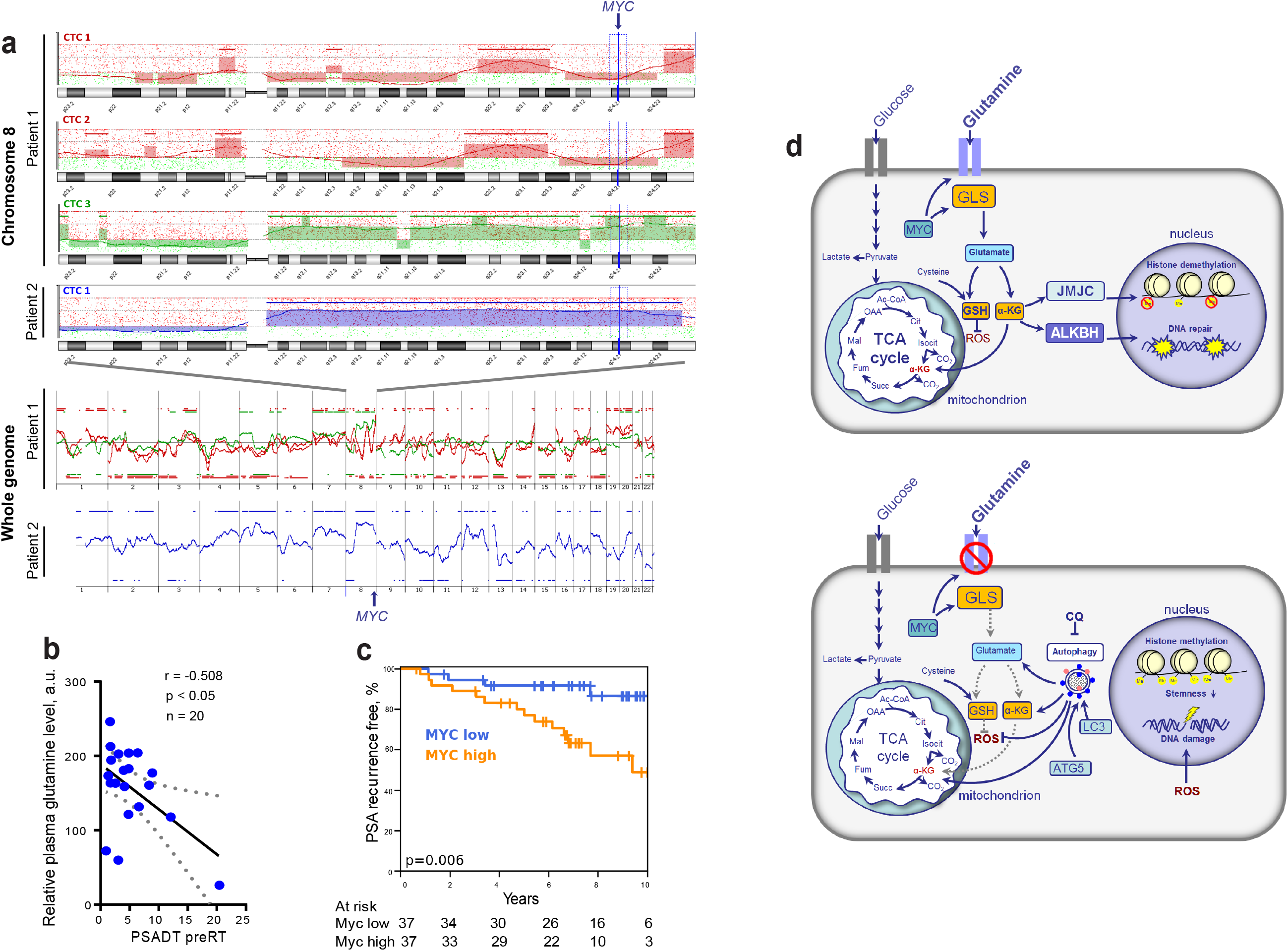
Gln metabolism as a prognostic biomarker in PCa. **a** Microarray-based comparative genomic hybridization (aCGH) analysis of single CTCs isolated from the blood of two patients with metastatic PCa by CellCelector system. **b** Blood plasma level of Gln measured by LC-MS/MS negatively correlated with prostate-specific antigen doubling time (PSA-DT) in PCa patients prior to radiotherapy. **c** *MYC* expression was correlated with PSA relapse-free survival in patients with PCa treated with radiotherapy at the Department of Radiotherapy and Radiation Oncology, University Hospital Carl Gustav Carus and Faculty of Medicine (n = 74). **d** The role of Gln metabolism in regulation of PCa radiosensitivity. Gln serves not only for energy production in the TCA cycle but also for maintaining the redox state by contribution to GSH production, DNA repair by regulation of ALKBH dioxygenases, and tumor epigenetic resetting by control of JmjC-domain-containing histone demethylases. The lack of glutamine leads to the oxidative stress, depletion of CSCs, and tumor radiosensitization. The blockade of Gln metabolism results in the activation of the pro-survival autophagy pathway, and autophagy inhibition can be a promising way for PCa radiosensitization if combined with Gln metabolism targeted drugs. GSH - glutathione; ROS – reactive oxygen species; α-KG – alpha ketoglutarate; TCA - tricarboxylic acid cycle; CQ – chloroquine; GLS – glutaminase; JMJC - Jumonji C (JmjC)-domain-containing histone demethylases; ALKBH - AlkB Homolog 1.

Next, we tested whether blood plasma levels of Gln can be used for identifying patients at risk of disease progression. For this we measured plasma concentrations of 35 amino acid metabolites in castration resistant PCa patients (n = 20) and correlated these measures with prostate specific antigen doubling time (PSA-DT). A short PSA-DT is an important predictor of the metastatic development ^42^. PCa patients were diagnosed with castration resistant oligorecurrent PCa and received local ablative radiotherapy, while the course of androgen-deprivation therapy was continued. The clinical characteristics of PCa patients are provided in **Supplementary Table 1**. We found that among all measured amino acids in blood plasma, only Gln showed significant negative correlation with PSA-DT before radiotherapy (**Figure 8 b**). This relationship was not observed after the course of radiotherapy that can be partially explained by the heterogeneous impact of radiation on PCa cell metabolism and small patient cohort (n = 15) (**Supplementary Table 2**).

To understand the possible implications of Gln metabolism-related genes as biomarkers for PCa progression after radiation therapy, we analyzed The Cancer Genome Atlas (TCGA) gene expression dataset and found that high combined expression of both *MYC* and *GLS* is significantly associated with decreased relapse-free survival in PCa patients treated with radiotherapy (n = 56) (**Supplementary Figure 8 c**). Consistent with this finding, *MYC* expression correlated with PSA relapse-free survival in patients with PCa treated with curatively-intended radiotherapy (n = 74) (**Figure 8 c**). In this study, *MYC* expression was analyzed by nanoString technology using formalin-fixed paraffin-embedded (FFPE) tissues of PCa patients. All patients were diagnosed with a localized intermediate-or high-risk PCa and had a minimum follow-up of 7 years (**Supplementary Table 3**). These findings suggest that Gln metabolism-related biomarkers are associated with tumor progression and biochemical recurrence of PCa after radiotherapy.

## Discussion

Radiation therapy is one of the main pillars of curative treatment for clinically localized PCa. However, treatment-related normal tissue toxicities as well as therapy resistance of the tumors still constitute an important clinical and scientific challenge. Prediction of cancer recurrence after therapy is important for personalized PCa treatment ^43^. The current risk stratification for PCa used for treatment decision-making is based on prostate-specific antigen (PSA) level, the tumor, node and metastasis (TNM) staging, and Gleason score. Although of great clinical importance, this stratification system is associated with significant heterogeneity in clinical outcomes. Therefore, additional biological markers that predict tumor progression after radiation therapy offer clinical value ^1^.

Reprogramming of cellular metabolism is one of the main hallmarks of tumor cells playing a vital role in tumor development and therapy resistance ^17^. Deregulated metabolic pathways in PCa might provide specific biomarkers for identifying patients most likely to have a reduced response to radiotherapy and specific therapeutic targets to enhance the efficacy of conventional treatment approaches. An abundant supply of amino acids is critical to sustain highly proliferative tumor tissues, and reprogramming of amino acid metabolism emerged as a valuable source of biomarkers and therapeutic targets for cancer treatment ^44-46^.

Gln is the most abundant amino acid in human tissues ^47^. It fuels energy production through the TCA cycle and is an important donor of nitrogen and carbon for the biosynthesis of amino acids, nucleotides, and fatty acids ^48^. Although Gln can be synthesized by normal tissues, it becomes conditionally essential for highly proliferative tumor tissues under nutrient-deprived conditions. Gln-dependence was demonstrated not only for cancer cells in culture, but also for *in vivo* tumor models ^6-8, 49, 50^. This is, in particular, true for PCa tissues with Myc-induced metabolic reprogramming ^51, 52^. Independent analyses of large patient cohorts by different institutions revealed that the majority of PCa tissues possess an upregulation of *MYC* mRNA compared to matched benign prostatic specimens ^53^. These elevated MYC levels are caused by gene amplification, which is found in about 30% of castration-resistant PCa ^54^, or by epigenetic regulation of MYC expression ^10^. MYC acts as a broad transcriptional regulator, which contributes to tumor cell adaptation to nutrient-deprived conditions through regulation of metabolic genes, including glutaminase *GLS1* ^55^ and Gln transporters ^56^.

Our data indicate that high expression levels of *MYC* and *GLS1* in PCa patients are significantly associated with decreased relapse-free survival after radiotherapy. Our study showed that Gln metabolism is a critical regulator of PCa radioresistance as Gln-derived α-ketoglutarate (α-KG) is important for GSH production and ROS scavenging. Inhibition of Gln metabolism leads to the induction of stress responses, accumulation of DNA DSBs, and an increase in susceptibility to ionizing radiation. GLS1 inhibitor CB-839 is currently in phase 1 clinical evaluations in patients with advanced solid tumors (e.g., NCT02071862, NCT03875313, NCT02861300) showing encouraging clinical activity and tolerability, and further studies are warranted to establish its utility as a radiosensitizing treatment. One of our important findings is that in contrast to cancer tissues, chemical inhibition of Gln metabolism with GLS1 and MYC inhibitors in BPH cells did not result in radiosensitization or even increase cell radioresistance. Thus, this treatment approach may potentially improve the therapeutic ratio in a radiotherapy setting.

The different effect of GLS1 and MYC inhibitors in cancer and BPH cells can be partially attributed to the significant upregulation of ALDH1A1 expression in BPH tissues in response to Gln starvation. Recent studies demonstrated that ALDH proteins are important for energy metabolism in tumor and normal tissues ^57-59^, and that reduced nicotinamide adenine dinucleotide (NADH) produced by ALDH is essential for maintaining redox homeostasis ^60, 61^.

ALDH proteins are also potent regulators of CSCs. Our previous findings showed that the PCa cells with high ALDH activity are enriched for CSCs and possess self-renewal, tumor-initiating, and radioresistant properties ^22, 23^. One of the key findings from the present work is that radioresistant PCa CSCs are highly dependent on Gln supplementation for their tumorigenic properties, have high Gln uptake, and can be depleted by Gln deprivation. The subset of CTCs that initiates development of metastasis has CSC properties ^62^, and CTCs from PCa patients with metastatic disease carry *MYC* gene amplifications. In line with this finding, Gln levels in blood plasma significantly correlated with a short PSA-DT in PCa patients before radiotherapy. Consistently, it was previously reported that chemical inhibition of Gln transporter ASCT2 results in inhibition of tumor growth and metastatic spread in xenograft prostate tumor models ^63^, and that high blood levels of Gln, which is derived from tumor stroma, are associated with resistance to androgen deprivation therapy in PCa patients ^64^. Our data showed that the radioresistant phenotypes in PCa cells and CSCs are associated with a high production of α-KG, which is implicated in epigenetic resetting by regulation of histone methylation, and that Gln depletion led to the accumulation of the repressive chromatin mark H3K27me3. These results are consistent with recent findings demonstrated that intracellular α-KG level is critical for the prevention of embryonic cell differentiation as it serves as a co-factor for Tet family DNA demethylases and Jumonji C-domain containing histone demethylases ^14^. Interestingly, decreased levels of H3K27me3 in human PCa tissues are associated with high MYC expression as well as with high Gleason score and pathological stage ^65^.

Our study and other findings demonstrated that prostate cancer cells exhibit different sensitivities towards inhibition of MYC and Gln deprivation ^66^. Culturing sensitive PCa cells such as DU145 in Gln-deprived medium led to a substantial drop in the intracellular levels of Glu and α-KG, and consequently induced ER stress response, oxidative stress, and apoptosis. In contrast, PCa cells insensitive to Gln starvation like LNCaP, activated adaptive mechanisms, namely pro-survival autophagy, and maintained intracellular Glu and α-KG levels (**Figure 8 d**). We demonstrated for the first time that inhibition of autophagy in these cells by genetic knockdown of the *ATG5* gene or by chemical inhibition with chloroquine significantly increased the radiosensitizing effect of Gln deprivation. This observation suggest that autophagy activation in response to inhibition of Gln metabolism acts as a pro-survival mechanism, and its inhibition might be a promising approach for further PCa radiosensitization. Of note, recent studies showed that the diabetes drug metformin substantially suppressed MYC expression, reduced GLS activity, and inhibited autophagy in tumor cells ^67-69^. Metformin treatment improved radiotherapy response in mice and was associated with a decrease in early biochemical relapse. Therefore, metformin might represent a promising metabolic treatment for prostate tumor radiosensitization acting through inhibition of Gln metabolism and autophagy ^70^.

Overall, our findings indicate that genes regulating Gln metabolism in PCa may serve as potential biomarkers for predicting tumor radiosensitivity and also represent promising targets for tumor radiosensitization. We found that the combination of Gln metabolism inhibitors with irradiation did not cause toxic effects on BPH cells indicating a reduced risk of normal tissue complication for future clinical application. Future research is needed to validate our findings by *in vivo* experiment combining GLS inhibition and radiotherapy in a setting of established tumors, in particular, in patient-derived xenograft models. Besides, further clinical studies are warranted to ascertain the potential therapeutic utility of Gln metabolic pathways for PCa radiosensitization and the implication of Gln plasma levels as well as MYC and GLS1 expression levels as biomarkers of tumor aggressiveness and recurrence after radiotherapy.

## Methods and Materials

### Cell lines

Prostate cancer cell lines DU145, PC3 and LNCaP, were obtained from ATCC. Their radioresistant (RR) derivatives were generated by *in vitro* selection using fractionated X-ray irradiation as descried earlier ^22^. Cell were cultured in Dulbecco’s modified Eagle’s medium (DMEM; Sigma-Aldrich, USA) or RPMI-1640 medium (Sigma-Aldrich), respectively, supplemented with 10 % fetal bovine serum (FBS; Fischer Scientific) and 2 mM L-glutamine (Sigma-Aldrich). For glutamine starvation experiments, cells were cultured in DMEM or RPMI medium supplemented with 10 % dialyzed FBS (Sigma-Aldrich). Cells cultured in DMEM or RPMI medium supplemented with 10 % dialyzed FBS (Sigma-Aldrich) and 2 mM L-glutamine (Sigma-Aldrich) served as control for these experiments. Cells were cultured in humidified incubators at 37°C and 5 % CO_2_. Before experimentation, the relative radiosensitivity of the non-irradiated parental cells and age-matched RR cells was validated by radiobiological colony-forming assay. All cell lines were tested mycoplasma negative and genotyped using microsatellite polymorphism analyses.

### Tumor growth in mice xenograft models

The animal experiments were conducted in the SPF animal facility of OncoRay, Dresden according to the institutional guidelines and the German animal welfare regulations (protocol number TVV2014/30). The experiments were performed using 8-to 10-week-old male NMRI (nu/nu) mice. For immunosuppression, the animals received total body irradiation one day before tumor transplantation with 4 Gy (200 kV X-rays, 0.5 mm Cu filter, 1.3 Gy/min). Subcutaneous (s.c.) xenograft tumors were established by injection of tumor cell suspension in Matrigel. Limiting dilution analysis was performed via s.c. injection of 10.000, 1.000 and 100 tumor cells into NMRI nu/nu mice. Tumor growth was observed for 200 days or until tumors reached maximum allowed tumor volume. For analysis of tumor growth after Gln starvation and irradiation, PCa cells were pre-incubated in medium with or without Gln for 72 h, irradiated with 6 Gy of X-rays or sham irradiated, and injected into the mice in 100 µL of Matrigel. Tumor growth was measured once per week using electronic caliper by measuring length (L), width (W) and height (H). The tumor volume (V) was calculated based on the equation for hemiellipsoid tumors: V = (L * W * H)/2. Tumor take rates were compared using Kaplan-Meier analysis in tumor-bearing mice with a threshold of 100 mm^³^. P values were determined using Log-rank (Mantel-Cox) test. The frequency of tumor-initiating cells was calculated based on tumor-bearing mice using Extreme Limiting Dilution Analysis (ELDA) software ^71^.

### Primary tissue cultures and 3-D clonogenic analysis

Clinical material was collected with informed consent and approval from the local ethics committee (Institutional Review Board of the Faculty of Medicine, Technische Universität Dresden, EK152052013). The clinical characteristics of PCa patients are provided in **Supplementary Table 4**. Primary prostate cancer and adjacent normal tissues (benign hyperplasia, BPH) were pathologically evaluated upon radical prostatectomy. Patient-derived prostate cancer and benign prostatic hyperplasia cell cultures were established as described previously ^72^, and maintained in primary culture medium containing keratinocyte growth medium (Thermo Fisher) supplemented with 5 ng/mL of EGF (Peprotech), bovine pituitary extract (Life Technologies), 2 ng/mL leukemia inhibitory factor (Peprotech) 2 ng/mL stem cell factor (Peprotech), 100 ng/mL cholera toxin (Sigma), and 1 ng/mL granulocyte macrophage colony-stimulating factor (GM-CSF) (Miltenyi Biotec). Primary cell cultures were maintained in the presence of mouse embryonic fibroblasts (STO) irradiated with 60 Gy of X-rays, and used for the experiments at the early (up to 5) passages. After expansion, cells were plated at a density of 8 × 10^4^ cells/mL in collagen I–coated 6 well plates (Corning) without feeder cells. Cells were pretreated with MYC inhibitor 10058-F4 or with GLS inhibitor CB-839 at concentrations of 100 nM or 500 nM for 24 h directly prior 3-D clonogenic analysis. For 3-D clonogenic assays, cells were harvested with 0.25% trypsine-EDTA solution (Gibco) and centrifuged at 1000 rpm for 5 min. The supernatant was discarded, cells were re-suspended in the complete primary culture medium, counted, mixed with Matrigel diluted 1:20 with DMEM medium, and seeded onto 96 ULA plate (Corning Inc.) as 1000 cells per well. Next day cells were carefully overlaid with 50 µl of the complete primary culture medium and irradiated with 4 Gy of X-rays or sham-irradiated. Colonies were analyzed after 7 to 10 days.

### Cell viability assay

Cells were seeded in 96-well plates (3000 cells/well) into DMEM or RPMI with 10% dialyzed FBS and without glutamine and incubated at 37°C and 5 % CO_2_. Cells grown in glutamine-containing media were used as control. Relative cell viability was measured every day using 3-(4,5-Dimethylthiazol-2-yl)-2,5-diphenyltetrazolium bromide (MTT) assay. For this assay, 10µL of thiazolyl blue tetrazolium bromide(Sigma-Aldrich) aqueous solution were added directly to the culture media; cells were incubated 3h in the humidified CO_2_ incubator until the purple formazan crystals were formed, as described previously ^22^. Then the media was removed, and formazan crystals were dissolved in the mixture of 100 % ethanol: 100 % DMSO (1:1). The absorbance was measured at TECAN plate reader (Tecan Trading AG) at 560 nm wavelength.

### Trypan blue exclusion assay

To analyze the number of viable cells after glutamine inhibition, cells were cultured in 6-well plates. 24 h after seeding, cells were washed by PBS, and supplemented with fresh media with or without glutamine. Cells grown in glutamine-containing media were used as control. The number of cells and their viability were quantified by Trypan blue staining.

### Chemical inhibition

For the chemical inhibition of c-Myc, 10058-F4 compound (Sigma, USA) was used, and for the inhibition of GLS1, CB-839 compound (Selleck Chemicals LLC) was used; both inhibitors were dissolved in 100% DMSO. The LD_50_ values were determined for DU145, 22Rv1, PC3 and LNCaP prostate cancer cell lines. For this, 10^5^ cells per well for PC3, DU145 and 22Rv1 cell lines and 3 × 10^5^ cells per well for LNCaP cells were seeded into 6-well plate in appropriate medium with 3% FBS and incubated overnight. The same medium containing different concentrations of the compounds from 0 to 100 μM or DMSO as control was added to the cells next day for DU145, 22Rv1, PC3 cells and two days later for LNCaP cells. Cells were incubated in the presence of inhibitors for 48 h, and counted by using CASY cell counter. LD50 values (50% lethal dose) were determined by non-linear regression using GraphPad Prism software (San Diego, USA). Cells pretreated with inhibitors at LD_50_ concentration for 48 h were directly used for clonogenic survival analysis. For analysis of the intracellular Glu levels, 2.5 × 10^4^ cells per well were seeded into 96-well plate in DMEM with 10% FBS, treated with LD_50_ concentrations of 10058-F4 or CB-839 for 12 h in DMEM without FBS then analyzed by Glutamate-Glo^™^ Assay (Promega) according to the manufacturer’s instructions. Cells treated with DMSO were used as control.

### Clonogenic cell survival assay

For glutamine starvation experiments, cells were cultured in glutamine starvation medium for 24h. Cells grown in glutamine-containing media were used as control. Cells were harvested with 0.25% trypsine-EDTA solution (Gibco) and centrifuged at 1000 rpm for 5 min. The supernatant was discarded, cells were re-suspended in complete DMEM or RPMI, counted, and seeded onto 6-well plates (1000 or 2000 cells per well). Next day plates were irradiated with 0 Gy, 2 Gy, 4 Gy and 6 Gy of X-rays (Yxlon Y.TU 320; 200 kV X-rays, dose rate 1.3Gy/min at 20 mA, filtered with 0.5 mm Cu). After the irradiation, cells were left in the humidified CO_2_ incubator for 7 to 10 days. Colonies were fixed with 10% formaldehyde in PBS for 30 min and stained with a water solution of 0.05% crystal violet for 30 min at room temperature, then dried on the bench and counted manually. Results were presented as survival fractions depicted on a logarithmic scale and plotted against applied X-ray doses.

### Cell rescue with dimethyl-alpha-ketoglutarate

Cells were seeded into 6-well plates in 2 ml of complete DMEM or RPMI supplemented with 10 % FBS for overnight (300 000 cells per well). The next day medium was replaced with glutamine-containing or glutamine-free medium. Cells were incubated for additional 24 h, after that glutamine-free medium in rescue condition was replaced by the medium containing 10% of dialyzed FBS and 2 mM of dimethyl-alpha-ketoglutarate (α-KG) (Sigma). Cells were incubated for additional 24 h and used for clonogenic survival assay.

### Analysis of the oxidative stress

For the reactive oxygen species (ROS) analysis, cells were cultured in either glutamine-containing or glutamine-free medium for 30 min, 24 h, 48 h and 72 h. At the appropriate time points, cells were harvested with Accutase (StemCell Technologies), washed in PBS, and re-suspended in flow cytometry buffer (PBS without Ca^2+^ and Mg^2+^, 2mM EDTA, 3% FBS, 25 mM HEPES). 150 000 cells in 100 μL of flow cytometry buffer were stained with 2’,7’-dichlorodihydrofluorescein diacetate (H2DCFDA) (Sigma-Aldrich) for 15 min at 37°C, then washed once and re-suspended in flow cytometry buffer. The exclusion of dead cells was done by adding 7-aminoactinomycin D (7-AAD) (Sigma-Aldrich) to the cell suspension to the final concentration of 1 μg/mL. Measurement of the intensity of green fluorescence in cells has been done on flow cytometer Canto II (BD Biosciences). Analysis of flow cytometry data has been made with FlowJo software (version 10.6.2).

Measurement of reduced to oxidized glutathione ratio (GSH/GSSG) was performed with luciferase-based GSH/GSSG-Glo assay (Promega) according to the manufacturer’s instructions. In brief, cells were trypsinized, washed once with PBS, and then centrifuged at 2000 rpm for 5 min. 3000 cells were lyzed in 50 μL of passive lysis buffer for 5 min, then mixed with 50 μL of luciferin generation reagent for 30 min on a shaker plate; 100 μL of luciferase detection reagent was added to the mixture and samples were incubated for 15 min at room temperature. Samples were then transferred to the white opaque 96-well plates, and luminescence signal was read at TECAN microplate reader with the integration time of 1 sec.

### Cell death analysis

Cell death has been evaluated by staining the cells with eBioscience^™^ Annexin V Apoptosis detection kit (Thermo Fisher Scientific) according to the manufacturer’s instructions. Briefly, all cells were harvested using Accutase, washed once with PBS, and once – with 1X Annexin V buffer. 150 000 cells were stained in 200 μL of 1X Annexin V buffer for 15 min at room temperature with FITC-conjugated Annexin V, washed once with 1X Annexin V buffer and re-suspended in 200 μL Annexin V buffer. 5 μL of propidium iodide, PI (Thermo Fisher Scientific) was added to the cell suspension, and cell fluorescence was measured on flow cytometer Canto II (BD Biosciences). Analysis of flow cytometry data was performed using FlowJo software (version 10.6.2).

### Sphere forming assay

For the Gln starvation experiment, cells were seeded in complete DMEM or RPMI-1640 medium supplemented with 10 % fetal bovine serum and 2 mM L-glutamine and allowed to attach overnight. Next day cells were washed with PBS and medium was changed to glutamine-containing or glutamine-free. Cells were kept in a humidified CO_2_-containing incubator at 37°C for 72 hours. Then cells were harvested with Accutase, washed in PBS and re-suspended in MEBM (Lonza) supplemented with 4 μg/ml insulin (Sigma), B27 (Invitrogen), 20 ng/mL epidermal growth factor (EGF) (Peprotech), 20 ng/ml basic fibroblast growth factor (FGF) (Peprotech), 1mM L-glutamine and 1% penicillin-streptomycin solution. 1000 viable cells were seeded into 1 well of 24-well ULA plate (Corning Inc.) in 1 mL of MEBM. The next day 1 plate was irradiated with 4 Gy of X-rays, and the control plate was sham-irradiated. One week after, 1 mL of fresh MEBM was added to all wells. Spheres were scanned at Celigo S Imaging Cell Cytometer (Brooks) and counted using FIJI software 2 weeks after the seeding. Spheres were intensively re-suspended before scanning to dissociate cell aggregates. Cell clumps were excluded from the analysis based on their shape, size, and structure.

### ALDEFLUOR assay

Measurement of aldehyde dehydrogenase activity was performed using Aldefluor kit (Stem Cell Technology) according to the manufacturer’s instructions. In brief, cells were harvested with Accutase, washed once with PBS, and re-suspended in Aldefluor buffer (provided within the kit), supplemented with 5% FBS and 1 mM EDTA. 150000 of cells were stained with Aldefluor reagent at dilution of 1:200 in 100 μL of Aldefluor buffer at 37°C in the dark. ALDH inhibitor N,N-diethylaminobenzaldehyde (DEAB) at dilution of 1:50 was added to the control tubes together with the same amount of Aldefluor reagent used for staining of samples, and control cells were stained under the same conditions. After the staining cells were washed once with Aldefluor buffer and measured by flow cytometry on Canto II (BD Biosciences). The exclusion of dead cells was done by adding 7AAD (1 µg/mL) to the cells prior to measurement.

### Fluorescence microscopy of cell cultures

For the fluorescence microscopy analysis, cells were seeded on EZ chamber slides (Merck Millipore) overnight. Next day media was changed to glutamine-containing or glutamine-free. Cells were kept in a humidified CO_2_-containing incubator at 37°C for 24 h, then irradiated with 4Gy of X-rays or sham-irradiated. 24 h after irradiation, cells were fixed 30 min at 37°C with 4% formaldehyde in PBS, permeabilized with 1% Triton X-100 in PBS (7 min at room temperature), washed thoroughly with PBS, blocked with 10% bovine serum albumin (BSA) 30 min at 37°C and incubated with primary anti-γH2A.X mouse antibodies (Merck Millipore) or rabbit anti-LC3B antibodies (Cell Signaling Technology) diluted as 1:400 or 1:200, correspondingly, for overnight at 4°C. Next day cells were washed 10 times with PBS and incubated with secondary AlexaFluor^™^ 488 goat anti-mouse or anti-rabbit antibodies (Invitrogen) (diluted 1:500) combined with 4′,6-diamidino-2-phenylindole(DAPI) at a final concentration of 1 μg/mL for 1 hour at 37°C in the dark. After washing with PBS, a few drops of Mowiol mounting solution (Thermo Fisher Scientific) have been placed on top of cells, and then slides were covered with glass coverslips. The immunofluorescence staining was analyzed at Axioimager M1 microscope (Zeiss). The number of foci, number of autophagosomes, and the size of the cellular nuclei were analyzed by ImageJ software. The numbers of foci were normalized to the size of the cell nucleus.

### Fluorescence microscopy of prostate tissues

Clinical material was collected with informed consent and approval from the local ethics committee (Institutional Review Board of the Faculty of Medicine, Technische Universität Dresden, EK152052013). Primary prostate cancer and adjacent normal tissues (benign hyperplasia, BPH) were pathologically evaluated upon radical prostatectomy, cut into 5 mm^³^ pieces and placed for 24 h in DMEM/F12 media containing 1 % penicillin/streptomycin and 1 % HEPES buffer in 24-well ULA plates (Corning Inc.) for reoxygenation. Then tissues were irradiated with 4 Gy of X-rays or sham-irradiated and kept in a humidified incubator at 37°C with 5% CO_2_ for 24 h with or without glutamine deprivation. Next, tissues were fixed with 4 % paraformaldehyde for 4 h and conserved in 30 % sucrose in phosphate-buffered saline (PBS) before embedding in TissueTek compound reagent. 10 µm tissue sections were blocked in 5 % horse serum in PBS overnight. The biopsies were stained with anti-ALDH1A1 (1:50). Hematoxylin eosin (HE) staining and co-staining with anti-α-Methylacyl-CoA racemase (AMACR, 1:100) and anti-CK5 (1:100) was used as control to discriminate normal from malignant glands. After washing, slides were stained with secondary AlexaFluor^™^ 488 goat anti-mouse or anti-rabbit antibodies (Invitrogen) (diluted 1:500) combined with 4′,6-diamidino-2-phenylindole (DAPI) at a final concentration of 1 μg/mL. After washing, the slides were mounted using Mowiol mounting solution (Thermo Fisher Scientific). The slides were scanned with the slide scanner Axio Scan.Z1 (Zeiss). Mean pixel intensity, mean area and cellular positivity were measured using ImageJ software. Each treatment group was evaluated in five patients as 3 biopsies per patient and two fields of view per biopsy.

### RT-qPCR

Total RNA was isolated by using the RNeasy Kit (Qiagen), and cDNA synthesis was performed using SuperScript II Reverse Transcriptase (Thermo Fisher Scientific) according to the manufacturer’s protocol. SYBR Green-based real-time quantitative PCR was performed in the StepOnePlus^™^ Real-Time PCR System (Applied Biosystems). Primers used in this study are listed in **Supplementary Table 5**.

### Western blotting

Cells were harvested by scraping into RIPA lysis buffer on ice. Lysates were centrifuged at 13000 rpm for 10 min, and the supernatant was transferred to the new tubes. Measurement of the total protein concentration was done by using BCA assay kit (Pierce) in 96-well plate format. Samples were mixed with 4x Laemmli Sample Buffer (Bio-Rad) containing dithiothreitol (DTT) at a final concentration of 10 mM. Samples were denaturated at 96°C for 3 min, and 10 to 30 μg of total protein was loaded onto NuPAGE Bis-Tris 4-12% acrylamide gels (Thermo Fischer Scientific) along with prestained protein ladder PageRuler (10-180 kDa) (Thermo Fisher Scientific) and run at 100 V. Proteins were transferred to the Protran® nitrocellulose membrane (Whatman) for 1.5 h at 100 V with cooling. The transfer quality was confirmed with Ponceau S solution (Sigma). Membranes were blocked in 5% BSA in PBS for 1 h at room temperature and incubated with primary antibodies overnight on the rocker shaker at 4°C. Primary antibodies were used at concentrations recommended by the manufacturer. Next day membranes were washed 3 times for 10 minutes in 0.05% PBS/Triton X-100 solution, and corresponding secondary antibodies conjugated with horseradish peroxidase (GE Healthcare) were added. Membranes were incubated with secondary antibodies 1.5 hours on the shaker at room temperature, then washed 3 times with PBS/Triton X-100 solution, covered with freshly prepared SuperSignal West Dura Extended Duration Substrate (Thermo Fischer Scientific), wrapped in Saran wrap and exposed to the LucentBlue X-ray film (Advansta Inc.). The optical density (OD) of the bands on the film was quantified using ImageJ software, and OD signals were normalized to housekeeping protein levels in each lane. For quantification of the western blot results for H3K27me3, H3 signal was used for normalization. Antibodies used for Western blot analysis are listed in **Supplementary Table 5**.

### siRNA-mediated gene expression knockdown

For knockdown of *MYC, GLS*, and *ATG5* gene expression, cells were transfected with RNAiMAX (Life Technologies GmbH) according to the manufacturer’s protocol with siRNA duplexes at the final concentration of 40 pM. The RNA duplexes were synthesized by Eurofins. Gene expression knockdown was validated by Western blot analysis with specific antibodies. The siRNA sequences are listed in **Supplementary Table 5**.

### Mass spectrometry-based analysis of Krebs cycle metabolites

Cells were seeded into 6-well plates in 2 ml of culture medium for overnight (500.000 cells per well). The next day cells were washed three times with ice cold PBS and directly scratched in 500 µl of methanol. Using liquid chromatography-mass spectrometry, eight organic acids of the Krebs cycle (succinate, fumarate, malate, citrate, isocitrate, cis-aconitate, α-ketoglutarate, 2-hydroxyglutarate) as well as pyruvate, lactate and four amino acids (glutamate, glutamine, aspartate, and asparagine) were measured in DU145, DU145 RR, LNCaP, LNCaP RR as well as primary prostate cancer and BPH cell cultures. Metabolites were analyzed as previously described ^73^. In brief, metabolites were extracted from cells with cold methanol. Supernatants derived after centrifugation of cell extracts, were dried, re-suspended in mobile phase, and further cleaned with a 0.2 µm centrifugal filter before injection. For chromatographic separation of metabolites, the elution gradient applied was as follows: 99% A (0.2% formic acid in water), 1% B (0.2% formic acid in acetonitrile) for 2.00 min, then 100% B at 2.50 to 2.65 min, 1% B at 3.40 min and equilibration with 1% B until 5.00 min. Quantification of Krebs cycle metabolites was done by comparisons of ratios of analyte peak areas to peak areas obtained from stable isotope labeled internal standards obtained in samples to those peak area ratios in calibrators. Results were normalized to cell number.

### Targeted metabolomics of amino acids and biogenic amines

Cells growing at 10 cm dished were briefly washed in ice-cold PBS and scratched directly into methanol. For each sample, cell number was determined by trypsinization and counting of the cells from the control dish. The numbers of harvested cells were ≥ 1.8 × 10^6^ cells/sample. The samples were shipped to Biocrates Life Sciences AG at dry ice and stored at −80°C after the reception. Biocrates’ commercially available Kit plates were used for the quantification of amino acids. The fully automated assay was based on PITC (phenylisothiocyanate) derivatization in the presence of internal standards followed by FIA-MS/MS (acylcarnitines, (lyso-) phosphatidylcholines, sphingomyelins, hexoses) and LC-MS/MS (amino acids, biogenic amines) using a SCIEX 4000 QTRAP (SCIEX) or Waters Xevo TQ-S (Waters) instrument with electrospray ionization (ESI). Accuracy of the measurements (determined with the accuracy of the calibrators) was in the normal range of the methods (deviations from target ≤ 20 %). Quality control samples were within the pre-defined tolerances of the method. The raw data (μM) were recalculated and normalized to nmol/10^6^ cells values (**Supplementary Table 6**). The data were further cleaned, applying a modified 80% rule for statistical analysis, meaning that at least 80% valid values above LOD need to be available per analyte in the samples for each group. If at least one group, in this case, the individual cell lines LNCaP, PC3 and DU145, did fulfill this criterion, the analyte was included for further statistical analysis.

### ^13^C-labeled glutamine tracing and mass spectrometry analysis

Cells were seeded into 6-well plates in 2 ml of culture medium for overnight (500.000 cells per well). The next day cells were washed three times with PBS and cultured with DMEM (DU145 cells) or RPMI (LNCaP cells) medium supplemented with 2 mM [U-^13^C_5_] L-glutamine (Sigma) for 10, 30 and 90min. Three independent experiments were performed with one replicate. Targeted analysis of metabolites was performed by mass spectrometry using an ABSCIEX 5500 QTRAP equipped with SelexION for differential mobility separation (DMS) and acquired using multiple reaction monitoring (MRM) in negative mode. Cells were quenched with an ice-cold solution containing 20% methanol, 0.1% formic acid, 10 µM D4-taurine, 3mM NaF, 1mM phenylalanine and 100 µM EDTA. Cell extracts were then collected, frozen immediately, lyophilized overnight and resuspended in water before mass spectrometry analysis. Samples were injected onto a Hypercarb column (3 μm particle size, 4.6 × 100 mm, Thermo Fisher Scientific) at a flow rate of 0.6 mL/min on an ACQUITY UPLC System. Chromatographic separation was achieved with a gradient of ionic strength: mobile phase A contained 15 mM ammonium formate and 20 μL EDTA; mobile phase B contained 15 mM ammonium formate and a acetonitrile: isopropyl alcohol mixture (2:1). The gradient had a duration of 6min with the following profile: 0min, 0% B; 1 min, 0% B; 1.5 min, 20% B; 2 min, 20% B; 2.5 min, 0% B; 6 min, 0% B. Samples were ionized by electrospray into the mass spectrometer with the following source parameters: CUR: 20, CAD: 7, IS: −4500,TEM: 550, GS1: 70 and GS2: 60. DMS was used as an additional separation axis (DT: low, MD: 2-propanol, MDC: high, DMO: 3 and DR: off). DMS-related Separation Voltage (SV) and Compensation Voltage (CoV) for each metabolite was optimized before each experiment. Peak identification and chromatographic separation of all metabolites was confirmed with known standards. Peaks were integrated with ElMaven v0.6.1 (Elucidata). The ensuing peak areas were used for the calculation of metabolite concentrations and ^13^C-enrichments as previously described ^74^.

### Flux calculation

Glutamine and mitochondrial metabolism were determined by fitting a one compartment metabolic model to the labeling time courses of glutamine, glutamate, malate, aspartate and citrate according to the model depicted in **Supplementary Figure 1 d**. No significant enrichment was detected in phosphoenolpyruvate, pyruvate and lactate and the corresponding reactions were not included in the model. Metabolite concentrations were averaged from all time points (0, 10, 30 and 90min) and kept constant in the model to respect metabolic steady-state (**Supplementary Table 7**). Flux calculations were performed with CWave software running in Matlab R2018b and using the time course of [U-^13^C_5_] glutamine as the driving function using the principles described previously ^74^. Two main features are different from the previous studies that reflect the nature of these cells and the labelling strategy. An interaction of bicarbonate with the other (de)carboxylating reactions was included to explain the generation of [M+6] citrate. In the absence of ^13^C-label in the glycolytic intermediates, only a combination of VIDHr and ^13^C-bicarbonate can generate that labeling pattern from [U-^13^C_5_] glutamine. Finally, the addition of VGDH allowed for the possibility to calculate the rate of glutamine oxidation, an important feature of cancer cells. The three time courses were modelled separately and and the final fluxes (μM/min/500K cells) were averaged at the end (i.e., 3 flux values per cell line).

### Analysis of amino acid concentrations in blood plasma

The concentrations of plasma amino acids were determined by a Biochrom 30plus Amino Analyzer (Biochrom Ltd.), a cation-exchange chromatography system^75^. In brief, amino acids were purified by mixing of 200 μl plasma with 50 μl of sulfosalicylic acid, 10% in distilled water (Sigma Aldrich Corp.). Precipitation was completed by freezing the samples at −20 °C for 10 min. After centrifugation at 10,000 rpm for 5min, 180 μl of the supernatant were mixed with 180 µl sample dilution buffer containing the internal standard norleucine, 500 μmol/L (Sigma Aldrich Corp.) The mixture was passed through syringe filter polyvinylidene difluoride (PVDF) membranes. Post-column derivatization with ninhydrin allowed the detection of amino acid at the wavelength of 570 nm. Standard lithium citrate buffers with pH 2.80, 3.00, 3.15, 3.50, and 3.55 and ninhydrin reagents utilized during separation were provided ready to use by Biochrom Ltd. 125 µM amino acid standard solution was prepared by mixing physiological basis with acids and neutrals and internal standard solution (Sigma Aldrich Corp.). Data analysis was performed using the EZChrome software (Agilent Technologies). Areas of the peaks were measured to determine amino acid concentrations, expressed in terms of μmol/L (**Supplementary Table 2**).

### Array comparative genome hybridization of single CTCs

Enrichment of CTCs was performed using the CellSearch Circulating Epithelial Cell Kit (Menarini) based on immunomagnetic ferrofluid conjugated with an EpCAM-directed antibody. 7.5 ml blood was used for enumeration of CTCs. Identification of CTCs was performed using immunofluorescent staining directed against cytokeratins, CD45 to exclude leucocytes and 4′,6-diamidino-2-phenylindole (DAPI) to confirm nucleo-morphological integrity.

Single CTCs were isolated from CellSearch cartridge by micromanipulation with the CellCelector (ALS) as described previously ^76^. Chromosomal DNA of single isolated cells was amplified by whole genome amplification with the Ampli1 WGA Kit (Menarini). Afterwards DNA integrity was determined with the Ampli1 QC Kit (Menarini).

Whole genome amplification products of highest quality were processed for aCGH as previously described ^77^. 1 μg DNA was processed. As reference, the whole genome amplification product of a single female white blood cell was used. For data analysis, the output image files were normalized and fluorescence ratios for each probe were determined using Feature Extraction software (Agilent Technologies, Santa Clara, United States; version 10.7.3.1, Protocol CGH_1105_Oct09). Data were visualized and analyzed with the Genomic Workbench 6.5.0.18 software by applying the ADM-2 algorithm with a threshold of 6.0. The centralization algorithm was set to a threshold of 4.0 with a bin size of 10. To identify copy number alterations, an aberration filter with a minimum log2 ratio of ±0.3 and a minimum of 100 consecutive probes was set.

### Seahorse analysis

Cells were seeded in triplicates in Seahorse XFp miniplates (seeding surface of 0.106 cm^2^) in RPMI (LNCaP and LNCaP RR cells) or DMEM (DU145 and DU145 RR cells) supplemented with 10% FBS. 24 h later, medium was changed to glutamine-free medium for glutamine oxidation assay, and miniplates were incubated at 37°C in a CO_2_ incubator overnight. The Seahorse XFp sensor cartridges were hydrated at 37°C with Seahorse XF calibrant in a non-CO_2_ incubator overnight. Next day cell medium was changed to substrate-free XF base medium and cells were incubated at 37°C for 1 h in a non-CO_2_ incubator. The Seahorse XF Mitostress test was then run using a 4-injection protocol: after 18 min basal condition, 2 mM glutamine was injected, followed by serial injections of 1.5 µM oligomycin, 1 µM FCCP and 0.5 µM rotenone/antimycin A. To measure glycolytic rate in basal conditions, 24 h after seeding in Seahorse XFp miniplates medium was substituted with Glycolytic rate assay medium supplemented with 1 mM pyruvate, 2 mM glutamine and 10 mM pyruvate, and cells were incubated at 37°C for 1 h in a non-CO_2_ incubator. The XFp glycolytic rate assay was then run by using serial injections of 0.5 µM rotenone/antimycin A and 50 µM 2-deoxy-D-glucose (2-DG) as final concentration. Oxygen consumption rate (OCR) and extracellular acidification rate (ECAR) were measured using a Seahorse XFp Analyzer and data were normalized to the final number of cells.

### Analysis of MYC expression in tumor tissues

Expression of MYC gene was analyzed by nanoString technology using formalin-fixed paraffin-embedded (FFPE) tumor tissues of prostate cancer patients (n=74) treated with curatively-intended, definitive radiotherapy at the Department of Radiotherapy and Radiation Oncology, University Hospital Carl Gustav Carus and Faculty of Medicine. All patients were diagnosed with intermediate- or high-risk localized prostate cancer and have a minimum follow-up of 7 years. Patient characteristics are shown in **Supplementary Table 3**. The clinical endpoint was PSA relapse-free survival. The experiments were approved by the local ethics committee (Institutional Review Board of the Faculty of Medicine, Technische Universität Dresden, EK49022015). Survival curves were estimated by the Kaplan–Meier method. To compare patient groups stratified by median MYC marker expression, Log-rank tests were employed. The analyses are conducted by SPSS 23 software (IBM Corporation, Armonk, NY, USA). Two-sided tests were performed, and p-values <0.05 were considered as statistically significant.

### Microarray analysis of the prostate cancer cell lines

Comparative gene expression profiling of the prostate cancer cells starved for glutamine for 24 h or cultured with glutamine supplementation was performed using SurePrint G3 Human Gene Expression 8×60K v3 Microarray Kit (Design ID 039494, Agilent Technologies) according to manufacturer’s recommendations as described previously ^4, 20^. Briefly, cells were cultured in either glutamine-containing or glutamine-free medium for 24 h and then used for total RNA isolation using the RNeasy kit (Qiagen). Sample preparation for analysis and processing of the arrays were performed using the protocol provided by Agilent Technologies. Agilent Feature Extraction Software was used to obtain the probe values from image files. The dataset was analyzed using SUMO software package: http://www.oncoexpress.de. Data deposition: all data is MIAME compliant. The raw data has been deposited in the Gene Expression Omnibus (GEO) database, accession no GSE148016. Gene expression analysis of the prostate cancer cells growing under monolayer and sphere-forming conditions was performed in the same way. The raw data has been deposited in the Gene Expression Omnibus (GEO) database, accession no GSE148013. Gene expression analysis of the ALDH^+^ and ALDH^-^ cells as well as parental and radioresistant prostate cancer cells was described previously ^4, 78^ (Gene Expression Omnibus (GEO) database accession no GSE134499 and GSE53902).

### Statistics

The results of amino acid analysis, Krebs cycle metabolites, colony formation assays, γH2AX foci assay, autophagosome analysis, sphere-forming assays, Seahorse metabolism analysis, flow cytometry, and Western blotting were analyzed by paired t-tests. The differences between cell survival curves were analyzed using the statistical package for the social sciences (SPSS) v23 software by fitting the data into the linear-quadratic model S(D)/S(0)=exp(αD+βD^2^) using stratified linear regression as described previously^4^. Multiple comparison analysis was performed using one-way ANOVA analysis by GraphPad Prism software. Multiple testing corrections were not applied for the data depicted in **Figure 3 a, 7 a, 7 b** and **Supplementary Figure 7 d** since the experimental treatments were non-independent. A p-value < 0.05 was regarded as statistically significant. Correlation was evaluated by SUMO software (**Figure 1 a, 2 a, 2 g, Supplementary Figures 1 a, 5 b, 6 a** and **6 b**) or by GraphPad Prism software (**Figure 6 b** and **Supplementary Table 2**) using the Pearson correlation coefficient. Principle component analysis (PCA) using JMPPro software was applied to visualize metabolic differences and determine the variation present in the radioresistant prostate cancer cell lines in comparison to their parental counterpart (DU145, PC3 and LNCaP) based on the mass spectrometry quantification of amino acids and biogenic amines (AbsoluteIDQ® p180 Kit assay, Biokrates). The correlation matrix contained 41 parameters including 14 significant variables to determine eigenvector for multivariate analysis. For correlation analysis a dataset of 490 cases for PCa from The Human Cancer Genome Atlas (TCGA) was downloaded from cBioportal: http://www.cbioportal.org/. The data were processed following the TCGA policy. Expression heat maps were generated using SUMO software package: http://www.oncoexpress.de. For Kaplan-Meier analysis in **Supplementary Figure 8 c**, a dataset for TCGA prostate adenocarcinoma (PRAD) gene expression was downloaded from UCSC Xena Functional Genomics Explorer: https://xenabrowser.net/. For the patient cohort described in supplementary Table 3, the primary endpoint was PSA recurrence, which was calculated from the first day of radiotherapy to the occurrence of PSA relapse or censoring. Corresponding survival curves were estimated by the Kaplan-Meier method and compared by log-rank tests.

## Acknowledgments

The authors thank the patients that contributed to this study.

The authors thank Dr. Ina Kurth for help with establishing the procedure for CTC analyses and Dr. Linda Hein for assistance with animal experiments. We greatly appreciate the members of the Group of Biomarkers for Individualized Radiotherapy for their critical reading of the manuscript and the team of The Core Facilities of the Technology Platform at BIOTEC TU Dresden for their constant technical support.

## Author Contributions

AM, CP and AD conceived the study. AM, UK, AL, OC, TA, SR, MR, VT, GN, BA, PM, LY, VL, AF, BB, NHS, MT, US, SZ, MR, FL, TH, CG and AD performed the experiments; SL, AL, VL, CS, AM, UK, CP, AD analyzed the data; AM, HN, AL, IIS, SS, GE, AA, TA, LKS, GBB, MB, MK, CP and AD supervised the experiments and edited the manuscript; AD supervised the study, provided project administration and wrote the manuscript; all the authors reviewed the final version.

## Ethics Approval and Consent to Participate

The experiments were approved by the local ethics committee (Institutional Review Board of the Faculty of Medicine, Technische Universität Dresden, EK49022015). Study was conducted in accordance with the Declaration of Helsinki.

## Data availability

All data supporting the findings of this study and unique biological materials used in this study are available from the corresponding authors upon request. Gene expression data generated during the study available in the public repository Gene Expression Omnibus (GEO), accession no GSE148013, GSE134499, GSE53902 and GSE148016.

## Standard abbreviations

ALDH: aldehyde dehydrogenase
ALKBH: Alkylated DNA Repair Protein AlkB Homolog family
α-KG: alpha-ketoglutarate
ATG5: autophagy related 5
BPH: benign prostate hyperplasia
CSC: cancer stem cell
DDR: DNA damage response
Gln: glutamine
GLS: glutaminase
Glu: glutamate
GSH: reduced glutathione
GSSG: oxidized glutathione
LC3B: Microtubule-associated proteins 1 light chain 3B
OCR: oxygen consumption rate
PCa: prostate cancer
PDPC: patient-derived primary cell cultures
PSA: prostate-specific antigen
ROS: reactive oxygen species
RR: radioresinance
TCA cycle: tricarboxylic acid cycle
TCGA: The Cancer Genome Atlas
OXPHOS: oxidative phosphorylation

