## Supplementary material for "GLS-driven glutamine catabolism contributes to prostate cancer radiosensitivity by regulating the redox state, stemness and ATG5-mediated autophagy": Figures S1-S8, Tables S1-S7

**Supplementary information**

**Supplementary Table 1.** Characteristics of PCa patients whose blood plasma samples were analyzed for amino acid metabolites (n = 20).

**Supplementary Table 2.** Analysis of blood plasma metabolites and PSA-DT in PCa patients before and after radiotherapy.

**Supplementary Table 3.** Characteristics of PCa patients treated with curatively-intended, definitive radiotherapy at the Department of Radiotherapy and Radiation Oncology, University Hospital Carl Gustav Carus and Faculty of Medicine (n = 74). Formalin-fixed paraffin-embedded (FFPE) tumor tissues from these patients were used for analysis of *MYC* gene expression by nanoString technology.

**Supplementary Table 4.** Characteristics of PCa patients (n = 14). Primary prostate cancer and adjacent normal tissues (benign hyperplasia, BPH) from these patients were used for primary tissue cultures and radiobiological colony forming assays.

**Supplementary Table 5.** Primers, siRNA oligos and antibodies used in the study.

**Supplementary Table 6.** Targeted metabolomics of amino acids and biogenic amines.

**Supplementary Table 7.** Concentration of metabolites used in for metabolic flux analysis. Concentrations are reported as µM/500k cells (Average ± Standard Error).

**Supplementary Figure 1. Analysis of Gln metabolism in DU145 RR and LNCaP RR cells.** **a** Gene expression analysis in parental and RR PCa cells showed similarity in deregulation pattern of genes involved in the control of TCA. **b** Absolute log2 fold change of genes involved in TCA cycle regulation. **c** Principal component analysis (PCA) to visualize metabolic differences and the relationship of the analyzed parental and radioresistant (RR) cells. The PCA was performed using the changes in metabolic profiles of amino acids and biogenic amines in parental and RR cells. The differences between analyzed cells are driven e.g., by altered levels of the glucogenic amino acids, including Gln and Glu. The variance explained by PC1 = 52.6% and PC2 = 17.3%. **d** Schematic of the model used to quantify the metabolic fluxes in DU145 and LNCaP cell lines. [U-$^{13}C_5$]Glutamine was used as the driving function. The quantified reactions were: glutaminase ($V_{Glnase}$), glutamate consumption other than the TCA cycle ($V_{GluOut}$), glutamate dilution from sources other than the TCA cycle ($V_{GluDil}$), aspartate transaminase ($V_X$), glutamate dehydrogenase ($V_{GDH}$), oxidative and reductive directions of isocitrate dehydrogenase ($V_{IDH}$ and $V_{IDHr}$, respectively), the lower portion of the TCA cycle ($V_{TCA}$), the scrambling of label due to the equilibrium with fumarate ($V_{SC}$), citrate synthase ($V_{CS}$) and ATP-dependent citrate lyase ($V_{Acly}$). **e** Metabolic fluxes analysed in LNCaP and DU145 parental and RR cells. Data are mean ± s.d. **f** Glutaminase / TCA cycle ratio based on the metabolic flux data. Data are mean ± s.d. *$p < 0.05$; ***$p < 0.001$.

**Supplementary Figure 2. Radiobiological pathways associated with Gln deprivation**. **a** Western blot analysis of DDR regulations in response to Gln depletion for 72 h and quantification of the Western blot data. Cells grown in presence of glutamine were used as control. Cells were cultured in swapped medium: LNCaP cells were cultured in DMEM, and DU145 cells were cultured in RPMI. The results showed no impact of culture medium on the observed effect as compared to **Figure 2 b**. Data are mean ± s.d. **b** Ingenuity pathway analysis revealed downregulation of ATM signaling network in DU145 cells in response to Gln starvation. **c** Gln starvation lowered intracellular GSH/GSSG ratio in PCa cells. Data are mean ± s.d. **d** Gln depletion in DU145 P and RR cells upregulated the hallmark genes of endoplasmic reticulum (ER) stress response. **e** qPCR analysis of ALKBH1 and ALKBH3 expression in PCa cells with or without Gln starvation for 24 h. Data are mean ± s.d. **f** Plating efficacy of PCa cells incubated with or without Gln depletion for 24 h. Data are mean ± s.d. **g** Radiobiological colony-forming assay and plating efficacy of PCa cells grown with or without Gln supplementation in swapped medium: LNCaP cells were cultured in DMEM, and DU145 cells were cultured in RPMI. The effect of Gln starvation can be seen with a greater extent for DU145 cells independent on a culture medium similar to **Figure 2 e**. Data are mean ± s.d. **h** The radiosensitizing effect of Gln starvation can be partially rescued in DU145 cells by supplementation of α-KG at final concentration of 2 mM for 24 h. Data are mean ± s.d. $*p < 0.05$; $**p < 0.01$.

**Supplementary Figure 3. GLS and MYC expression levels in parental and radioresistant PCa cells and plating efficiency after GLS and MYC knockdown.** **a** Quantitative real-time PCR (RT-qPCR) analysis of MYC and GLS expression in DU145, DU145 RR, LNCaP and LNCaP RR cell lines. Data are mean ± s.d. **b** Plating efficiency of PCa cells in response to siRNA-mediated knockdown of *MYC* or *GLS* gene expression with or without Gln supplementation for 24 h. **c** Quantitative real-time PCR (RT-qPCR) analysis of GLS and SLC1A5 expression after MYC knockdown in DU145, DU145 RR, LNCaP and LNCaP RR cells. Data are mean ± s.d. $*p < 0.05$.

**Supplementary Figure 4. Chemical inhibition of GLS and MYC in PCa cells. a** Radiobiological colony-forming assay of PCa cells with or without inhibition of MYC. Cells were pretreated with MYC inhibitor 10058-F4 at $LD_{50}$ concentrations for 48 h directly prior clonogenic analysis. Cells were irradiated with indicated X-ray doses directly upon plating. **b** Plating efficiency of PCa cells in response to chemical inhibition of MYC, GLS or DMSO treatment. **c** DU145, 22Rv1, PC3 and LNCaP were treated for 48 h with different doses of MYC inhibitor 10058-F4 or with GLS inhibitor

CB-839 at concentrations ranging from 100 µM to 0 µM and counted with CASY cell counter. $LD_{50}$ values (50% lethal dose) were determined for each cell line individually by non-linear regression analysis using GraphPad Prism software (San Diego, USA). **d** Cells were incubated overnight with $LD_{50}$ concentrations of MYC inhibitor 10058-F4. The levels of Glu production were measured to confirm GLS enzymatic activity inhibition. Cells treated with DMSO were used as control. Data are mean ± s.d. **e** Plating efficiency of patient-derived primary cultures (PDPC) in response to chemical inhibition of MYC and GLS or DMSO treatment. Data are mean ± s.e.m. *p < 0.05; **p < 0.01, ***p < 0.001, ****p < 0.0001.

**Supplementary Figure 5. Mechanisms of PCa radiosensitization in response to Gln starvation. a** The growth-inhibiting effect of Gln deprivation analyzed by MTT assay. Data are mean ± s.d. **b** Comparative analysis of gene expression in LNCaP and DU145 parental (P) and radioresistant (RR) cells grown with or without Gln supplementation for 24 h. Gln deprivation results in consistent deregulation pattern of genes involved in the apoptotic mechanisms. **c** Absolute log2 fold change of genes involved in apoptosis regulation signature. **d** Plating efficiency analyzed after autophagy inhibition in Gln starved cells. Cells were cultured in medium with or without Gln for 24h. For autophagy inhibitions, cells were then treated with CQ at a concentration of 10 µM for 2 h. **e** Plating efficiency of PCa cells in response to *ATG5* knockdown. Cells were transfected with two *ATG5* siRNA or scrambled siRNA for 24 h and then supplemented with fresh media without glutamine for additional 24 h. Data are mean ± s.d. *p < 0.05; **p < 0.01; n.s. – p-value > 0.05.

**Supplementary Figure 6. Gln metabolism as a regulator of CSC-related gene expression. a** Gene expression profiling of PCa cells replenished or depleted for Gln for 24 h revealed that Gln starvation induced consistent deregulation pattern of genes involved in the CSC maintenance. **b** Western blot analysis of LNCaP and

DU145 cells treated with DZNep at a concentration of 1 µmol/L for 72 hours. **c** PCa cells growing under sphere-forming conditions have deregulation of genes involved in amino acid metabolism.

**Supplementary Figure 7. Gln metabolism regulates CSC phenotype and properties.** **a** Gln starvation decreased the total number of formed tumor spheres. Data are mean ± s.d. **b** Fluorescence microscopy analysis of ALDH1A1 expression in patient-derived PCa and BPH tissues. MFI - Mean fluorescence intensity. Data are mean ± s.d. **c** Supplementation of DU145 cells with a MYC inhibitor 10058-F4 at a concentration of 10µM for 48 h significantly lowered radiation-induced $ALDH^+$ cell subset. Data are mean ± s.d. **d** Analysis of tumor take in NMRI (nu/nu) xenograft models. PCa cells were pre-incubated in the presence or absence of Gln for 72 h, irradiated with 6 Gy of X-rays or sham irradiated, and injected into the mice in Matrigel as $5\times10^3$ viable cells/implant. *p < 0.05; **p < 0.01; n.s. – p-value > 0.05.

**Supplementary Figure 8. Analysis of Gln metabolism as clinically relevant biomarker.** **a** Gene expression of MYC in parental and radioresistant (RR) PCa cells. **b** Enrichment of MYC target gene signature in DU145 Gln replenished cells (DU145 Gln+) as compared to DU145 Gln depleted cells (DU145 Gln-); NES – normalized enrichment score; FDR – false discovery rate; p value was determined by random permutation test. **c** Analysis of The Cancer Genome Atlas (TCGA) gene expression dataset revealed that high combined expression of MYC and GLS genes is significantly associated with a decrease in relapse-free survival only in PCa patients treated with radiotherapy (n = 56). Median was determined for entire dataset before extraction of ‘no RT’ and ‘RT’ groups. Combined expression for GLS + MYC was determined as mean of normalized expression values.

Supplementary Table 1. Characteristics of PCa patients whose blood plasma samples were analyzed for amino acid metabolites

| **Study ID** | 1st PSA pre RT | 2nd PSA pre RT | **PSA-DT pre RT** | 1st PSA after RT | 2nd PSA after RT | **PSA-DT post RT** | TNM | Gleason | Therapy |
|---|---|---|---|---|---|---|---|---|---|
| **HT1** | 0.19 | 2.57 | 1.63 | 0.64 | 0.10 | -5.05 | pT3,pN1 | 4+4 | ADT + local RT |
| **HT2** | 21.10 | 126.00 | 12.06 | | | | cT3 cN1 cM1 | 5+4 | ADT + local RT |
| **HT3** | 10.30 | 18.10 | 1.72 | 13.80 | 50.30 | 1.48 | pT3, pN0 | 4+5 | ADT + local RT |
| **HT4** | 50.30 | 67.40 | 0.95 | 48.20 | 81.30 | 0.31 | cT4, cN1, cM1 | 5+5 | ADT + local RT |
| **HT6** | 2.47 | 4.40 | 4.84 | 0.90 | 3.5 | 8.93 | pT2,pN0,pM0 | 3+4 | ADT + local RT |
| **HT7** | 0.18 | 4.00 | 1.34 | | | | pT3,pN1,cM0 | 4+4 | ADT + local RT |
| **HT8** | 12.40 | 24.40 | 6.59 | 0.80 | 1 | 73.72 | pT2,pN0,pM0 | 3+3 | ADT + local RT |
| **HT9** | 1.6 | 2.20 | 4.79 | 0.06 | 0.13 | 7.14 | pT2,pN0,pM0 | | ADT + local RT |
| **HT10** | 5.90 | 7.10 | 8.36 | 0.10 | 0.11 | 78.54 | pT3,pN1,cM0 | | ADT + local RT |
| **HT11** | 1.00 | 8.00 | 2.57 | 21.40 | 43.20 | 1.05 | pT3,pN1 | 4+4 | ADT + local RT |
| **HT12** | 5.70 | 13.00 | 1.71 | 2.2 | 11.00 | 3.27 | M1 | | ADT + local RT |
| **HT13** | 0.9 | 1.50 | 8.91 | 3.10 | 3.15 | 31.77 | pT1,pN0 | 3+3 | ADT + local RT |
| **HT15** | 0.17 | 2.30 | 4.10 | 0.10 | 0.01 | -2.08 | pT3a, pN0 | 4+4 | ADT + local RT |
| **HT16** | 2.00 | 4.26 | 3.09 | 0.05 | 13.00 | 1.18 | pT2,pN1,Mo | 3+4 | ADT + local RT |
| **HT17** | 2.80 | 4.75 | 3.98 | 0.06 | 0.70 | 1.75 | pT3b, pN0, cM0 | 4+5 | ADT + local RT |
| **HT18** | 53.00 | 60.00 | 4.84 | 75.00 | 78.00 | 105.45 | pT2c,pN0 | 3+4 | ADT + local RT |
| **HT21** | 66.00 | 168.00 | 3.04 | 90.00 | 128.00 | 7.35 | M1 | 4+5 | ADT + local RT |
| **HT22** | 0.03 | 0.10 | 6.37 | 0.05 | | | pT3b,pN1,M0 | 5+4 | ADT + local RT |
| **HT23** | 3.50 | 4.30 | 20.43 | | | | cT2,cN0,cM0 | 3+4 | ADT + local RT |
| **HT24** | 0.17 | 0.55 | 1.79 | | | | cT3b,cN0,cM0 | 4+3 | ADT + local RT |

- Data are not available

Supplementary Table 2. Analysis of blood plasma metabolites and PSA-DT in PCa patients before and after radiotherapy, μmol/L

| | 1 | 2 | 3 | 4 | 5 | 6 |
|---|---|---|---|---|---|---|
| **Study ID** | **1-methyl-histidine** | **3-methyl-histidine** | **alpha aminoadipic acid** | **alpha aminobutyric acid** | **alanine** | **arginine** |
| HT1 | 5.7 | 0 | 2.7 | 7.3 | 147.3 | 34.1 |
| HT2 | 0 | 2.4 | 3.4 | 10.8 | 198.9 | 19.7 |
| HT3 | 5.6 | 0 | 3.4 | 6.5 | 130.1 | 14.3 |
| HT4 | 0 | 0 | 0 | 2.1 | 56 | 6.6 |
| HT6 | 30 | 5.7 | 5.3 | 14 | 160.6 | 26.1 |
| HT7 | 6.7 | 0.9 | 1.4 | 4.5 | 105.1 | 29.3 |
| HT8 | 3.7 | 0 | 4.5 | 3.7 | 160.4 | 19.3 |
| HT9 | 3.7 | 0 | 3.6 | 3.3 | 151.2 | 15.9 |
| HT10 | 4.4 | 1.3 | 2.2 | 6.3 | 184.2 | 16.1 |
| HT11 | 5.1 | 0 | 0 | 3.5 | 108.1 | 14.2 |
| HT12 | 0 | 1.4 | 0 | 6.9 | 80.7 | 11.6 |
| HT13 | 7.2 | 12.7 | 0.8 | 10 | 120.7 | 19.6 |
| HT15 | 5.4 | 5.2 | 0.8 | 6.8 | 94.3 | 14.9 |
| HT16 | 1.6 | 3.3 | 1 | 5.8 | 154.4 | 28 |
| HT17 | 1.9 | 0 | 2.1 | 4.6 | 161.5 | 12.8 |
| HT18 | 3.6 | 0 | 1.9 | 2.1 | 120.4 | 12 |
| HT21 | 0 | 0 | 0 | 5.4 | 78.5 | 19.3 |
| HT22 | 0 | 0 | 5.3 | 7.7 | 193.1 | 6.4 |
| HT23 | 7.1 | 0 | 0 | 2.9 | 119.5 | 5.3 |
| HT24 | 3.8 | 0 | 0.9 | 7 | 135.9 | 22.8 |

**Correlation with PSA DT before RT**

| | | | | | | |
|---|---|---|---|---|---|---|
| **Pearson r** | 0.07208 | 0.1667 | 0.05045 | 0.06993 | 0.3414 | -0.3386 |
| **95% confidence interval** | -0.3827 to 0.4988 | -0.2979 to 0.5674 | -0.4011 to 0.4823 | -0.3846 to 0.4972 | -0.1193 to 0.6811 | -0.6794 to 0.1224 |
| **R square** | 0.005196 | 0.02777 | 0.002545 | 0.004891 | 0.1165 | 0.1146 |
| **P value (two-tailed)** | 0.7627 | 0.4825 | 0.8327 | 0.7695 | 0.1408 | 0.1443 |
| **P value summary** | ns | ns | ns | ns | ns | ns |
| **Significant? (alpha = 0.05)** | No | No | No | No | No | No |
| **Number of XY Pairs** | 20 | 20 | 20 | 20 | 20 | 20 |

**Correlation with PSA DT after RT**

| | | | | | | |
|---|---|---|---|---|---|---|
| **Pearson r** | -0.03787 | -0.03828 | 0.2392 | -0.2338 | 0.3246 | -0.1612 |
| **95% confidence interval** | -0.5398 to 0.4839 | -0.5401 to 0.4836 | -0.3113 to 0.6695 | -0.6664 to 0.3165 | -0.2252 to 0.7176 | -0.6222 to 0.3828 |
| **R square** | 0.001434 | 0.001466 | 0.05722 | 0.05467 | 0.1054 | 0.02599 |
| **P value (two-tailed)** | 0.8934 | 0.8923 | 0.3906 | 0.4016 | 0.2379 | 0.566 |
| **P value summary** | ns | ns | ns | ns | ns | ns |
| **Significant? (alpha = 0.05)** | No | No | No | No | No | No |
| **Number of XY Pairs** | 15 | 15 | 15 | 15 | 15 | 15 |

(continued)

| 7 | 8 | 9 | 10 | 11 | 12 | 13 | 14 | 15 |
|---|---|---|---|---|---|---|---|---|
| **asparagine** | **aspartic acid** | **beta-aminoisobutyric acid** | **carnosine** | **citrulline** | **cystine** | **cystathionine** | **gamma-aminobutyric acid** | **glutamine** |
| 16.5 | 5 | 0 | 0 | 8.7 | 0 | 0.4 | 0 | 246.1 |
| 23 | 4 | 0 | 0 | 7.2 | 2 | 0.5 | 10.6 | 117.8 |
| 11 | 4 | 0 | 0 | 6.6 | 0 | 0 | 0 | 163.7 |
| 5.8 | 5.2 | 4.4 | 0 | 4.2 | 2.4 | 0 | 0 | 72.3 |
| 18.2 | 4.5 | 0 | 0 | 16.6 | 1.3 | 0 | 5.7 | 203.9 |
| 8.1 | 3 | 3.1 | 0 | 7.2 | 0.6 | 0 | 4.5 | 173.4 |
| 9.4 | 3.6 | 2.8 | 0 | 4 | 0 | 0 | 5.2 | 131.7 |
| 9.7 | 2.2 | 0 | 0 | 9.9 | 1.9 | 0.5 | 1.7 | 121.3 |
| 9.6 | 3.7 | 0 | 0 | 11.9 | 5.3 | 0 | 13 | 160.7 |
| 11.8 | 3.5 | 13.4 | 0 | 5.6 | 0 | 0 | 0 | 163.4 |
| 10.3 | 4.3 | 0 | 0 | 5.3 | 1.4 | 0 | 9.1 | 212.7 |
| 11.1 | 5.9 | 0 | 0 | 10 | 1.8 | 0 | 3.4 | 177 |
| 9.3 | 2.5 | 0 | 0 | 6.8 | 2.4 | 0 | 0 | 158.8 |
| 14.3 | 2.4 | 0 | 0 | 15.8 | 7.7 | 0 | 6.7 | 202.5 |
| 10.8 | 5.8 | 0 | 0 | 8.2 | 0 | 0 | 0 | 180.6 |
| 11.3 | 3 | 8.1 | 0 | 12.6 | 1.1 | 0 | 0 | 182.8 |
| 0 | 5.2 | 1.6 | 5.8 | 1.8 | 0 | 119.3 | 52.5 | 59.8 |
| 8.7 | 3.9 | 0 | 1 | 6.4 | 3.6 | 0 | 0 | 204 |
| 2.8 | 7.2 | 0 | 0 | 3.2 | 6.3 | 0 | 142.4 | 26.1 |
| 15.4 | 5.8 | 0 | 0 | 3.8 | 0 | 1.4 | 75.5 | 194.4 |

| | | | | | | | | |
|---|---|---|---|---|---|---|---|---|
| -0.07989 | 0.3814 | -0.2194 | -0.0963 | -0.06296 | 0.5107 | -0.1089 | 0.6044 | -0.5075 |
| -0.5047 to 0.3760 | -0.07371 to 0.7050 | -0.6034 to 0.2472 | -0.5169 to 0.3617 | 0.4919 to 0.3905 | 0.08804 to 0.7776 | -0.5261 to 0.3507 | 0.2209 to 0.8261 | -0.7758 to -0.08367 |
| 0.006382 | 0.1454 | 0.04814 | 0.009273 | 0.003964 | 0.2609 | 0.01185 | 0.3654 | 0.2575 |
| 0.7378 | 0.0971 | 0.3527 | 0.6863 | 0.792 | 0.0214 | 0.6478 | 0.0048 | 0.0224 |
| ns | ns | ns | ns | ns | * | ns | ** | * |
| No | No | No | No | No | Yes | No | Yes | Yes |
| 20 | 20 | 20 | 20 | 20 | 20 | 20 | 20 | 20 |

| | | | | | | | | |
|---|---|---|---|---|---|---|---|---|
| -0.05716 | -0.1913 | 0.2346 | -0.1072 | 0.2027 | 0.0857 | -0.1084 | -0.00657 | -0.02764 |
| -0.5533 to 0.4689 | -0.6409 to 0.3559 | -0.3157 to 0.6668 | -0.5873 to 0.4287 | 0.3456 to 0.647[illegible] | -0.4463 to 0.5729 | -0.5881 to 0.4277 | -0.5172 to 0.5075 | -0.5325 to 0.4917 |
| 0.003267 | 0.03661 | 0.05503 | 0.01149 | 0.04107 | 0.007345 | 0.01174 | 0.00004316 | 0.0007638 |
| 0.8397 | 0.4945 | 0.4 | 0.7038 | 0.4688 | 0.7614 | 0.7007 | 0.9815 | 0.9221 |
| ns | ns | ns | ns | ns | ns | ns | ns | ns |
| No | No | No | No | No | No | No | No | No |
| 15 | 15 | 15 | 15 | 15 | 15 | 15 | 15 | 15 |

(continued)

| 16 | 17 | 18 | 19 | 20 | 21 | 22 | 23 | 24 | 25 |
|---|---|---|---|---|---|---|---|---|---|
| **glutamic acid** | **glycine** | **histidine** | **isoleucine** | **leucine** | **lysine** | **methionine** | **hydroxyproline** | **ornithine** | **phosphoetanolamine** |
| 19.1 | 69.3 | 30.6 | 34.2 | 51.5 | 65.1 | 10.5 | 3 | 30.3 | 10 |
| 142.8 | 92.7 | 27.6 | 33.2 | 51.3 | 67.3 | 10.8 | 10.3 | 41.5 | 9.3 |
| 22.2 | 61.7 | 20.5 | 23.3 | 45.2 | 49.4 | 3.8 | 5.6 | 32.5 | 12.3 |
| 12.6 | 30.3 | 7.6 | 13.8 | 15.2 | 40.9 | 3.2 | 11.4 | 15.7 | 3.2 |
| 32.5 | 90.2 | 23.7 | 75.2 | 115.5 | 98.7 | 16.7 | 39 | 53.3 | 8.7 |
| 24.9 | 61.2 | 15 | 32.5 | 39 | 48.5 | 5.6 | 1.4 | 42.6 | 11.6 |
| 22.5 | 62.2 | 20.7 | 29.5 | 51.6 | 56.8 | 7.9 | 8.9 | 22.2 | 5.3 |
| 35.9 | 40 | 10.7 | 25.3 | 48.4 | 54.3 | 7.6 | 5.8 | 22.7 | 9.5 |
| 25.6 | 136.9 | 19.1 | 22.7 | 40.6 | 41.4 | 6 | 5.2 | 25.4 | 7.7 |
| 23.4 | 91.7 | 20.6 | 18.3 | 27.8 | 47 | 5.2 | 0 | 23.6 | 8 |
| 17.7 | 43.6 | 16.8 | 30.5 | 58.6 | 54.6 | 6.8 | 0.6 | 27.6 | 6.3 |
| 50.8 | 50.5 | 21.7 | 23.2 | 42.6 | 42.6 | 2 | 0 | 19.9 | 1.8 |
| 31 | 81.5 | 18.7 | 41.8 | 70 | 46.3 | 5.1 | 4.4 | 27 | 8.8 |
| 24.7 | 78.1 | 30.1 | 23 | 40.5 | 69 | 8.3 | 4 | 23.9 | 9.4 |
| 19.9 | 83.1 | 20.8 | 24 | 41.3 | 43.1 | 5.6 | 20.3 | 22.2 | 10.3 |
| 23 | 63.1 | 23.2 | 13.4 | 31.2 | 39.5 | 3.9 | 16.7 | 22.2 | 7.9 |
| 20 | 29.3 | 30.9 | 49.8 | 3.2 | 16.4 | 15.8 | | 3 | 60.5 |
| 13.7 | 90.5 | 14.9 | 26.1 | 34.6 | 51.7 | 6.1 | 15.4 | 23.6 | 1.9 |
| 57.3 | 22.2 | 37.1 | 46.2 | 63.7 | 11.1 | 6.3 | 40.9 | | 31.1 |
| 19.9 | 68.5 | 26.7 | 27.4 | 39.7 | 53.3 | 12.1 | 4.6 | 20.9 | 6.5 |

| | | | | | | | | | |
|---|---|---|---|---|---|---|---|---|---|
| 0.6107 | -0.04078 | 0.4546 | 0.2243 | 0.2526 | -0.3188 | -0.05187 | 0.5585 | 0.1539 | 0.1784 |
| 0.2303 to 0.8292 | -0.4748 to 0.4092 | 0.01496 to 0.7469 | -0.2424 to 0.6067 | -0.2139 to 0.6253 | -0.6673 to 0.1441 | -0.4834 to 0.3999 | 0.1396 to 0.8078 | -0.3230 to 0.5685 | -0.2869 to 0.5756 |
| 0.3729 | 0.001663 | 0.2066 | 0.0503 | 0.06381 | 0.1016 | 0.00269 | 0.3119 | 0.02369 | 0.03183 |
| 0.0042 | 0.8645 | 0.0441 | 0.3418 | 0.2826 | 0.1707 | 0.8281 | 0.0129 | 0.5293 | 0.4517 |
| ** | ns | * | ns | ns | ns | ns | * | ns | ns |
| Yes | No | Yes | No | No | No | No | Yes | No | No |
| 20 | 20 | 20 | 20 | 20 | 20 | 20 | 19 | 19 | 20 |

| | | | | | | | | | |
|---|---|---|---|---|---|---|---|---|---|
| 0.09475 | 0.2699 | 0.01212 | -0.2699 | -0.1033 | -0.1873 | -0.2076 | 0.1008 | -0.09838 | -0.1575 |
| -0.4389 to 0.5790 | -0.2814 to 0.6872 | -0.5034 to 0.5213 | -0.6872 to 0.2814 | -0.5847 to 0.4319 | -0.6384 to 0.3596 | -0.6508 to 0.3410 | -0.4541 to 0.5995 | -0.5814 to 0.4360 | -0.6198 to 0.3860 |
| 0.008977 | 0.07284 | 0.000147 | 0.07284 | 0.01067 | 0.03506 | 0.04311 | 0.01017 | 0.009679 | 0.0248 |
| 0.737 | 0.3306 | 0.9658 | 0.3306 | 0.7141 | 0.504 | 0.4578 | 0.7316 | 0.7272 | 0.5752 |
| ns | ns | ns | ns | ns | ns | ns | ns | ns | ns |
| No | No | No | No | No | No | No | No | No | No |
| 15 | 15 | 15 | 15 | 15 | 15 | 15 | 14 | 15 | 15 |

(continued)

| 26 | 27 | 28 | 29 | 30 | 31 | 32 | 33 | 34 | 35 |
|---|---|---|---|---|---|---|---|---|---|
| **phenylalanine** | **proline** | **phosphoserine** | **serine** | **taurine** | **threonine** | **tryptophan** | **tyrosine** | **urea** | **valine** |
| 29 | 112.3 | 7.2 | 34.5 | 19.1 | 65.7 | 11.4 | 34 | 3.1 | 105.5 |
| 88.3 | 112 | 10.3 | 43.1 | 46.8 | 54.8 | 20.4 | 35.4 | 3 | 91.9 |
| 45.6 | 90.7 | 8.1 | 24.9 | 18 | 26.3 | 15.3 | 30.3 | 3.4 | 85.6 |
| 14.5 | 33.7 | 7.7 | 17.6 | 25.3 | 20.5 | 4.4 | 11.3 | 1.1 | 35.2 |
| 49.4 | 102.3 | 5 | 45.4 | 24.3 | 67.9 | 36.2 | 48.4 | 3.9 | 189.3 |
| 24.8 | 76 | 7.3 | 32.9 | 19.5 | 37.5 | 6.8 | 34.5 | 3.9 | 75.3 |
| 21.6 | 67.8 | 7.7 | 33.2 | 60.9 | 37.4 | 17.2 | 31.6 | 2.1 | 83.9 |
| 25.1 | 47 | 6.5 | 20.3 | 22.5 | 23.7 | 8.8 | 27.8 | 2.5 | 89.3 |
| 31.9 | 87.3 | 6 | 38.9 | 21.8 | 41 | 10.7 | 26 | 3 | 84.4 |
| 19.3 | 64.9 | 6.4 | 28.1 | 18.7 | 35.8 | 2.1 | 25.9 | 2.1 | 55.7 |
| 30.9 | 71.9 | 8.3 | 32.5 | 14.3 | 38.6 | 0 | 28.2 | 2.1 | 90.4 |
| 23.5 | 51.7 | 7.7 | 27.9 | 82 | 23.6 | 12.7 | 18.9 | 3.4 | 77.4 |
| 23.3 | 42.8 | 7.2 | 40.9 | 28.4 | 32.6 | 15.9 | 22 | 2.7 | 126.1 |
| 28.3 | 85.6 | 7.9 | 35.3 | 22.6 | 38.7 | 12.2 | 34.5 | 5 | 84.4 |
| 22.9 | 73.3 | 8.1 | 29.2 | 21.7 | 29.5 | 13.5 | 25.5 | 1.6 | 84.2 |
| 22.8 | 50.9 | 9.8 | 29.9 | 21.6 | 25.6 | 9.9 | 23.1 | 2.4 | 66.6 |
| 8.8 | 29.4 | 13 | 20.8 | 2.7 | 22.8 | 2.1 | 76.8 | 1 | 2.9 |
| 27.3 | 89.9 | 7.3 | 37.9 | 19.6 | 47.5 | 0 | 29.5 | 2.1 | 78.5 |
| 129.8 | 6.6 | 37.5 | 51.3 | 39.9 | 1.9 | 33.4 | 1.9 | 95.8 | 1 |
| 27.5 | 97.7 | 6.5 | 39.5 | 14.1 | 48.8 | 17.3 | 32.7 | 2.4 | 81.3 |

| | | | | | | | | | |
|---|---|---|---|---|---|---|---|---|---|
| 0.8282 | -0.3019 | 0.7685 | 0.592 | 0.5301 | -0.3153 | 0.5605 | -0.3796 | 0.7796 | -0.2654 |
| 0.6087 to 0.9299 | -0.6567 to 0.1625 | 0.4939 to 0.9037 | 0.2025 to 0.8198 | 0.1144 to 0.7879 | -0.6651 to 0.1479 | 0.1568 to 0.8037 | 0.7039 to 0.0757 | 0.5146 to 0.9087 | 0.6336 to 0.2008 |
| 0.6859 | 0.09112 | 0.5906 | 0.3505 | 0.281 | 0.09944 | 0.3142 | 0.1441 | 0.6078 | 0.07043 |
| < 0.0001 | 0.1958 | < 0.0001 | 0.006 | 0.0162 | 0.1756 | 0.0101 | 0.0988 | < 0.0001 | 0.2581 |
| **** | ns | **** | ** | * | ns | * | ns | **** | ns |
| Yes | No | Yes | Yes | Yes | No | Yes | No | Yes | No |
| 20 | 20 | 20 | 20 | 20 | 20 | 20 | 20 | 20 | 20 |

| | | | | | | | | | |
|---|---|---|---|---|---|---|---|---|---|
| -0.06295 | -0.07165 | 0.08505 | 0.1591 | 0.3151 | -0.1183 | 0.07318 | -0.1484 | -0.02575 | -0.09277 |
| -0.5573 to 0.4644 | -0.5633 to 0.4575 | -0.4468 to 0.5725 | -0.3846 to 0.6209 | -0.2352 to 0.7125 | -0.5946 to 0.4195 | -0.4563 to 0.5644 | -0.6141 to 0.3939 | 0.5311 to 0.493 | 0.5777 to 0.4405 |
| 0.003963 | 0.005133 | 0.007233 | 0.02533 | 0.09931 | 0.01399 | 0.005355 | 0.02202 | 0.0006632 | 0.008606 |
| 0.8236 | 0.7997 | 0.7631 | 0.571 | 0.2526 | 0.6746 | 0.7955 | 0.5977 | 0.9274 | 0.7423 |
| ns | ns | ns | ns | ns | ns | ns | ns | ns | ns |
| No | No | No | No | No | No | No | No | No | No |
| 15 | 15 | 15 | 15 | 15 | 15 | 15 | 15 | 15 | 15 |

**Supplementary Table 3.** Clinical characteristics of the patients included in the *MYC* gene expression analysis depicted on Figure 6c.

| **Parameter** | **Number of patients** | **%** |
|---|---|---|
| | | In total: 74 patients, treated in 2000-2011 |
| Gleason score<br>≤6/7/8/9/missing | 41/23/5/4/1 | 55.4/31.1/6.8/5.4/1.4 |
| T stage<br>1/2/3/4 | 44/21/8/1 | 59.4/28.4/10.8/1.4 |
| D'Amico risk score<br>low/intermediate/high/missing | 24/26/23/1 | 32.4/35.1/31.1/1.4 |
| Hormone therapy<br>yes/no | 40/34 | 54.1/45.9 |
| Radiotherapy<br>external beam/low dose rate brachy therapy | 60/14 | 81.1/18.9 |

| **Parameter** | **Median** | **Range** |
|---|---|---|
| Age (years) | 71.2 | 55.5-85.0 |
| Dose (Gy) | 76.0 | 70.2-145.0 |
| Tumour volume (ml) | 34.0 | 16.0-121.0 (missing: 11) |
| PSA initial (ng/ml) | 7.3 | 0.39-62.4 |
| Follow-up time all patients (months) | 116.3 | 25.7-187.4 |

Supplementary Table 4. Characteristics of PCa patients (n = 14). Primary prostate cancer and adjacent normal tissues (benign hyperplasia, BPH) from these patients were used for primary tissue cultures and radiobiological colony forming assays.

| ID | Gleason score | TNM staging | PSA (ng/ml) | Age |
|---|---|---|---|---|
| BM_PC_11 RR | 5+5 | pT3ay pN0 | 38,0 ng/ml | 64 |
| BM_PC_12 RR | 4+3 | pT2c pN0 | 8,87 ng/ml | 76 |
| BM_PC_14* RR | 3+3 | pT2c pN0 | 7,85 ng/ml | 68 |
| BM_PC_16* RR | 3+4 | pT3a pN0 | 22,42 ng/ml | 63 |
| BM_PC_19 RS | 4+3 | cT1c | 7,17 ng/ml | 73 |
| BM_PC_20 RS | 4+3 | pT3b pN0 | 27,25 ng/ml | 65 |
| BM_PC_21 RR | 3+3 | pT3b pN1 | 8,11 ng/ml | 73 |
| BM_PC_22 | 4+3 | pT3b,pN0 | 8,77 ng/ml | 68 |
| BM_PC_23 RS | 5+4 | pT3b pN1 | 22,11 ng/ml | 65 |
| BM_PC_25 RS | 3+4 | pT2c pN0 | 6,4 ng/ml | 62 |
| BM_PC_26 RS | 4+3 | pT3b pN0 | 8,21 ng/ml | 76 |
| BM_PC_28 RS | 3+3 | pT3a, pN0 | 15,3 ng/ml | 66 |
| BM_PC_31 | 4+5 | pT3b, pN0 | 5,14 ng/ml | 68 |
| BM_PC_42 | 3+4 | pT2c pN0 | 10,38 ng/ml | 69 |

| | |
|---|---|
| RR | specimens used for the experiments depicted on Figure 2 g and characterized as radioresistant |
| RS | specimens used for the experiments depicted on Figure 2 g and characterized as radiosensitive |
| | specimens used for the experiments depicted on Figure 3 e |
| | * specimens used for both types of experiments |

Supplementary Table 5. Nucleotide sequences of siRNAs, primers used for qPCR and antibodies used for western blotting and fluorescence microscopy

| **siRNA** | **Nucleotide sequence** |
|---|---|
| Scr #1 | 5'-GGCUAAAGGAAACGAAAGA -3' |
| Scr #2 | 5'-UGCGCUAGGCCUCGGUUGC -3' |
| Scr #3 | 5'-GCAGCUAUAUGAAUGUUGU -3' |
| c-Myc #1 | 5'-GGAACGAGCUAAAACGGAG-3' |
| c-Myc #2 | 5'-GGAGGAGAACUUCUACCAG-3' |
| c-Myc #3 | 5'-CGACGAGACCUUCAUCAAA-3' |
| GLS #1 | 5'-GCAGUUCGAAAUACAUUGA-3' |
| GLS #2 | 5'-GGGUCUGUUACCUAGCUUG -3' |
| GLS #3 | 5'-GGACAAGAGAAAAUACCUG -3' |
| ATG5#1 | 5'-GGAUGAGAUAACUGAAAGG -3' |
| ATG5#2 | 5'-GGCAACCUGACCAGAAACA -3' |
| **Oligonucleotide primers** | **Nucleotide sequences** |
| c-Myc | Forward: 5'-CTCCGTCCTCGGATTCTCTGC-3'<br>Reverse: 5'-CTCCAGCAGAAGGTGATCCAG-3' |
| ACTB | Forward: 5'-ATGGAGTCCTGTGGCATCCA-3'<br>Reverse: 5'-AGTACTTGCGCTCAGGAGGA-3' |
| ATF3 | Forward: 5'- GTGAATGCTGAACTGAAGGC -3'<br>Reverse: 5'- CCATACCACGACTGCTTAGC -3' |
| ALKBH1 | Forward: 5'- AGAAGCGACTAAACGGAGACC -3'<br>Reverse: 5'- GGGAAAGGTGTGTAATGATCTGC -3' |
| ALKBH3 | Forward: 5'- GCTTGTGTCGCTCCTTCCAT -3'<br>Reverse: 5'- ACACGCACATTTGAGATGAGAA -3' |
| ATF4 | Forward: 5'- GGTCAGTCCCTCCAACAACAG -3'<br>Reverse: 5'- GACTAGGGGGGCAAAGAGATCAC -3' |
| GADD34 | Forward: 5'- GAGACAGAGGAAGAGGAAGCT -3'<br>Reverse: 5'- GGAAATGGACAGTGACCTTCTC -3' |
| GLS1 | Forward: 5'- GAGGCATTCTACTGGAGATACC-3'<br>Reverse: 5'- GCTCCAGCATTTACCATAGG-3' |
| IRE1 | Forward: 5'- CAGAATTGGTGCAGGCATCCC-3'<br>Reverse: 5'- GGTGTCGTACATGGTGATGGTG-3' |
| SLC1A5 | Forward: 5'- CACCATGGTTCTGGTCTCCTG-3'<br>Reverse: 5'-GGTGAAGAGGAAGTAGATGAGG-3 |
| B2M | Forward: 5'-TTTCATCCATCCGACATTGA-3'<br>Reverse: 5'-CCTCCATGATGCTGCTTACA-3' |

| **Antibodies / application** | **Manufacturer ID** |
|---|---|
| ACTB / Western blotting | Cell Signaling Technology, #4967 |
| ALDH1A1 / Immunofluorencence microscopy | Santa Cruz Biotechnology, (H-4), #sc-374076 |
| ATG5 / Western blotting | Cell Signaling Technology, #12994 |
| ATM / Western blotting | Cell Signaling Technology, #2873 |
| ATR / Western blotting | Cell Signaling Technology, #13934 |
| pATR / Western blotting | Cell Signaling Technology, #2853 |
| Chk1 / Western blotting | Cell Signaling Technology, #2360 |
| pChk1 S296 / Western blotting | Cell Signaling Technology, #2349 |
| GAPDH / Western blotting | Santa Cruz Biotechnology, #sc-25778 |
| GLS / Western blotting | Cell Signaling Technology, #88964 |
| γH2A.X / Western blotting, Immunofluorencence microscopy | Merck KGaA, # 05-636 |
| H3 / Western blotting | Cell Signaling Technology, #2650 |
| H3K27me3 / Western blotting | Cell Signaling Technology, #9733 |
| H3K36me3 / Western blotting | Cell Signaling Technology, #4909 |
| H3K9me3 / Western blotting | Cell Signaling Technology, #13969 |
| LC3B / Western blotting, Immunofluorencence microscopy | Cell Signaling Technology, #3868 |
| c-Myc / Western blotting | Cell signaling Technology, #9402 |
| PARP / Western blotting | Cell Signaling Technology, #9532 |
| cleaved-PARP / Western blotting | Cell Signaling Technology, #5625 |
| Tubulin / Western blotting | Cell Signaling Technology, #3873 |

### Supplementary Table 6. Targeted metabolomics of amino acids and biogenic amines

Concentration [**µM**]

| Cell Lines/Irridiation | Total Protein Mass | Ala | Arg | Asn | Asp | Cit | Gln |
|---|---|---|---|---|---|---|---|
| DU145 p9, line A | 56 | 133 | 2.9 | 15.2 | 33.6 | < LOD | < LOD |
| DU145 p9, line A | 56 | 144 | 2.54 | 15.8 | 18.7 | < LOD | < LOD |
| DU145 p9, line B | 44 | 102 | 4.36 | 11.8 | 34.2 | < LOD | 8.37 |
| DU145 p9, line B | 44 | 105 | 2.99 | 12.7 | 32.8 | < LOD | 3.49 |
| DU145 RR p9, line A, 21x4Gy, Irradiated 25.01.16 | 35 | 77.6 | 1.52 | 16.4 | 18.4 | < LOD | 27.4 |
| DU145 RR p9, line A, 21x4Gy, Irradiated 25.01.17 | 35 | 76.4 | 2.6 | 16.9 | 24.5 | < LOD | 36.9 |
| DU145 RR p9, line B, 21x4Gy, Irradiated 25.01.18 | 48 | 77.6 | 4.72 | 14.3 | 25.1 | < LOD | 35.1 |
| DU145 RR p9, line B, 21x4Gy, Irradiated 25.01.19 | 48 | 76.7 | 3.59 | 15.1 | 21.7 | < LOD | 34.3 |
| LNCaP p9, line A | 32 | 102 | 13.9 | 43.1 | 22 | < LOD | 27.8 |
| LNCaP p9, line A | 32 | 113 | 18.5 | 47.3 | 18.3 | < LOD | 30.5 |
| LNCaP p9, line B | 29 | 87.6 | 12.6 | 37.4 | 19.4 | < LOD | 27.4 |
| LNCaP p9, line B | 29 | 108 | 23.8 | 48.9 | 27.5 | < LOD | 32.7 |
| LNCaP RR p9, line A, 21x4Gy, Irradiated 25.01.16 | 50 | 126 | 16.1 | 39.9 | 5.59 | < LOD | < LOD |
| LNCaP RR p9, line A, 21x4Gy, Irradiated 25.01.17 | 50 | 129 | 17.5 | 44.8 | 3.29 | < LOD | 2.68 |
| LNCaP RR p9, line B, 21x4Gy, Irradiated 25.01.18 | 44 | 102 | 17.5 | 42.1 | 11 | < LOD | 23.2 |
| LNCaP RR p9, line B, 21x4Gy, Irradiated 25.01.19 | 44 | 89.8 | 13.6 | 37.6 | 9.68 | < LOD | 20 |
| PC3 p8, line A | 26 | 27.6 | 4.93 | 5.54 | 29.9 | < LOD | 64.4 |
| PC3 p8, line A | 26 | 34.7 | 9.29 | 7.04 | 31.5 | < LOD | 78.7 |
| PC3 p8, line B | 33 | 32.2 | 6.61 | 6.97 | 25.5 | < LOD | 65.8 |
| PC3 p8, line B | 33 | 28.9 | 2.76 | 5.29 | 27.5 | < LOD | 53.7 |
| PC3 RR p8, line A, 14x4Gy, Irradiated 27.01.16 | 19 | 21.9 | 2.54 | 4.41 | 27.6 | < LOD | 40.3 |
| PC3 RR p8, line A, 14x4Gy, Irradiated 27.01.17 | 19 | 21.1 | 2 | 4.39 | 24.6 | < LOD | 45.1 |
| PC3 RR p8, line B, 14x4Gy, Irradiated 27.01.18 | 18 | 23.6 | 4.04 | 4.73 | 33.1 | < LOD | 54.6 |
| PC3 RR p8, line B, 14x4Gy, Irradiated 27.01.19 | 18 | 24.8 | 1.6 | 3.79 | 21.8 | < LOD | 48.9 |
| DU145 p9, line A + 24h irradiated | 52 | 72.9 | 1.27 | 7.24 | < LOD | < LOD | < LOD |
| DU145 p9, line A + 24h irradiated | 52 | 100 | 1.93 | 10.3 | < LOD | < LOD | < LOD |
| DU145 RR p9, line A, 21x4Gy, Irradiated 25.01.16 + 24h irradiated | 77 | 92.3 | 0.888 | 13.6 | 8.32 | < LOD | < LOD |
| DU145 RR p9, line A, 21x4Gy, Irradiated 25.01.16 + 24h irradiated | 77 | 98.1 | 1.05 | 13.9 | 12.2 | < LOD | < LOD |
| Medium control | | < LOD | < LOD | < LOD | < LOD | < LOD | < LOD |
| Medium control | | < LOD | < LOD | < LOD | < LOD | < LOD | < LOD |

Concentration [**pmol/10$^6$ cells**]

| Cell Lines/Irradiation | Total Protein Mass | Ala | Arg | Asn | Asp | Cit | Gln |
|---|---|---|---|---|---|---|---|
| DU145 p9, line A | 56 | 23750000 | 517857.1429 | 2714285.714 | 6000000 | < LOD | < LOD |
| DU145 p9, line A | 56 | 25714285.71 | 453571.4286 | 2821428.571 | 3339285.714 | < LOD | < LOD |
| DU145 p9, line B | 44 | 23181818.18 | 990909.0909 | 2681818.182 | 7772727.273 | < LOD | 1902272.727 |
| DU145 p9, line B | 44 | 23863636.36 | 679545.4545 | 2886363.636 | 7454545.455 | < LOD | 793181.8182 |
| DU145 RR p9, line A, 21x4Gy, Irradiated 25.01.16 | 35 | 22171428.57 | 434285.7143 | 4685714.286 | 5257142.857 | < LOD | 7828571.429 |
| DU145 RR p9, line A, 21x4Gy, Irradiated 25.01.17 | 35 | 21828571.43 | 742857.1429 | 4828571.429 | 7000000 | < LOD | 10542857.14 |
| DU145 RR p9, line B, 21x4Gy, Irradiated 25.01.18 | 48 | 16166666.67 | 316666.6667 | 3416666.667 | 3833333.333 | < LOD | 5708333.333 |
| DU145 RR p9, line B, 21x4Gy, Irradiated 25.01.19 | 48 | 15916666.67 | 541666.6667 | 3520833.333 | 5104166.667 | < LOD | 7687500 |
| LNCaP p9, line A | 32 | 31875000 | 4343750 | 13468750 | 6875000 | < LOD | 8687500 |
| LNCaP p9, line A | 32 | 35312500 | 5781250 | 14781250 | 5718750 | < LOD | 9531250 |
| LNCaP p9, line B | 29 | 30206896.55 | 4344827.586 | 12896551.72 | 6689655.172 | < LOD | 9448275.862 |
| LNCaP p9, line B | 29 | 37241379.31 | 8206896.552 | 16862068.97 | 9482758.621 | < LOD | 11275862.07 |
| LNCaP RR p9, line A, 21x4Gy, Irradiated 25.01.16 | 50 | 25200000 | 3220000 | 7980000 | 1118000 | < LOD | < LOD |
| LNCaP RR p9, line A, 21x4Gy, Irradiated 25.01.17 | 50 | 25800000 | 3500000 | 8960000 | 658000 | < LOD | 536000 |
| LNCaP RR p9, line B, 21x4Gy, Irradiated 25.01.18 | 44 | 23181818.18 | 3977272.727 | 9568181.818 | 2500000 | < LOD | 5272727.273 |
| LNCaP RR p9, line B, 21x4Gy, Irradiated 25.01.19 | 44 | 20409090.91 | 3090909.091 | 8545454.545 | 2200000 | < LOD | 4545454.545 |
| PC3 p8, line A | 26 | 10615384.62 | 1896153.846 | 2130769.231 | 11500000 | < LOD | 24769230.77 |
| PC3 p8, line A | 26 | 13346153.85 | 3573076.923 | 2707692.308 | 12115384.62 | < LOD | 30269230.77 |
| PC3 p8, line B | 33 | 9757575.758 | 2003030.303 | 2112121.212 | 7727272.727 | < LOD | 19939393.94 |
| PC3 p8, line B | 33 | 8757575.758 | 836363.6364 | 1603030.303 | 8333333.333 | < LOD | 16272727.27 |
| PC3 RR p8, line A, 14x4Gy, Irradiated 27.01.16 | 19 | 11526315.79 | 1336842.105 | 2321052.632 | 14526315.79 | < LOD | 21210526.32 |
| PC3 RR p8, line A, 14x4Gy, Irradiated 27.01.17 | 19 | 11105263.16 | 1052631.579 | 2310526.316 | 12947368.42 | < LOD | 23736842.11 |
| PC3 RR p8, line B, 14x4Gy, Irradiated 27.01.18 | 18 | 13111111.11 | 2244444.444 | 2627777.778 | 18388888.89 | < LOD | 30333333.33 |
| PC3 RR p8, line B, 14x4Gy, Irradiated 27.01.19 | 18 | 13777777.78 | 888888.8889 | 2105555.556 | 12111111.11 | < LOD | 27166666.67 |
| DU145 p9, line A + 24h irradiated | 52 | 14019230.77 | 244230.7692 | 1392307.692 | < LOD | < LOD | < LOD |
| DU145 p9, line A + 24h irradiated | 52 | 19230769.23 | 371153.8462 | 1980769.231 | < LOD | < LOD | < LOD |
| DU145 RR p9, line A, 21x4Gy, Irradiated 25.01.16 + 24h irradiated | 77 | 11987012.99 | 115324.6753 | 1766233.766 | 1080519.481 | < LOD | < LOD |
| DU145 RR p9, line A, 21x4Gy, Irradiated 25.01.16 + 24h irradiated | 77 | 12740259.74 | 136363.6364 | 1805194.805 | 1584415.584 | < LOD | < LOD |

| Cell Lines/Irridiation | p-values (t-test) | Ala | Arg | Asn | Asp | Cit | Gln |
|---|---|---|---|---|---|---|---|
| DU145 vs DU145 RR | | 0.030162485 | 0.352027018 | 0.012079201 | 0.50939231 | | 0.012481778 |
| LNCaP vs LNCaP RR | | 0.002491473 | 0.054468031 | 0.00088859 | 0.000894671 | | 0.006314457 |
| PC3 vs PC3 RR | | 0.183689218 | 0.318175256 | 0.448306837 | 0.042075717 | | 0.469804156 |
| DU145 vs DU145 + 4Gy, 24 h | | 0.013950174 | 0.127818838 | 0.004912586 | | | |
| DU145 RR vs DU145 RR + 4Gy, 24 h | | 0.06218887 | 0.047940296 | 0.014248204 | 0.015949674 | | |

(continued)

| Glu | Gly | His | Ile | Leu | Lys | Met | Orn | Phe | Pro | Ser | Thr |
|---|---|---|---|---|---|---|---|---|---|---|---|
| 216 | 123 | 6.58 | 18.9 | 17.9 | 2.2 | 8.53 | < LOD | 13.7 | 30.1 | 92.8 | 129 |
| 202 | 151 | 7.86 | 23.9 | 29.5 | 1.68 | 11.3 | < LOD | 15.9 | 32.8 | 108 | 139 |
| 254 | 111 | 7.8 | 26.3 | 41.1 | 6.68 | 11.3 | < LOD | 15.8 | 26.3 | 95.3 | 102 |
| 266 | 104 | 6.64 | 21.2 | 17.5 | 2.05 | 8.48 | < LOD | 13.8 | 25.5 | 90.4 | 103 |
| 357 | 127 | 5.12 | 19.1 | 20.7 | < LOD | 5.06 | < LOD | 13.3 | 43.2 | 53.2 | 85.5 |
| 384 | 145 | 6.62 | 23.5 | 33 | 2.86 | 5.82 | < LOD | 15.5 | 48.7 | 60.6 | 85 |
| 346 | 144 | 7.78 | 28.4 | 19 | 7.67 | 7.01 | < LOD | 17.5 | 44.9 | 64.5 | 86.7 |
| 357 | 141 | 6.92 | 24.7 | 15.8 | 5.13 | 6.09 | < LOD | 15.1 | 45.8 | 62.8 | 86 |
| 111 | 131 | 0.759 | 6.47 | 10.9 | < LOD | 0.207 | 5.98 | 1.45 | 25.4 | 52.3 | 19.2 |
| 97.2 | 165 | 1.67 | 7.6 | 12.2 | 1.63 | 0.162 | 10.4 | 2.34 | 30.5 | 62.4 | 21.3 |
| 104 | 100 | < LOD | 5.05 | 9.81 | < LOD | < LOD | 4.82 | 1.24 | 18.3 | 46.2 | 17.3 |
| 125 | 129 | 2.32 | 7.97 | 21.3 | 4.36 | 0.567 | 13.5 | 3.26 | 27.1 | 57.8 | 22.1 |
| 73.3 | 221 | 0.833 | 5.31 | 10.6 | < LOD | < LOD | 7.13 | 1.53 | 37.9 | 39.9 | 20.5 |
| 82.4 | 244 | 1.1 | 5.66 | 6.36 | < LOD | < LOD | 10.3 | 1.82 | 43.6 | 45.1 | 21.5 |
| 115 | 151 | 1.05 | 5.9 | 7.03 | < LOD | 0.22 | 6.33 | 1.92 | 31.9 | 50.1 | 19.7 |
| 110 | 141 | < LOD | 4.72 | 5.26 | < LOD | < LOD | 4.16 | 1.44 | 28.9 | 44.2 | 17.2 |
| 130 | 51.9 | 5.15 | 17.3 | 15.1 | 6.8 | 7.38 | < LOD | 11.6 | 9.95 | 33.5 | 43.8 |
| 138 | 64.1 | 7.88 | 24.2 | 30.8 | 17.3 | 8.33 | < LOD | 15.6 | 11.6 | 42.5 | 54.9 |
| 135 | 55.7 | 6.01 | 17.9 | 25.5 | 10.7 | 7.8 | < LOD | 12.7 | 9.73 | 32.5 | 50.8 |
| 129 | 53.2 | 4.69 | 17.4 | 15.9 | 1.22 | 7.51 | < LOD | 10.4 | 10.1 | 31.5 | 47 |
| 153 | 31 | 3.59 | 14.4 | 12.4 | 1.77 | 4.3 | < LOD | 8.21 | 5.52 | 24 | 31.9 |
| 136 | 33.1 | 3.51 | 13.6 | 10.1 | 0.97 | 4.48 | < LOD | 8.05 | 5.74 | 25.5 | 33.3 |
| 146 | 35 | 4.68 | 16.7 | 27.4 | 5.32 | 5.16 | < LOD | 10.2 | 6.37 | 28.2 | 37.8 |
| 162 | 35.3 | 3.55 | 12.5 | 18.6 | < LOD | 4.39 | < LOD | 8.42 | 6.26 | 27.1 | 35.1 |
| 15.4 | 87.3 | 4.54 | 13.7 | 19.2 | < LOD | 6.17 | < LOD | 11.2 | 20 | 62.9 | 83.5 |
| 37.5 | 110 | 6.62 | 19.7 | 13.7 | < LOD | 7.97 | < LOD | 13.2 | 24.5 | 72.8 | 108 |
| 253 | 131 | 3.32 | 12.8 | 11.5 | < LOD | 2.84 | < LOD | 10.4 | 49.4 | 33.1 | 83.3 |
| 287 | 156 | 4.49 | 17.9 | 16.6 | < LOD | 3.32 | < LOD | 12 | 55.6 | 42.2 | 93.8 |
| < LOD | < LOD | < LOD | < LOD | < LOD | < LOD | < LOD | < LOD | < LOD | < LOD | < LOD | < LOD |
| < LOD | < LOD | < LOD | < LOD | 2.8 | < LOD | < LOD | < LOD | < LOD | < LOD | < LOD | < LOD |

| Glu | Gly | His | Ile | Leu | Lys | Met | Orn | Phe | Pro | Ser | Thr |
|---|---|---|---|---|---|---|---|---|---|---|---|
| 38571428.57 | 21964285.71 | 1175000 | 3375000 | 3196428.571 | 392857.1429 | 1523214.286 | < LOD | 2446428.571 | 5375000 | 16571428.57 | 23035714.29 |
| 36071428.57 | 26964285.71 | 1403571.429 | 4267857.143 | 5267857.143 | 300000 | 2017857.143 | < LOD | 2839285.714 | 5857142.857 | 19285714.29 | 24821428.57 |
| 57727272.73 | 25227272.73 | 1772727.273 | 5977272.727 | 9340909.091 | 1518181.818 | 2568181.818 | < LOD | 3590909.091 | 5977272.727 | 21659090.91 | 23181818.18 |
| 60454545.45 | 23636363.64 | 1509090.909 | 4818181.818 | 3977272.727 | 465909.0909 | 1927272.727 | < LOD | 3136363.636 | 5795454.545 | 20545454.55 | 23409090.91 |
| 102000000 | 36285714.29 | 1462857.143 | 5457142.857 | 5914285.714 | | 1445714.286 | < LOD | 3800000 | 12342857.14 | 15200000 | 24428571.43 |
| 109714285.7 | 41428571.43 | 1891428.571 | 6714285.714 | 9428571.429 | 817142.8571 | 1662857.143 | < LOD | 4428571.429 | 13914285.71 | 17314285.71 | 24285714.29 |
| 74375000 | 26458333.33 | 1066666.667 | 3979166.667 | 4312500 | | 1054166.667 | < LOD | 2770833.333 | 9000000 | 11083333.33 | 17812500 |
| 80000000 | 30208333.33 | 1379166.667 | 4895833.333 | 6875000 | 595833.3333 | 1212500 | < LOD | 3229166.667 | 10145833.33 | 12625000 | 17708333.33 |
| 34687500 | 40937500 | 237187.5 | 2021875 | 3406250 | < LOD | 64687.5 | 1868750 | 453125 | 7937500 | 16343750 | 6000000 |
| 30375000 | 51562500 | 521875 | 2375000 | 3812500 | 509375 | 50625 | 3250000 | 731250 | 9531250 | 19500000 | 6656250 |
| 35862068.97 | 34482758.62 | < LOD | 1741379.31 | 3382758.621 | < LOD | < LOD | 1662068.966 | 427586.2069 | 6310344.828 | 15931034.48 | 5965517.241 |
| 43103448.28 | 44482758.62 | 800000 | 2748275.862 | 7344827.586 | 1503448.276 | 195517.2414 | 4655172.414 | 1124137.931 | 9344827.586 | 19931034.48 | 7620689.655 |
| 14660000 | 44200000 | 166600 | 1062000 | 2120000 | < LOD | < LOD | 1426000 | 306000 | 7580000 | 7980000 | 4100000 |
| 16480000 | 48800000 | 220000 | 1132000 | 1272000 | < LOD | < LOD | 2060000 | 364000 | 8720000 | 9020000 | 4300000 |
| 26136363.64 | 34318181.82 | 238636.3636 | 1340909.091 | 1597727.273 | < LOD | 50000 | 1438636.364 | 436363.6364 | 7250000 | 11386363.64 | 4477272.727 |
| 25000000 | 32045454.55 | < LOD | 1072727.273 | 1195454.545 | < LOD | < LOD | 945454.5455 | 327272.7273 | 6568181.818 | 10045454.55 | 3909090.909 |
| 50000000 | 19961538.46 | 1980769.231 | 6653846.154 | 5807692.308 | 2615384.615 | 2838461.538 | < LOD | 4461538.462 | 3826923.077 | 12884615.38 | 16846153.85 |
| 53076923.08 | 24653846.15 | 3030769.231 | 9307692.308 | 11846153.85 | 6653846.154 | 3203846.154 | < LOD | 6000000 | 4461538.462 | 16346153.85 | 21115384.62 |
| 40909090.91 | 16878787.88 | 1821212.121 | 5424242.424 | 7727272.727 | 3242424.242 | 2363636.364 | < LOD | 3848484.848 | 2948484.848 | 9848484.848 | 15393939.39 |
| 39090909.09 | 16121212.12 | 1421212.121 | 5272727.273 | 4818181.818 | 369696.9697 | 2275757.576 | < LOD | 3151515.152 | 3060606.061 | 9545454.545 | 14242424.24 |
| 80526315.79 | 16315789.47 | 1889473.684 | 7578947.368 | 6526315.789 | 931578.9474 | 2263157.895 | < LOD | 4321052.632 | 2905263.158 | 12631578.95 | 16789473.68 |
| 71578947.37 | 17421052.63 | 1847368.421 | 7157894.737 | 5315789.474 | 510526.3158 | 2357894.737 | < LOD | 4236842.105 | 3021052.632 | 13421052.63 | 17526315.79 |
| 81111111.11 | 19444444.44 | 2600000 | 9277777.778 | 15222222.22 | 2955555.556 | 2866666.667 | < LOD | 5666666.667 | 3538888.889 | 15666666.67 | 21000000 |
| 90000000 | 19611111.11 | 1972222.222 | 6944444.444 | 10333333.33 | < LOD | 2438888.889 | < LOD | 4677777.778 | 3477777.778 | 15055555.56 | 19500000 |
| 2961538.462 | 16788461.54 | 873076.9231 | 2634615.385 | 3692307.692 | < LOD | 1186538.462 | < LOD | 2153846.154 | 3846153.846 | 12096153.85 | 16057692.31 |
| 7211538.462 | 21153846.15 | 1273076.923 | 3788461.538 | 2634615.385 | < LOD | 1532692.308 | < LOD | 2538461.538 | 4711538.462 | 14000000 | 20769230.77 |
| 32857142.86 | 17012987.01 | 431168.8312 | 1662337.662 | 1493506.494 | < LOD | 368831.1688 | < LOD | 1350649.351 | 6415584.416 | 4298701.299 | 10818181.82 |
| 37272727.27 | 20259740.26 | 583116.8831 | 2324675.325 | 2155844.156 | < LOD | 431168.8312 | < LOD | 1558441.558 | 7220779.221 | 5480519.481 | 12181818.18 |

| Glu | Gly | His | Ile | Leu | Lys | Met | Orn | Phe | Pro | Ser | Thr |
|---|---|---|---|---|---|---|---|---|---|---|---|
| 0.006454703 | 0.038876667 | 0.945250849 | 0.44053649 | 0.519827163 | 0.935554753 | 0.039130357 | | 0.247536515 | 0.002328088 | 0.021075263 | 0.237969921 |
| 0.007800999 | 0.591865919 | 0.130364886 | 0.003329638 | 0.024058791 | | | 0.105786587 | 0.095142295 | 0.421781935 | 0.00060389 | 0.001135342 |
| 0.000457772 | 0.586036624 | 0.972674965 | 0.35516379 | 0.532574725 | 0.339661096 | 0.485857978 | | 0.620326914 | 0.417038867 | 0.285071358 | 0.350059228 |
| 0.01082608 | 0.058761109 | 0.153164989 | 0.191155675 | 0.333294595 | | 0.128421243 | | 0.159630171 | 0.011127705 | 0.020615729 | 0.0295318 |
| 0.011672744 | 0.041583889 | 0.022030731 | 0.020717233 | 0.041221745 | | 0.0093226 | | 0.017833889 | 0.053229272 | 0.012006577 | 0.029637855 |

(continued)

| Trp | Tyr | Val | ADMA | Ac-Orn | Carnosine | Creatinine | DOPA | Dopamine | Histamine | Kynurenine | Met-SO |
|---|---|---|---|---|---|---|---|---|---|---|---|
| 1.73 | 14.1 | 20.8 | < LOD | < LOD | < LOD | < LOD | < LOD | < LOD | < LOD | < LOD | < LOD |
| 2.6 | 15.7 | 23.6 | < LOD | < LOD | < LOD | < LOD | < LOD | < LOD | < LOD | < LOD | 0.821 |
| 2.34 | 15.7 | 24.6 | < LOD | < LOD | < LOD | < LOD | < LOD | < LOD | < LOD | < LOD | 0.549 |
| 1.94 | 14 | 20.5 | < LOD | < LOD | < LOD | < LOD | < LOD | < LOD | < LOD | < LOD | < LOD |
| 1.59 | 12.3 | 18.4 | < LOD | < LOD | < LOD | < LOD | < LOD | < LOD | < LOD | < LOD | < LOD |
| 2.45 | 15.6 | 23.4 | < LOD | < LOD | < LOD | < LOD | < LOD | < LOD | < LOD | < LOD | < LOD |
| 2.75 | 16.7 | 26.5 | < LOD | < LOD | < LOD | < LOD | < LOD | < LOD | 0.028 | < LOD | 0.746 |
| 2.29 | 15.8 | 23 | < LOD | < LOD | < LOD | < LOD | < LOD | < LOD | < LOD | < LOD | 0.565 |
| < LOD | 1.32 | < LOD | < LOD | < LOD | < LOD | < LOD | < LOD | < LOD | < LOD | < LOD | 0.904 |
| < LOD | 2.39 | 1.87 | < LOD | < LOD | < LOD | 2.09 | < LOD | < LOD | < LOD | < LOD | 1.14 |
| < LOD | 1.07 | < LOD | < LOD | < LOD | < LOD | < LOD | < LOD | < LOD | < LOD | < LOD | 0.643 |
| < LOD | 3.27 | 3.43 | < LOD | < LOD | < LOD | < LOD | < LOD | < LOD | < LOD | < LOD | 0.889 |
| < LOD | 1.6 | < LOD | < LOD | < LOD | < LOD | < LOD | < LOD | < LOD | < LOD | < LOD | 0.871 |
| < LOD | 1.55 | < LOD | < LOD | < LOD | < LOD | 2.13 | < LOD | < LOD | < LOD | < LOD | 1.54 |
| < LOD | 1.92 | 0.842 | < LOD | < LOD | < LOD | < LOD | < LOD | < LOD | < LOD | < LOD | 0.666 |
| < LOD | 1.4 | < LOD | < LOD | < LOD | < LOD | < LOD | < LOD | < LOD | < LOD | < LOD | 0.988 |
| 1.69 | 11.4 | 17.6 | < LOD | < LOD | < LOD | < LOD | < LOD | < LOD | < LOD | < LOD | < LOD |
| 2.52 | 15.5 | 26 | < LOD | < LOD | < LOD | < LOD | < LOD | < LOD | < LOD | < LOD | 0.376 |
| 1.74 | 13.1 | 20.2 | < LOD | < LOD | < LOD | < LOD | < LOD | < LOD | < LOD | < LOD | < LOD |
| 1.21 | 10.7 | 15 | < LOD | < LOD | < LOD | < LOD | < LOD | < LOD | < LOD | < LOD | < LOD |
| 1.06 | 8.59 | 11.6 | < LOD | < LOD | < LOD | < LOD | < LOD | < LOD | < LOD | < LOD | < LOD |
| 0.869 | 8.55 | 12.5 | < LOD | < LOD | < LOD | < LOD | < LOD | < LOD | < LOD | < LOD | < LOD |
| 1.47 | 10.3 | 16 | < LOD | < LOD | < LOD | < LOD | < LOD | < LOD | < LOD | < LOD | < LOD |
| 0.856 | 8.08 | 12.3 | < LOD | < LOD | < LOD | < LOD | < LOD | < LOD | < LOD | < LOD | < LOD |
| 1.28 | 10.3 | 14.3 | < LOD | < LOD | < LOD | < LOD | < LOD | < LOD | < LOD | < LOD | < LOD |
| 1.61 | 13.5 | 19 | < LOD | < LOD | < LOD | < LOD | < LOD | < LOD | < LOD | < LOD | 0.584 |
| 1.23 | 9.62 | 13.6 | < LOD | < LOD | < LOD | < LOD | < LOD | < LOD | < LOD | < LOD | < LOD |
| 1.46 | 12.3 | 16.3 | < LOD | < LOD | < LOD | < LOD | < LOD | < LOD | < LOD | < LOD | < LOD |
| < LOD | < LOD | < LOD | < LOD | < LOD | < LOD | < LOD | < LOD | < LOD | < LOD | < LOD | < LOD |
| < LOD | < LOD | < LOD | < LOD | < LOD | < LOD | < LOD | < LOD | < LOD | < LOD | < LOD | < LOD |

| Trp | Tyr | Val | ADMA | Ac-Orn | Carnosine | Creatinine | DOPA | Dopamine | Histamine | Kynurenine | Met-SO |
|---|---|---|---|---|---|---|---|---|---|---|---|
| 308928.5714 | 2517857.143 | 3714285.714 | < LOD | < LOD | < LOD | < LOD | < LOD | < LOD | < LOD | < LOD | < LOD |
| 464285.7143 | 2803571.429 | 4214285.714 | < LOD | < LOD | < LOD | < LOD | < LOD | < LOD | < LOD | < LOD | 146607.1429 |
| 531818.1818 | 3568181.818 | 5590909.091 | < LOD | < LOD | < LOD | < LOD | < LOD | < LOD | < LOD | < LOD | 124772.7273 |
| 440909.0909 | 3181818.182 | 4659090.909 | < LOD | < LOD | < LOD | < LOD | < LOD | < LOD | < LOD | < LOD | < LOD |
| 454285.7143 | 3514285.714 | 5257142.857 | < LOD | < LOD | < LOD | < LOD | < LOD | < LOD | < LOD | < LOD | < LOD |
| 700000 | 4457142.857 | 6685714.286 | < LOD | < LOD | < LOD | < LOD | < LOD | < LOD | < LOD | < LOD | < LOD |
| 331250 | 2562500 | 3833333.333 | < LOD | < LOD | < LOD | < LOD | < LOD | < LOD | < LOD | < LOD | < LOD |
| 510416.6667 | 3250000 | 4875000 | < LOD | < LOD | < LOD | < LOD | < LOD | < LOD | < LOD | < LOD | < LOD |
| < LOD | 412500 | < LOD | < LOD | < LOD | < LOD | < LOD | < LOD | < LOD | < LOD | < LOD | 282500 |
| < LOD | 746875 | 584375 | < LOD | < LOD | < LOD | 653125 | < LOD | < LOD | < LOD | < LOD | 356250 |
| < LOD | 368965.5172 | < LOD | < LOD | < LOD | < LOD | < LOD | < LOD | < LOD | < LOD | < LOD | 221724.1379 |
| < LOD | 1127586.207 | 1182758.621 | < LOD | < LOD | < LOD | < LOD | < LOD | < LOD | < LOD | < LOD | 306551.7241 |
| < LOD | 320000 | < LOD | < LOD | < LOD | < LOD | < LOD | < LOD | < LOD | < LOD | < LOD | 174200 |
| < LOD | 310000 | < LOD | < LOD | < LOD | < LOD | 426000 | < LOD | < LOD | < LOD | < LOD | 308000 |
| < LOD | 436363.6364 | 191363.6364 | < LOD | < LOD | < LOD | < LOD | < LOD | < LOD | < LOD | < LOD | 151363.6364 |
| < LOD | 318181.8182 | < LOD | < LOD | < LOD | < LOD | < LOD | < LOD | < LOD | < LOD | < LOD | 224545.4545 |
| 650000 | 4384615.385 | 6769230.769 | < LOD | < LOD | < LOD | < LOD | < LOD | < LOD | < LOD | < LOD | < LOD |
| 969230.7692 | 5961538.462 | 10000000 | < LOD | < LOD | < LOD | < LOD | < LOD | < LOD | < LOD | < LOD | 144615.3846 |
| 527272.7273 | 3969696.97 | 6121212.121 | < LOD | < LOD | < LOD | < LOD | < LOD | < LOD | < LOD | < LOD | < LOD |
| 366666.6667 | 3242424.242 | 4545454.545 | < LOD | < LOD | < LOD | < LOD | < LOD | < LOD | < LOD | < LOD | < LOD |
| 557894.7368 | 4521052.632 | 6105263.158 | < LOD | < LOD | < LOD | < LOD | < LOD | < LOD | < LOD | < LOD | < LOD |
| 457368.4211 | 4500000 | 6578947.368 | < LOD | < LOD | < LOD | < LOD | < LOD | < LOD | < LOD | < LOD | < LOD |
| 816666.6667 | 5722222.222 | 8888888.889 | < LOD | < LOD | < LOD | < LOD | < LOD | < LOD | < LOD | < LOD | < LOD |
| 475555.5556 | 4488888.889 | 6833333.333 | < LOD | < LOD | < LOD | < LOD | < LOD | < LOD | < LOD | < LOD | < LOD |
| 246153.8462 | 1980769.231 | 2750000 | < LOD | < LOD | < LOD | < LOD | < LOD | < LOD | < LOD | < LOD | < LOD |
| 309615.3846 | 2596153.846 | 3653846.154 | < LOD | < LOD | < LOD | < LOD | < LOD | < LOD | < LOD | < LOD | 112307.6923 |
| 159740.2597 | 1249350.649 | 1766233.766 | < LOD | < LOD | < LOD | < LOD | < LOD | < LOD | < LOD | < LOD | < LOD |
| 189610.3896 | 1597402.597 | 2116883.117 | < LOD | < LOD | < LOD | < LOD | < LOD | < LOD | < LOD | < LOD | < LOD |

| Trp | Tyr | Val | ADMA | Ac-Orn | Carnosine | Creatinine | DOPA | Dopamine | Histamine | Kynurenine | Met-SO |
|---|---|---|---|---|---|---|---|---|---|---|---|
| 0.512573596 | 0.381930868 | 0.418710509 | | | | | | | | | |
| | 0.125523892 | | | | | | | | | | 0.133736439 |
| 0.746891133 | 0.543826715 | 0.858171886 | | | | | | | | | |
| 0.094483329 | 0.135476741 | 0.111185876 | | | | | | | | | |
| 0.048389917 | 0.027713469 | 0.022500939 | | | | | | | | | |

(continued)

| Nitro-Tyr | PEA | Putrescine | SDMA | Serotonin | Spermidine | Spermine | Taurine | alpha-AAA | c4-OH-Pro | t4-OH-Pro |
|---|---|---|---|---|---|---|---|---|---|---|
| < LOD | < LOD | 4.11 | < LOD | < LOD | 2.83 | 0.767 | 63 | < LOD | < LOD | 1.75 |
| < LOD | < LOD | 4.99 | < LOD | < LOD | 4.34 | 1.37 | 68.1 | < LOD | < LOD | 2.04 |
| < LOD | < LOD | 3.79 | < LOD | < LOD | 7.64 | 2.14 | 56.3 | < LOD | < LOD | 1.67 |
| < LOD | < LOD | 3.77 | < LOD | < LOD | 5.24 | 1.9 | 58.8 | < LOD | < LOD | 1.56 |
| < LOD | < LOD | 1.26 | < LOD | < LOD | 5.44 | 3.54 | 76 | < LOD | < LOD | 1.41 |
| < LOD | < LOD | 1.25 | < LOD | < LOD | 5.37 | 4.13 | 75.3 | 0.443 | < LOD | 1.5 |
| < LOD | < LOD | 1.18 | < LOD | < LOD | 5.52 | 3.94 | 74.4 | < LOD | < LOD | 1.52 |
| < LOD | < LOD | 1.29 | < LOD | < LOD | 6.59 | 5.93 | 71.3 | < LOD | < LOD | 1.42 |
| < LOD | < LOD | 2.74 | < LOD | < LOD | 4.37 | 2.98 | 8.24 | < LOD | < LOD | 15.4 |
| < LOD | < LOD | 2.59 | < LOD | < LOD | 4.2 | 2.56 | 8.76 | < LOD | < LOD | 17 |
| < LOD | < LOD | 2.74 | < LOD | < LOD | 5.97 | 2.81 | 7.81 | < LOD | < LOD | 12.7 |
| < LOD | < LOD | 2.76 | < LOD | < LOD | 5.38 | 2.1 | 8.7 | < LOD | < LOD | 16.2 |
| < LOD | < LOD | 1.5 | < LOD | < LOD | 2.64 | 3.71 | 10.9 | < LOD | < LOD | 16.4 |
| < LOD | < LOD | 1.93 | < LOD | < LOD | 1.88 | 2.31 | 13.1 | < LOD | < LOD | 18.3 |
| < LOD | < LOD | 2.01 | < LOD | < LOD | 5.26 | 5 | 10.9 | < LOD | < LOD | 16.2 |
| < LOD | < LOD | 1.56 | < LOD | < LOD | 3.43 | 3.06 | 10.3 | < LOD | < LOD | 14.2 |
| < LOD | < LOD | 4.97 | < LOD | < LOD | 4.47 | 1.9 | 24.4 | < LOD | < LOD | 0.566 |
| < LOD | < LOD | 5.65 | < LOD | < LOD | 5.8 | 2.54 | 27.8 | < LOD | < LOD | 0.676 |
| < LOD | < LOD | 5.4 | < LOD | < LOD | 5.62 | 2.08 | 27.2 | < LOD | < LOD | 0.624 |
| < LOD | < LOD | 5.51 | < LOD | < LOD | 4.88 | 1.44 | 27.5 | < LOD | < LOD | 0.577 |
| < LOD | < LOD | 2.74 | < LOD | < LOD | 3.92 | 1.86 | 25.7 | < LOD | < LOD | 0.37 |
| < LOD | < LOD | 3.09 | < LOD | < LOD | 4.48 | 1.96 | 26.9 | < LOD | < LOD | 0.454 |
| < LOD | < LOD | 3.06 | < LOD | < LOD | 4.67 | 1.76 | 29.6 | < LOD | < LOD | 0.492 |
| < LOD | < LOD | 2.99 | < LOD | < LOD | 4.33 | 1.6 | 30.8 | < LOD | < LOD | 0.508 |
| < LOD | < LOD | 2.48 | < LOD | < LOD | 1.61 | 0.511 | 24.4 | < LOD | < LOD | 1.25 |
| < LOD | < LOD | 2.46 | < LOD | < LOD | 1.67 | 0.463 | 31.4 | < LOD | < LOD | 1.33 |
| < LOD | < LOD | 0.853 | < LOD | < LOD | 0.927 | 0.68 | 38.5 | < LOD | < LOD | 1.37 |
| < LOD | < LOD | 0.912 | < LOD | < LOD | 1.4 | 1.2 | 43.9 | < LOD | < LOD | 1.48 |
| < LOD | < LOD | < LOD | < LOD | < LOD | < LOD | < LOD | < LOD | < LOD | < LOD | < LOD |
| < LOD | < LOD | < LOD | < LOD | < LOD | < LOD | < LOD | < LOD | < LOD | < LOD | < LOD |

| Nitro-Tyr | PEA | Putrescine | SDMA | Serotonin | Spermidine | Spermine | Taurine | alpha-AAA | c4-OH-Pro | t4-OH-Pro |
|---|---|---|---|---|---|---|---|---|---|---|
| < LOD | < LOD | 733928.5714 | < LOD | < LOD | 505357.1429 | 136964.2857 | 11250000 | < LOD | < LOD | 312500 |
| < LOD | < LOD | 891071.4286 | < LOD | < LOD | 775000 | 244642.8571 | 12160714.29 | < LOD | < LOD | 364285.7143 |
| < LOD | < LOD | 861363.6364 | < LOD | < LOD | 1736363.636 | 486363.6364 | 12795454.55 | < LOD | < LOD | 379545.4545 |
| < LOD | < LOD | 856818.1818 | < LOD | < LOD | 1190909.091 | 431818.1818 | 13363636.36 | < LOD | < LOD | 354545.4545 |
| < LOD | < LOD | 360000 | < LOD | < LOD | 1554285.714 | 1011428.571 | 21714285.71 | < LOD | < LOD | 402857.1429 |
| < LOD | < LOD | 357142.8571 | < LOD | < LOD | 1534285.714 | 1180000 | 21514285.71 | 126571.4286 | < LOD | 428571.4286 |
| < LOD | < LOD | 262500 | < LOD | < LOD | 1133333.333 | 737500 | 15833333.33 | < LOD | < LOD | 293750 |
| < LOD | < LOD | 260416.6667 | < LOD | < LOD | 1118750 | 860416.6667 | 15687500 | 92291.66667 | < LOD | 312500 |
| < LOD | < LOD | 856250 | < LOD | < LOD | 1365625 | 931250 | 2575000 | < LOD | < LOD | 4812500 |
| < LOD | < LOD | 809375 | < LOD | < LOD | 1312500 | 800000 | 2737500 | < LOD | < LOD | 5312500 |
| < LOD | < LOD | 944827.5862 | < LOD | < LOD | 2058620.69 | 968965.5172 | 2693103.448 | < LOD | < LOD | 4379310.345 |
| < LOD | < LOD | 951724.1379 | < LOD | < LOD | 1855172.414 | 724137.931 | 3000000 | < LOD | < LOD | 5586206.897 |
| < LOD | < LOD | 300000 | < LOD | < LOD | 528000 | 742000 | 2180000 | < LOD | < LOD | 3280000 |
| < LOD | < LOD | 386000 | < LOD | < LOD | 376000 | 462000 | 2620000 | < LOD | < LOD | 3660000 |
| < LOD | < LOD | 456818.1818 | < LOD | < LOD | 1195454.545 | 1136363.636 | 2477272.727 | < LOD | < LOD | 3681818.182 |
| < LOD | < LOD | 354545.4545 | < LOD | < LOD | 779545.4545 | 695454.5455 | 2340909.091 | < LOD | < LOD | 3227272.727 |
| < LOD | < LOD | 1911538.462 | < LOD | < LOD | 1719230.769 | 730769.2308 | 9384615.385 | < LOD | < LOD | 217692.3077 |
| < LOD | < LOD | 2173076.923 | < LOD | < LOD | 2230769.231 | 976923.0769 | 10692307.69 | < LOD | < LOD | 260000 |
| < LOD | < LOD | 1636363.636 | < LOD | < LOD | 1703030.303 | 630303.0303 | 8242424.242 | < LOD | < LOD | 189090.9091 |
| < LOD | < LOD | 1669696.97 | < LOD | < LOD | 1478787.879 | 436363.6364 | 8333333.333 | < LOD | < LOD | 174848.4848 |
| < LOD | < LOD | 1442105.263 | < LOD | < LOD | 2063157.895 | 978947.3684 | 13526315.79 | < LOD | < LOD | 194736.8421 |
| < LOD | < LOD | 1626315.789 | < LOD | < LOD | 2357894.737 | 1031578.947 | 14157894.74 | < LOD | < LOD | 238947.3684 |
| < LOD | < LOD | 1700000 | < LOD | < LOD | 2594444.444 | 977777.7778 | 16444444.44 | < LOD | < LOD | 273333.3333 |
| < LOD | < LOD | 1661111.111 | < LOD | < LOD | 2405555.556 | 888888.8889 | 17111111.11 | < LOD | < LOD | 282222.2222 |
| < LOD | < LOD | 476923.0769 | < LOD | < LOD | 309615.3846 | 98269.23077 | 4692307.692 | < LOD | < LOD | 240384.6154 |
| < LOD | < LOD | 473076.9231 | < LOD | < LOD | 321153.8462 | 89038.46154 | 6038461.538 | < LOD | < LOD | 255769.2308 |
| < LOD | < LOD | 110779.2208 | < LOD | < LOD | 120389.6104 | 88311.68831 | 5000000 | < LOD | < LOD | 177922.0779 |
| < LOD | < LOD | 118441.5584 | < LOD | < LOD | 181818.1818 | 155844.1558 | 5701298.701 | < LOD | < LOD | 192207.7922 |

| Nitro-Tyr | PEA | Putrescine | SDMA | Serotonin | Spermidine | Spermine | Taurine | alpha-AAA | c4-OH-Pro | t4-OH-Pro |
|---|---|---|---|---|---|---|---|---|---|---|
| | | 2.27654E-05 | | | 0.372782947 | 0.002557008 | 0.011415023 | | | 0.858949698 |
| | | 3.68842E-05 | | | 0.011085954 | 0.544030842 | 0.037084055 | | | 0.00180891 |
| | | 0.130129115 | | | 0.025280163 | 0.055487821 | 0.001040367 | | | 0.225601714 |
| | | 0.002299597 | | | 0.141115087 | 0.130727779 | 0.000904252 | | | 0.008997136 |
| | | 0.009748044 | | | 0.002894346 | 0.004705355 | 0.006338794 | | | 0.025044127 |

Supplementary Table 7. Concentration of metabolites used in for metabolic flux analysis. Concentrations are reported as μM/500k cells (Average of time points 0, 10, 30 and 90min).

| | **DU145** | | | **DU145-RR** | | | **LNCaP** | | | **LNCaP-RR** | | |
|---|---|---|---|---|---|---|---|---|---|---|---|---|
| | **#1** | **#2** | **#3** | **#1** | **#2** | **#3** | **#1** | **#2** | **#3** | **#1** | **#2** | **#3** |
| **Citrate** | 7.77 | 7.97 | 11.35 | 9.67 | 3.81 | 21.32 | 11.61 | 9.14 | 9.42 | 117.23 | 19.34 | 54.70 |
| **Malate** | 33.23 | 27.87 | 37.77 | 26.69 | 17.62 | 110.53 | 28.15 | 27.49 | 31.27 | 100.73 | 27.07 | 38.34 |
| **Aspartate** | 221.72 | 175.25 | 310.34 | 190.91 | 119.66 | 933.39 | 172.03 | 108.65 | 159.03 | 1560.42 | 331.40 | 829.06 |
| **Glutamate** | 993.99 | 1003.60 | 1352.51 | 932.34 | 596.93 | 4112.85 | 405.97 | 355.22 | 527.94 | 2756.83 | 705.82 | 1336.07 |
| **Succinate** | 15.72 | 9.64 | 15.78 | 39.01 | 10.51 | 66.35 | 21.48 | 9.20 | 9.71 | 430.59 | 76.31 | 251.79 |

Supplementary Figure 1

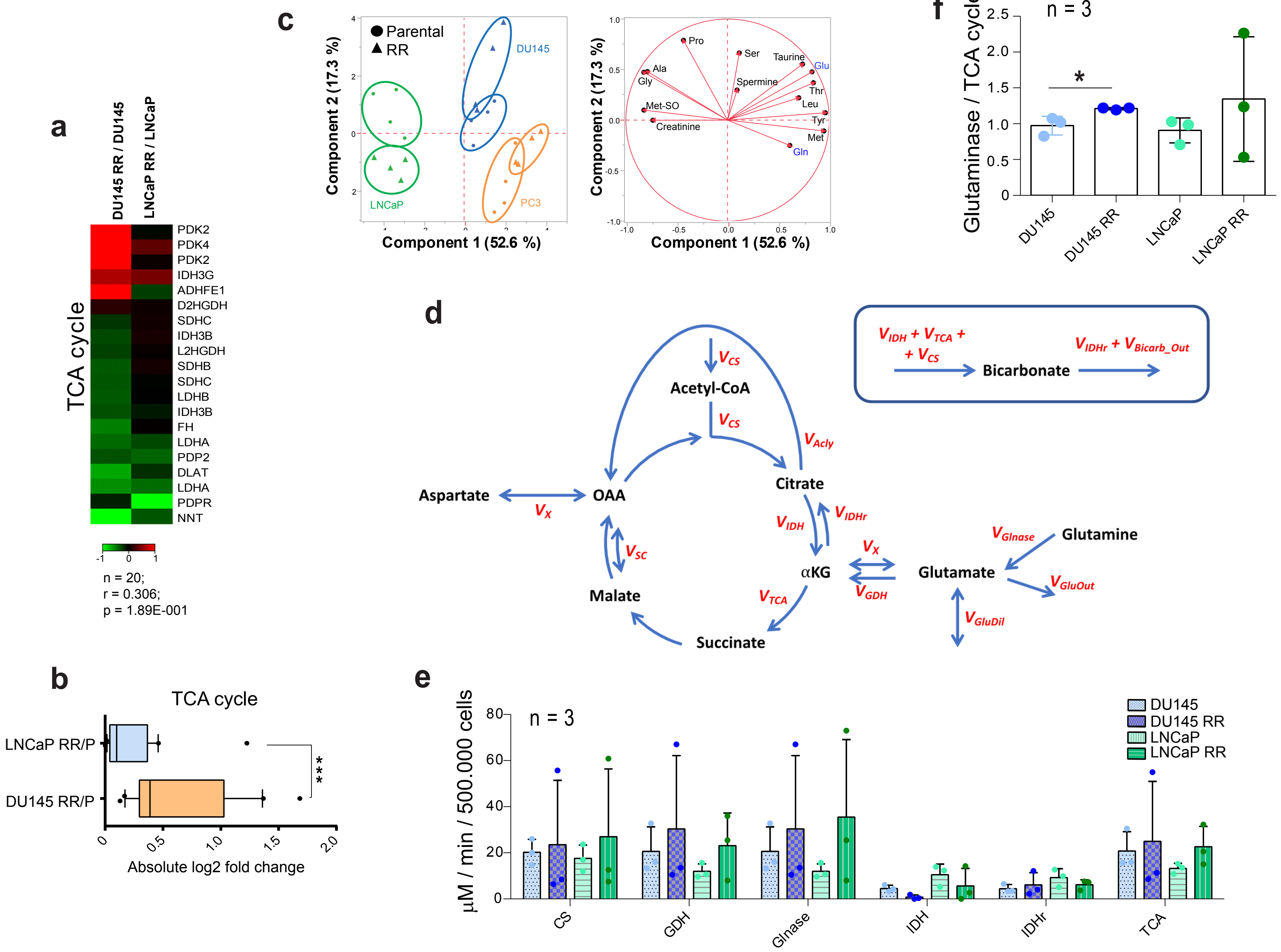

Supplementary Figure 2

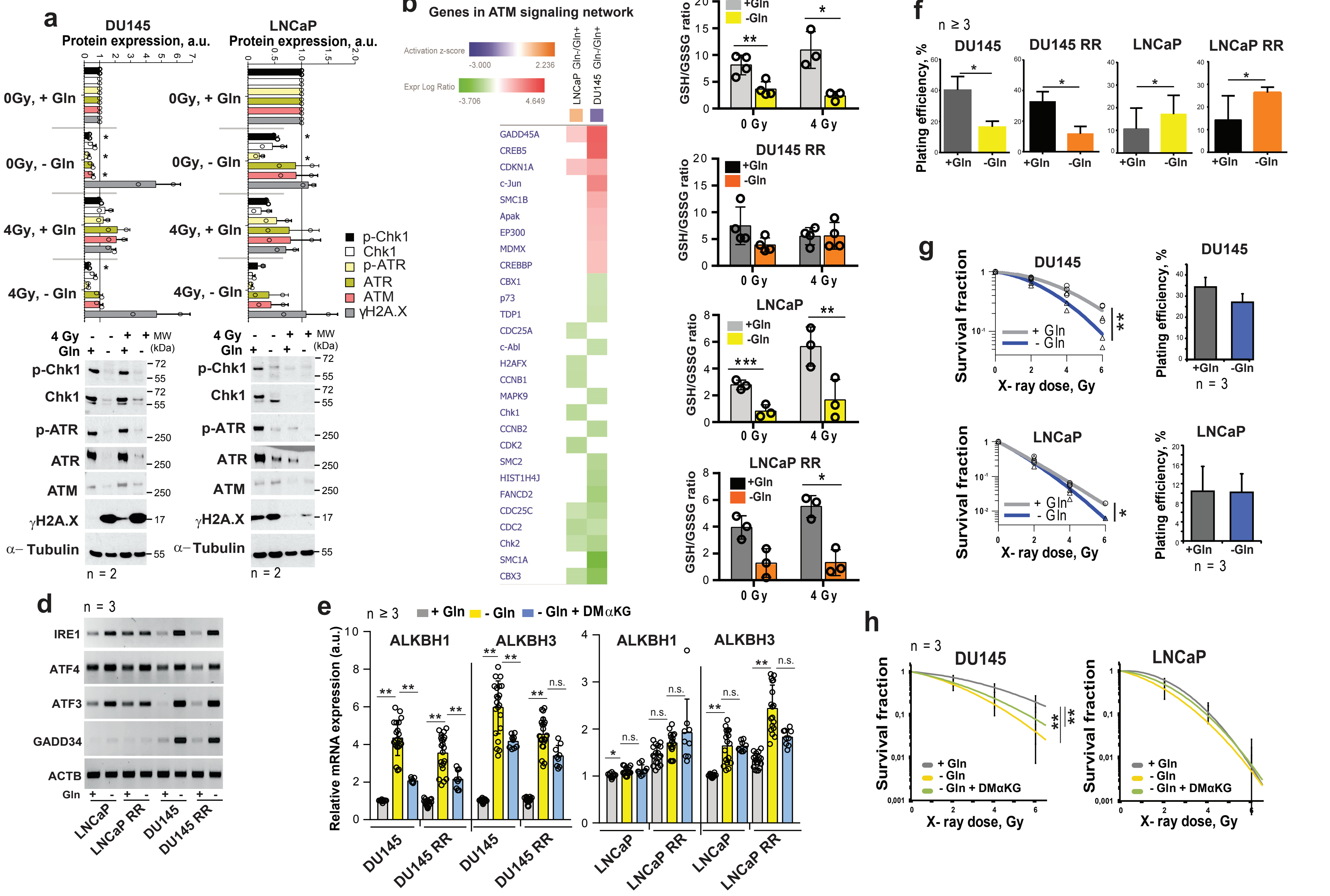

Supplementary Figure 3

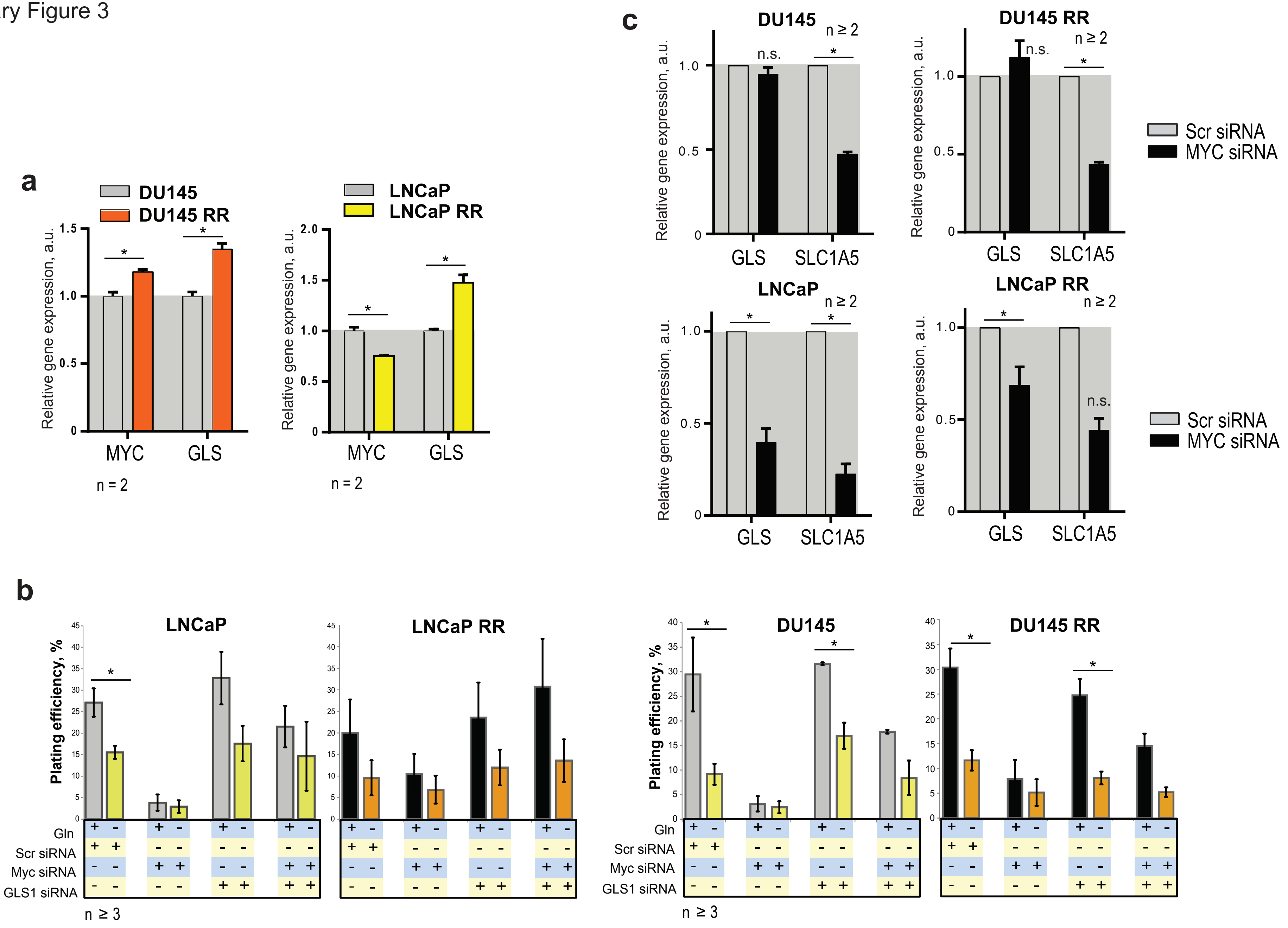

Supplementary Figure 4

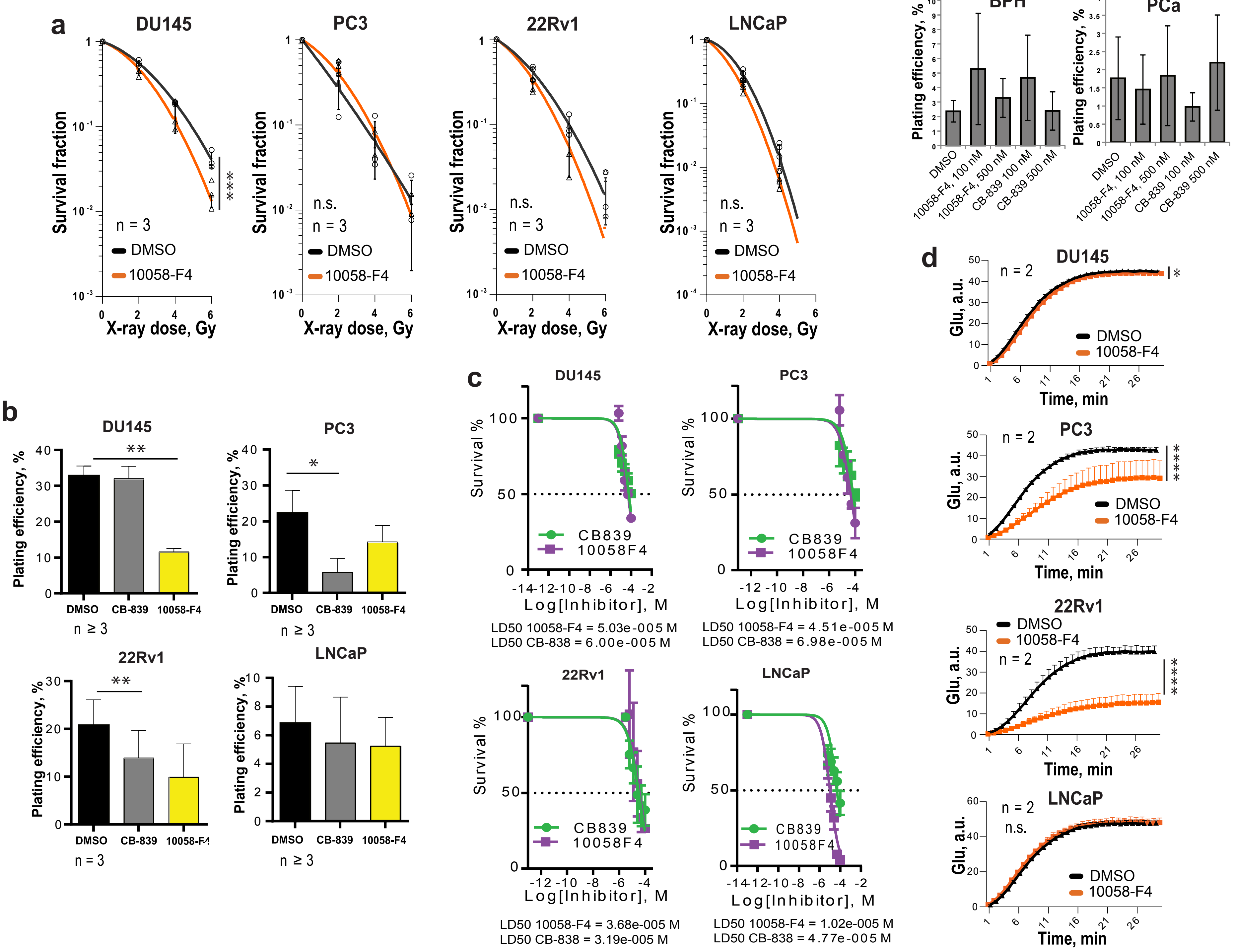

Supplementary Figure 5

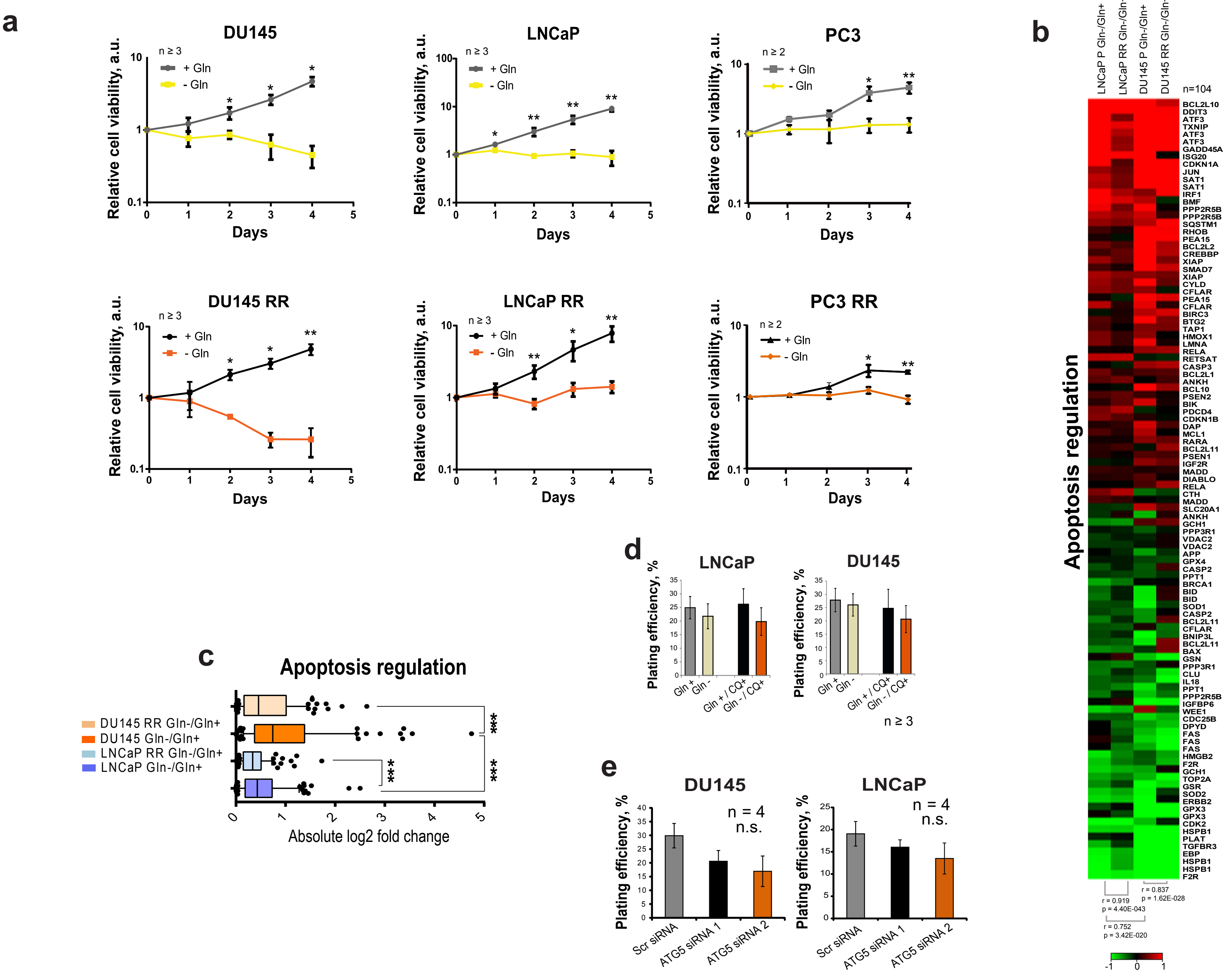

Supplementary Figure 6

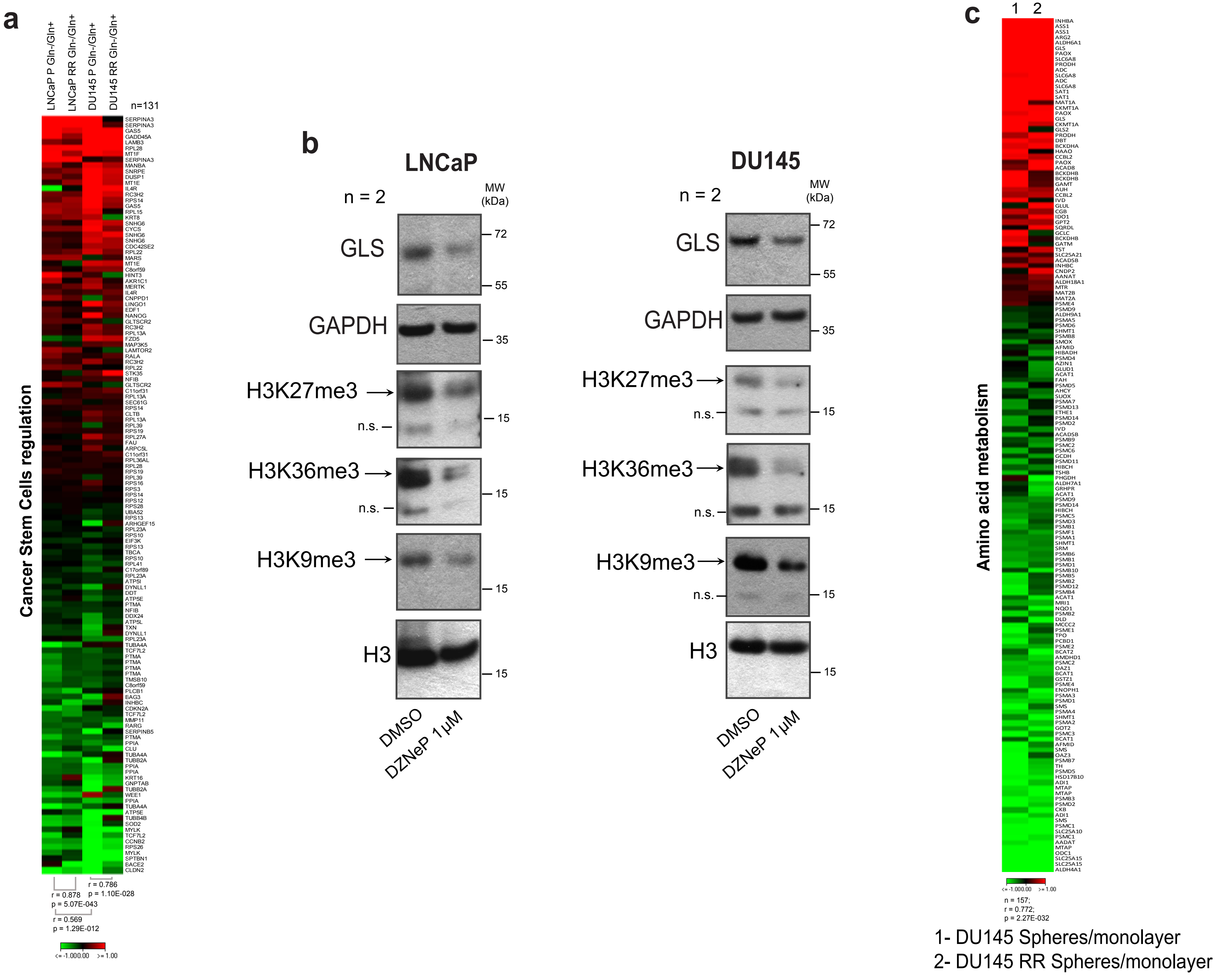

Supplementary Figure 7

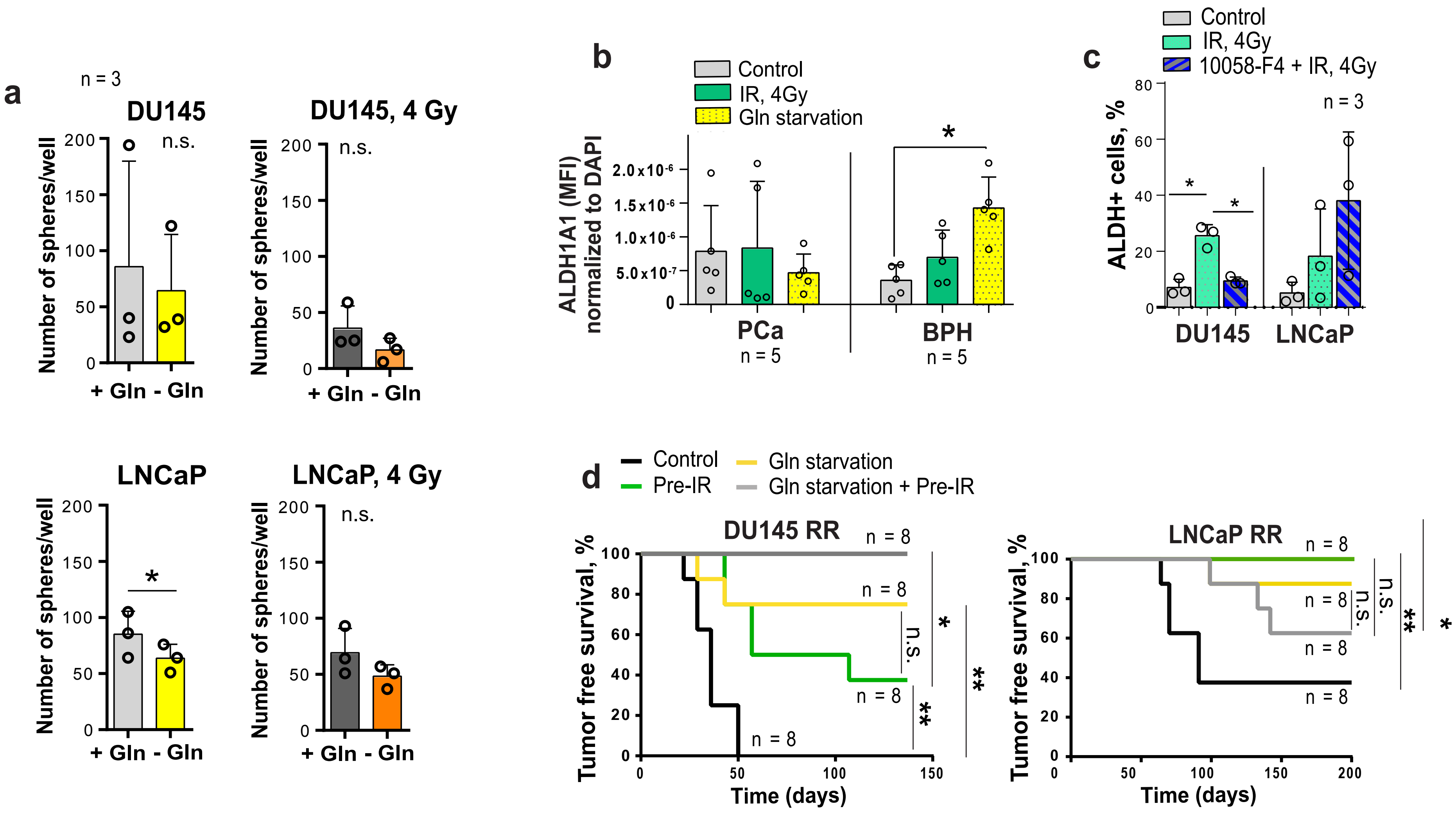

Supplementary Figure 8

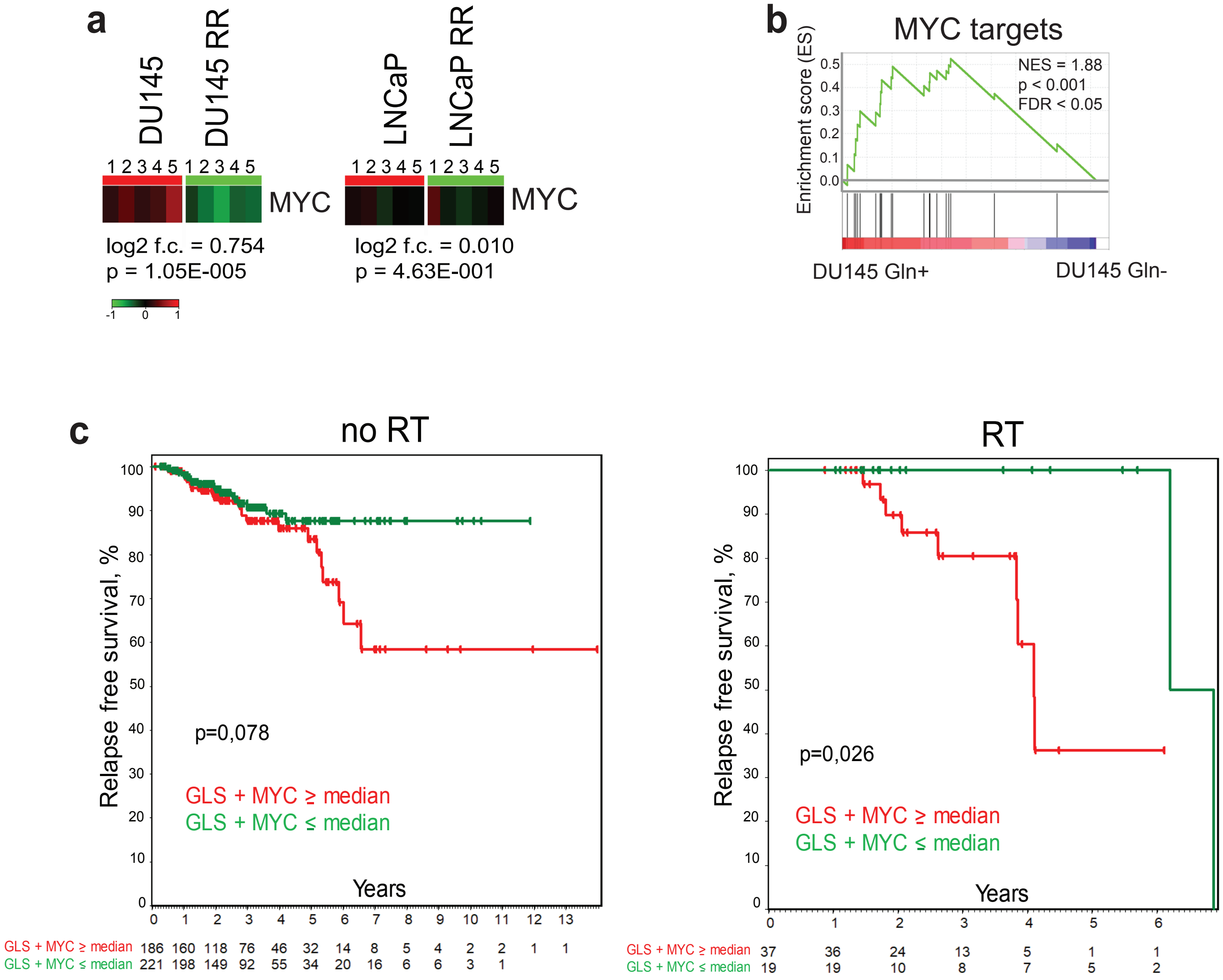
